# Comparative study on chromatin loop callers using Hi-C data reveals their effectiveness

**DOI:** 10.1101/2023.11.24.567971

**Authors:** H. M. A. Mohit Chowdhury, Terrance Boult, Oluwatosin Oluwadare

## Abstract

The chromosome is a fundamental component of cell biology, housing DNA that encapsulates hierarchical genetic information. DNA compresses its size by forming loops, and these loop regions contain numerous protein particles, including CTCF, SMC3, H3 histone, and Topologically Associating Domains (TADs). In this study, we conducted a comprehensive study of 22 loop calling methods. Additionally, we have provided detailed insights into the methodologies underlying these algorithms for loop detection, categorizing them into five distinct groups based on their fundamental approaches. Furthermore, we have included critical information such as resolution, input and output formats, and parameters. For this analysis, we utilized the primary and replicate GM12878 Hi-C datasets at 5KB and 10KB resolutions. Our evaluation criteria encompassed various factors, including loop count, reproducibility, overlap, running time, Aggregated Peak Analysis (APA), and recovery of protein-specific sites such as CTCF, H3K27ac, and RNAPII. This analysis offers insights into the loop detection processes of each method, along with the strengths and weaknesses of each, enabling readers to effectively choose suitable methods for their datasets. We evaluate the capabilities of these tools and introduce a novel Biological, Consistency, and Computational robustness score (*BCC*_*score*_) to measure their overall robustness ensuring a comprehensive evaluation of their performance.

## 1 Introduction

DNA and chromosomes hold the most important information about a species. Scientists have been working to reveal the internal structure of chromosomes and DNA to answer questions about intra-chromosomal interaction, hierarchical properties, and DNA segments^1,2^. Regulatory information is also important to solve real-life problems such as disease prediction and analysis^3^. Studies have revealed that each chromosome is positioned in a specific region known as a *chromosome territory*^4^, characterized by a specific pattern. Inside chromatin (Figure 1), a ring-shaped cohesin protein pulls DNA through the center of it to create a loop and is bounded by CTCF (called extrusion barrier)^5,6^. This loop results in the 3D structure of DNA in a small region inside the chromatin. Peaks are areas enriched in aligned reads due to protein binding from ChiP-sequencing or MDIP-sequencing^7,8^. These loops and peaks are important regions from which we can answer questions about gene interaction, structure, and protein structure^6,9^.Various proteins have been found in these regions, such as cohesin, CTCF, and some H3 protein markers like H3K27ac and H3K27me3^5,10^. Scientists have also observed that TADs around these loop regions are crucial for chromosome interaction^2,11^.

**Figure 1.**
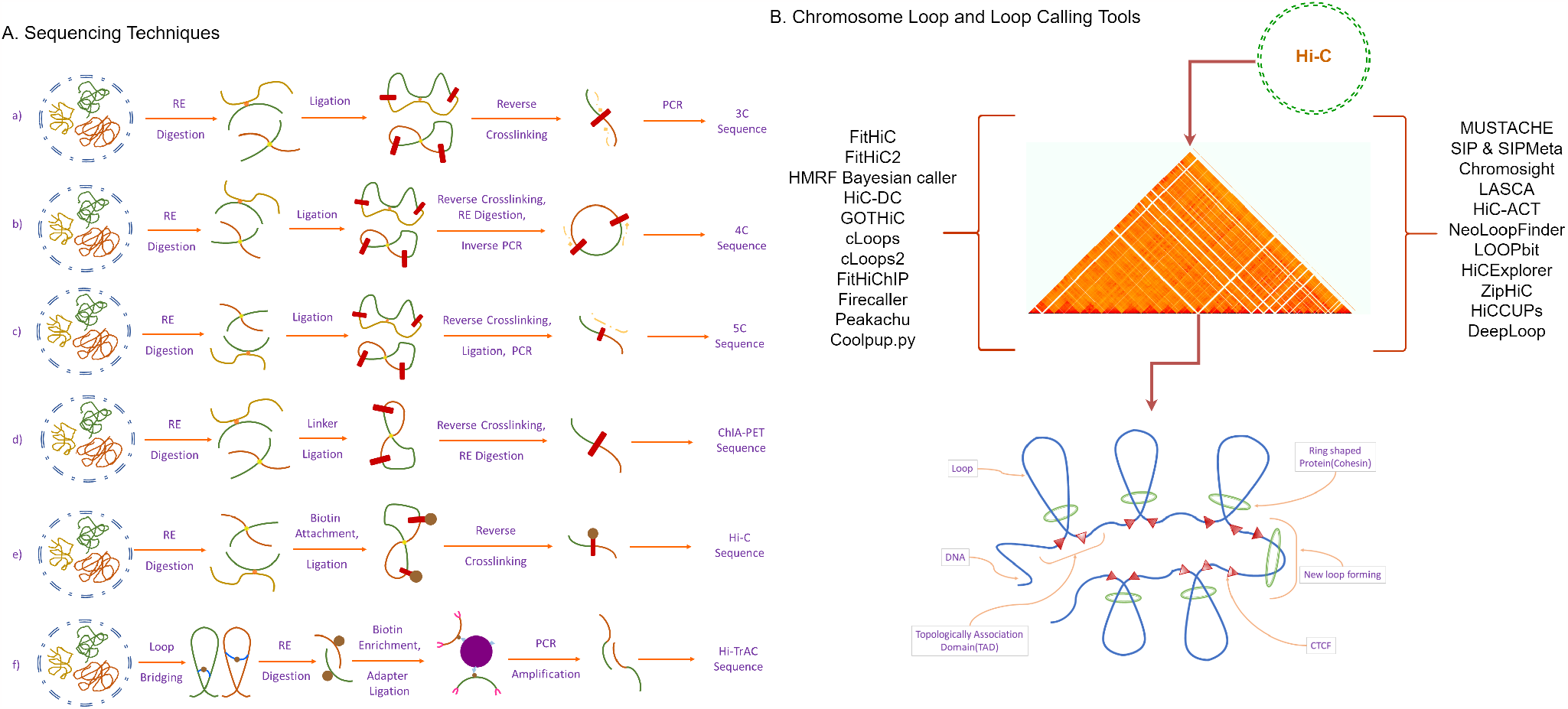
A brief overview of sequencing techniques and chromatin loops. **(A)** Chromosome conformation capture technologies: (a) Chromosome spatial organization analysis in 3C technology starts with cell population crosslinking and fragmentation with a restriction enzyme. Next, it goes through intramolecular ligation and reverse-crosslinking and performs semi-quantitative or quantitative PCR. (b) Chromosome spatial organization analysis in 4C technology starts with cell population crosslinking and fragmentation with restriction enzyme. Next, it goes through intramolecular ligation and reverse crosslinking. Next, it goes through digestion with a restriction enzyme and ligation, and finally, applies inverse PCR. (c) Chromosome spatial organization analysis in 5C technology starts with cell population crosslinking and fragmentation with a restriction enzyme. Next, it goes through intramolecular ligation and reverse-crosslinking and performs synthetic ligation and multiplex PCR. (d) Chromosome spatial organization analysis in ChIA-PET technology starts with cell population crosslinking and fragmentation with a restriction enzyme. Then, DNA linker ligation attracts nucleotides and performs reverse crosslinking and PCR. (e) Chromosome spatial organization analysis in Hi-C technology starts with cell population crosslinking and fragmentation with restriction enzyme. Then, it attaches biotin-labeled nucleotide, and goes through blunt ligation and PCR^1,2,10,15,64–66^. (f) First, HiC-TrAC creates a bridge on chromatin loops and splices DNA with restriction enzymes. Then, the process is fertilized with streptavidin beads. Finally, DNA fragments having a biotin label attach with a multiplexing adapter and go through a PCR amplification^9^. **(B)** Chromatin TADs are the region where chromatin interaction occurs and the starting region of a chromatin loop. The green-colored ring-shaped protein first pulls DNA through it creating a loop. CTCF as a binder or lock for this ring and widely known as CCCTC binding factor (transcription factor) or 11 zinc finger protein. It can function in many ways such as blocking communication, insulator protein, or transcriptional activator^5^. TADs^11^ are formed by the folding of chromatin, which is a complex of DNA, RNA, and proteins. TADs represent regions of the genome where DNA sequences are more likely to interact with each other spatially. The Ring-shaped protein, which tightens the loops is called the cohesin^67^.

The evolution in C-technology was initiated by Dekker et al. when they expanded the 3C method^1^. Subsequently, other 3C-based methods (Hi-C, ChIA-PET^12,13^, Hi-TrAC^14^) were developed sharing some common methodology briefly presented in Figure 1. Hi-C, a combination of 3C and next-generation sequencing techniques, represents a significant advance in genome analysis. One of its main advantages is that it is not subject to a set of any primers^15–17^. It is an unbiased and unsupervised method^2,18^ for genome analysis, generating genome-wide contact maps^15^. It is widely used for analyzing genomic organizational principles, chromosome structure at the mitotic stage, and anatomical changes in human disease^2,19–21^. The advent of Chromatin Conformation Capture (3C) technology^15^ and Hi-C technology^16^ has propelled gene analysis in various directions and has influenced the development of numerous loop and peak calling techniques^22–41^. Although their primary aim is to identify loops and peaks, these methods offer a secondary advantage by providing information about gene regulation, such as interaction, structure, and protein reactions^14,29,41^. The development of machine learning algorithms has propelled 3D genome spatial architecture analyses into a new dimension^42–46^. Specifically in the loop detection domain, Scientists have developed different tools to predict loop regions, employing various machine learning-based approaches such as computer vision and classification based methods. Mustache^39^, Chromosight^41^, SIP & SIPMeta^40^ have demonstrated the application of computer vision algorithms to predict loop regions, marking a new era in genomic analysis with many other tools.

In this manuscript, we present a comprehensive analysis of 22 loop detection tools based on Hi-C datasets. We evaluate how these tools perform in predicting loops, their recovery of biological features such as H3K27ac, RNAPII, and CTCF, and discuss their strengths and weaknesses. Our analysis goes beyond theory, giving practical insights into how these tools can be used, including the necessary technical details and parameters. By merging these aspects, we identify overlaps, uncovering connections, computational efficiency, similarities and results consistency in the studied techniques. To qualitatively measure the capabilities of these tools across these analysis categories, we created a novel aggregated score called the *BCC*_*score*_ to measure their overall robustness ensuring a comprehensive evaluation of their performance.

## 2 Results

We used GM12878^47^ (Human Lymphoblastoid) primary, replicate and Knight-Ruiz (KR) normalized Hi-C datasets for our analysis. We used chromosomes 1 and 6 at 5KB and 10KB resolution for our analysis. We prepared input data using HiCExplorer (cool), and sam and bam (bed and bedpe) tools. All methods were analyzed with their input and output details, and we compared their loop count across different resolutions. FitHiC2 loop count is represented with a factor of 1/10 to align with other loop counts. cLoops and cLoops2 are represented for chromosome 1 and 6 without any specific resolution as both algorithms input data format requirement (.bedpe) could not be normalized. For robustness assessments, we assessed their overlap, APA score, recovery (CTCF, H3K27ac, and RNAPII) and running time in the following sections.

### 2.1 Loop detection within different resolutions and normalization

We successfully executed 11 out of 22 methods in our analysis, assessing their loop prediction capabilities (Figure 2). The remaining 11 methods couldn’t be executed due to computational issues, with some failing to produce results or encountering errors during execution: ZipHiC^33^ does not provide a clear instruction to run their script, HMRF Bayesian caller^35^ has no public source code repository, LOOPBit^22^, Coolpup.py^38^, DeepLoop^48^ and FIREcaller^36^ errored out during analysis with no results, HiC-ACT^27^ and GOTHiC^31^ did not produce any output upon execution, and we couldn’t access a R library for HiC-DC^32^ or access its installation instruction.

**Figure 2.**
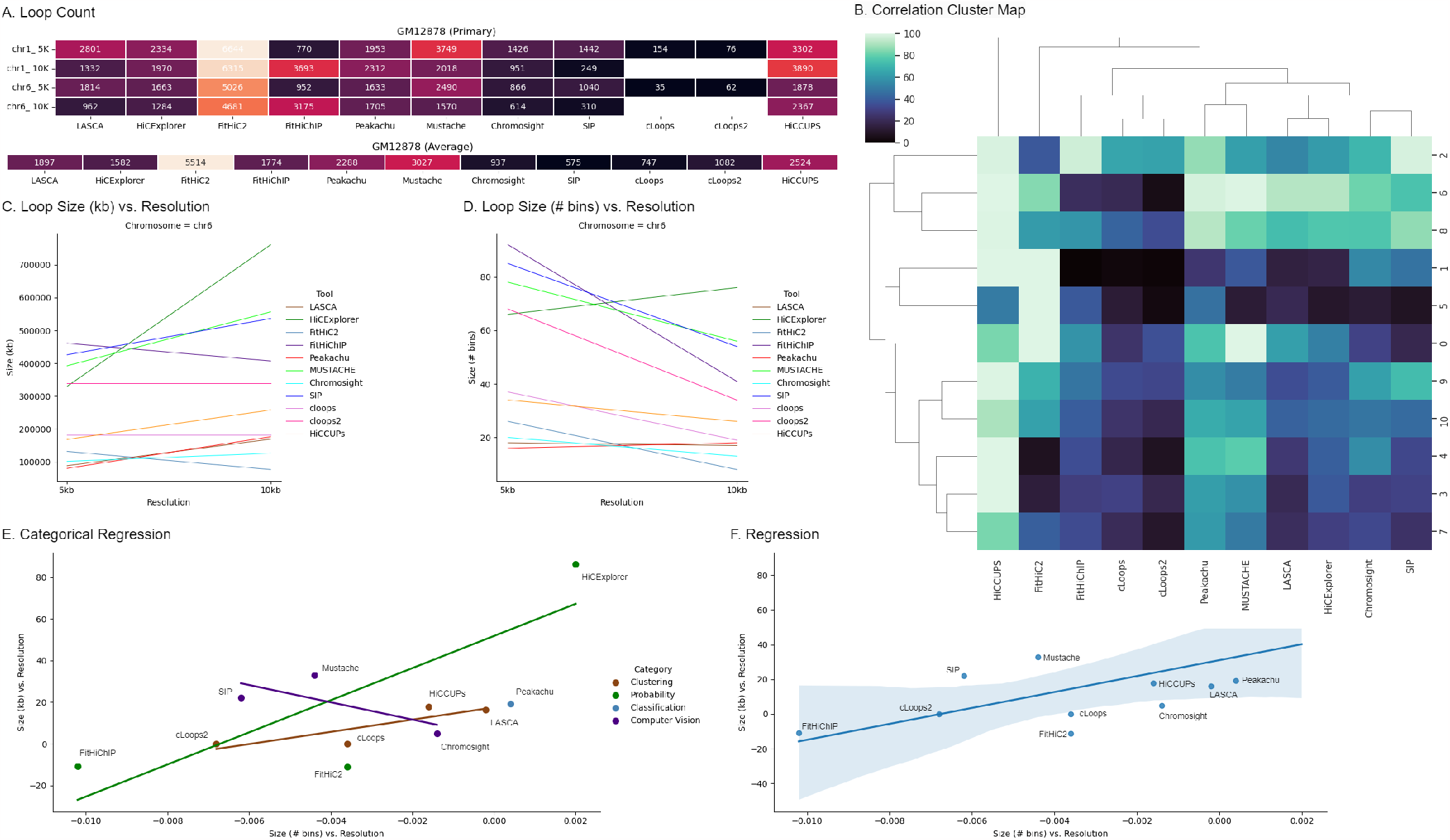
Identification of chromatin loops at 5KB and 10KB using chromosome 1 and 6 primary GM12878 Hi-C dataset. A. Chromatin loop count across chromosome 1 and 6 primary data at 5KB and 10KB. B. Clustering of chromatin loop calling tools according to the loop prediction. C. Representation of chromatin mean loop size at 5KB and 10KB resolution. D. Illustration of chromatin loop size in terms of number of mean bin sizes at 5KB and 10KB resolution. Regression plots of chromatin loop caller tools calculating the slope of resolution and loop size (kb) vs. loop size (# bins) and resolution where E represents categorical regression plot and F represents overall regression using chromosome 6 at 5KB and 10KB.

In Figure 2A, our analysis shows that while FitHiC2 predicts most chromatin interactions, we represented it with a factor of 1/10 to align with other tools. At 5KB resolution for chromosomes 1 and 6, Mustache detects the majority of loops. At 10KB resolution for chromosomes 1 and 6, HiCCUPS detects the most, whereas HiCExplorer identifies around half of what HiCCUPS does. In both cases, cLoops and cLoops2 detect the fewest loops, and they are not directly associated with a specific resolution but receive a range of resolutions. LASCA performs better at the 5KB resolution, almost comparable to HiCCUPS. cLoops and cLoops2 are represented here for chromosomes 1 and 6 without resolution, receiving bedpe input format; it wasn’t possible to use normalized data, as they lack an option for normalization. In terms of average loop counts, Mustache outperforms other tools. Overall, cLoops detects the fewest loops, and HiCCUPS and LASCA identify loops comparable to Mustache. In the cluster map (Figure 2B), FitHiC2 formed a separate cluster due to its prediction of a large number of contacts. HiCCUPS and Mustache shared a cluster with a high number of loops. cLoops and cLoops2 were in the same cluster with the lowest number of loops, while Chromosight and SIP formed another cluster. LASCA and HiCExplorer created yet another cluster, and Peakachu formed a cluster with Mustache and the HiCCUPS cluster. This cluster map highlighted hierarchical correlations among the tools in terms of loop count.

We compared loop sizes (Figure 2C and 2D) at 5KB and 10KB resolution, revealing a similar trend across different chromosomes. With the exception of FitHiC2 (chromosomes 1 and 6) and FitHiChIP (chromosome 6), all loop caller loop sizes (KB) increased with the resolution. Conversely, the average number of bins in a loop demonstrated an opposite trend, decreasing with the resolution increase. Only HiCExplorer and Peakachu exhibited an increase in loop size (# of bins) with the resolution increase. We calculated the slope of loop size and resolution, noting negative slopes for FitHiC2 and FitHiChIP, while all other tools showed positive slopes. Additionally, the slope of loop size (# bins) and resolution indicated negative slopes for LASCA, HiCExplorer, FitHiC2, Mustache, Chromosights, and SIP. A linear regression plot (Figure 2E and 2F) categorized tools and demonstrated that FitHiChIP, cLoops2, HiCCUPS, LASCA, Peakachu fell within the regression boundary for both chromosomes. HiCCUPS, LASCA, and Peakachu consistently maintained the same trends. The regression category-wise plot further elucidated individual category information. Supplemental Figure 1 shows the chromatin loop counts for GM12878 replicate and KR normalized dataset. Further individual analyses of each tool are presented in the subsequent paragraphs, comparing results related to loop counts and input parameter robustness.

LASCA^23^ implements Weibull distribution mechanism for loop detection and enhancer-promoter interaction using Hi-C data across different types of organisms. Though they do not provide any command line facility to run, we can use LASCA to identify loops at a specific chromosome at a specific resolution importing LASCA as a Python library. While analyzing LASCA, we counted 1332 and 2801 loops in chromosome 1 at 10KB and 5KB resolution using primary GM12878 cell line (Supplemental Table 5). LASCA performs better at 5KB resolution compared to 10KB resolution across primary, replicate, and normalized data (Supplemental Table 5, 6, 7).

HiCExplorer provides a robust toolset (such as normalization, data conversion, loop prediction) for chromosomal data analysis and performs well with high-resolution data. HiCExplorer provides an option to set user-specific p-value and threads, threads per chromosome. We used the default setting and got 1582 loop count at 5KB and 10KB resolution using chromosome 1 and 6 (GM12878) (Supplemental Table 5, 6, 7, 8). Though it does not provide any parameter for specific resolution, we extracted 5KB and 10KB chromosome-specific data using hicConvertFormat. hicConvertFormat also supports normalization (cool to cool format only). HiCExplorer also detects protein binding sites that correlate with detected loops and they used different types of dataset for their analysis such as ChIA-PET, HiChIP along with Hi-C.

FitHiC mainly identifies mid-range intra-chromosomal contacts considering the looping effect and biases and finds high-confidence contacts in insulator and heterochromatin regions. FitHiC2 is an updated version of FitHiC where they minimized the mid-range intra-chromosomal contact analysis limitation. They introduced genome-wide contacts analysis in high resolution without sacrificing significant loops. FitHiC2 can analyze data at a specific resolution and has an option to specify the intra-chromosome or inter-chromosome analysis. It requires input files from other analysis tools such as HiCPro and they provided all the scripts for getting these inputs. FitHiC2 produces outputs of significant interaction contact and indirectly we can infer loops at that region. Here we used FitHiC2 in our analysis and it produces 1324353 contacts for GM12878 chromosome 1 primary data at 5KB resolution. We filtered out FitHiC2 reported contact count considering 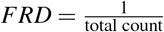as they suggested in their manuscript and got 55140 average loop count from chromosome 1 and 6 (Supplemental Table 8). While running their repository, we encountered a Python error which is also fixed in our fork repository and we uploaded a docker image for further analysis.

FitHiChIP^30^ is mainly focused on HiChIP/PLAC-seq data where they analyzed non-uniform coverage by scaling contact counts which ultimately produces loops even at 2.5KB resolution. This tool is a versatile tool providing differential loop analysis option. During our analysis, FitHiChIP produces 27220 contact loops from primary GM12878 chromosome 1 at 5KB resolution and we used 1*e*^−6^ threshold of p-value as they suggested (Supplemental Table 5). FitHiChIP accepts HiCPro valid pair files, bin interval and contact matrix, bed, cool, and hic formatted files. It requires a configuration file where we can pass all the settings. In our analysis, we used chromosome-wise cool files and considered peak-to-all interaction analysis using coverage bias correction setting. Though it does not support chromosome-wise analysis, it has a parameter for passing the bin size where user can specify their intended resolution in full form.

Peakachu^37^ is a Random Forest classifier and provides pre-build models for different combinations of intra reads and high confidence for different types of datasets such as Hi-C, Micro-C, HiChiP, etc. Though they provided an option to train a model, we used their pre-build model. They also accept specific chromosome numbers and resolutions which facilitate the user to analyze as needed. Though it provides balancing parameter for using ICE or KR matrix, it did not accept any specific parameter. In our analysis, we used KR normalization data from HiCExplorer along with primary data. From our primary data analysis, we got 1338705 (Supplemental Table 5) interactions from chromosome 1 at 5KB resolution using the 70% model. In addition to specific chromosome analysis, they have the option to analyze the whole genome. Peakachu can recover most of the loops from protein-centric datasets such as ChIP and ChIA-PET, and they also showed short-range interaction recovery in their analysis result.

MUSTACHE^39^ utilized the scale-space theory of computer vision to detect chromatin loops at different sequencing depths of Hi-C and Micro-C data. Mustache provides normalization techniques for users for hic and cool files and bias files for text-based contact map files along with process, thread,threshold (p-value), resolution, and chromosome-wise analysis. Mustache detects 3027 loops on average at 5KB and 10KB resolutions in our analysis from chromosomes 1 and 6 using GM12878 cells (Supplemental Table 8). Mustache can analyze chromosomes at 1KB resolution Micro-C and 5KB resolution Hi-C data.

Chromosight^41^ implemented pattern recognition technique to detect loops. From our analysis, Chromosight detects 937 at 10KB and 5KB resolutions from chromosomes 1 and 6 (Supplemental Table 8), and it performs better at 5KB resolution compared to 10KB resolution (Supplemental Table 5, 6, 7). It can identify borders, centromeres, etc and accepts thread parameter. Chromosight analyzes the whole genome and it does not have any parameter for specific resolution. We provided a specific chromosome contact map at a specific resolution in our analysis. It provides three normalizations (auto, raw, and forced) from the user and has inter chromosomal analysis option.

SIP^40^ developed to identify missing loops from previous loop callers considering the noise and sequencing depth. SIP can detect more loops at 5KB resolution compared to 10KB resolution. Overall, SIP identified 575 from chromosomes 1 and 6 in GM12878 cell (Supplemental Table 8). SIP provides UI for users flexibility. It accepts resolution, CPUs, normalization (VC, VC_SQRT, and KR), FDR, and threshold value parameters. In our analysis, we used cool files, but it support hic and processed files as input. It can analyze deeply sequenced genomes even at 1KB resolution cLoops^24^ and cLoops2^25^ are DBSCAN based loop detection algorithm. cLoops calculates the distance between two neighbors describing the distance between two neighbors and analyzes pair-end tags to identify loops with *O*(*Nlog*(*N*)) running time in addition to parallel computation. They provides different analysis plot scripts (heatmap, data quality plots) and chromosome-wise analysis. cLoops2 is the updated version of cLoops with an optimized DBSCAN clustering algorithm with running time *O*(*N*) and provides loop and peak calling algorithm in different ways along with differential loop and domain calling. cLoops2 was developed for detecting loops on Hi-TrAC/TrAC looping data. It can still be used for loop detection for ChIA-PET and HiChIP data like cLoops. cLoops2 provides chromosome-specific analysis but we cannot provide specific resolution to it. Though their loop detection is comparably close to other methods, both, cLoops and cLoops2, produce a poor loop count (747, 1082) in our analysis (Supplemental Table 8). Compared to cLoops, cLoops2 had more loop counts (Supplemental Table 5, 6, 7). Like cLoops, cLoops2 provides chromosome-wise genomic analysis regardless of any resolution. cLoops2 has analysis scripts such as aggregated peaks, domains, etc. cLoops and cLoops2 do not provide any normalization parameter in their command line. Users can specify the number of CPUs to be used in cLoops2 and it has a data conversion tool to other formats.

HiCCUPS provides different versions to run on CPU and GPU. They provide in-tool normalization techniques along with a specific chromosome and resolution. Users can specify thread, threshold value (FDR), and merging distance. During our analysis, we used default parameters, and on average, it predicts 2524 loops (Supplemental Table 8). It predicts the maximum number of loops at 10KB resolution rather than 5KB resolution (Supplemental Table 5, 6, 7). Though FitHiC2 predicts most of the contacts, it predicts the second maximum number of loops in our analysis compared with Mustache.

### 2.2 Overlap and Reproducibility

For empirical analysis, we executed all the methods using the GM12878 dataset to observe the overlap (Figure 3-left). We divided our dataset into various combinations, involving chromosomes 1 and 6 at 5KB and 10KB resolution. Notably, cLoops and cLoops2 accept bedpe files and do not allow any normalization parameter, leading our analysis without normalization data. For the 5KB data on chromosome 6, FitHiC2 exhibited a 95% overlap among primary, replicate, and KR normalized data, indicating high reproducibility. In contrast, cLoops, cLoops2, and FitHiChIP displayed the lowest overlap, nearly 0%. On average, Chromosight, HiCCUPS, and SIP exhibited 25% - 46% overlap. Chromosight showed more reproducibility between primary and replicate datasets, while SIP displayed more reproducibility between normalized and replicate data. Chromosight, FitHiC2, and HiCCUPS demonstrated higher reproducibility rates across our dataset combinations.

**Figure 3.**
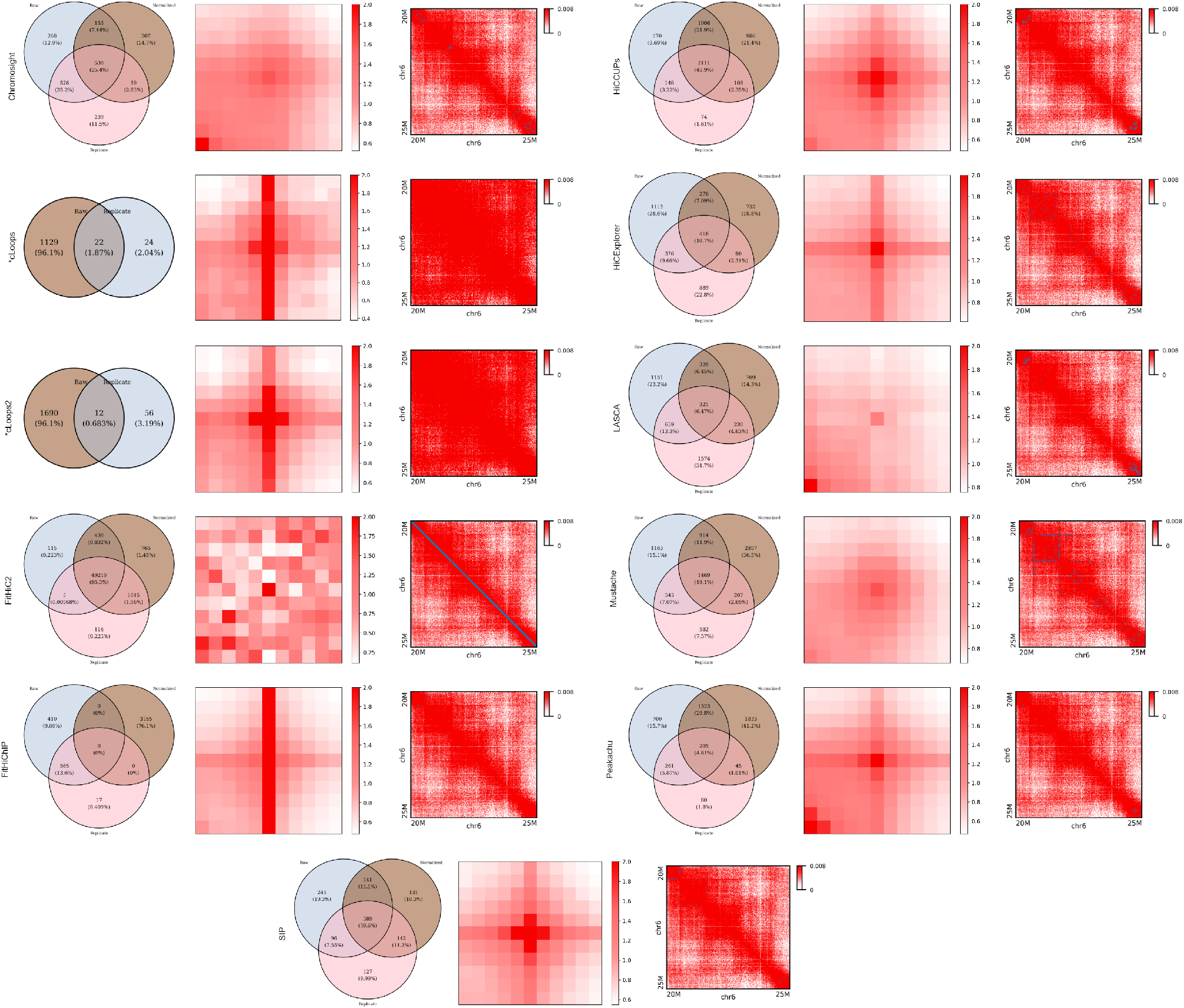
Overlap, APA, and Peak plots (from left to right) using primary GM12878 Hi-C dataset for chromosome 6 at 5KB. FitHiC2 overlaps 95.3% across the analysis. Apart from that, HiCCUPS has the highest amount (45.9%) of overlap across three different datasets and FitHiChIP does not overlap any loops. SIP (30.6%), Mustache (19.1%), and Chromosight (25.4%) have a significant amount of overlap. cLoops and cLoops2 produce results only for primary and replicate data and the overlap percentage is around 1. FitHiC2 shows enrichment in different regions and has the strong enrichment compared with other tools. cLoops, cLoops2, and FitHiChIP have enrichment in the middle vertical region and they are almost in the same shape. HiCCUPS, SIP, Peakachu, and HiCExplorer have enrichment in the middle region. Peakachu, Lasca, and Chromosight have enrichment in the left lower corner from the center and Mustache shows enrichment in the center pixel. Though FitHiC2 creates a diagonal dark straight line marking peaks, Mustache and HiCExplorer mark the highest number of peaks. Chromosight, LASCA, and SIP mark peaks in the upper left and lower right corners near 20M and 25M region.

For further analysis, we conducted the same assessment on chromosome 1 at 5KB and 10KB, and chromosome 6 at 10KB (Supplemental Figures 2, 3, 4). FitHiC2 consistently showed overlaps above 90%, reaching almost 100% for chromosome 6 at 10KB. In contrast, FitHiChIP exhibited no overlap, while cLoops and cLoops2 showed around 1% overlap. Chromosight and Mustache displayed an opposite trend for chromosome 1 and 6, increasing for chromosome 1 at 5KB and decreasing for chromosome 6 at 5KB. HiCCUPS consistently demonstrated overlaps of over 45% on 10KB resolution. Peakachu showed significant overlap on chromosome 6 at 10KB, ranging from 4.61% to 50.6%, and HiCExplorer, LASCA, and SIP consistently displayed overlaps ranging from 8% to 40% throughout the analysis.

**Figure 4.**
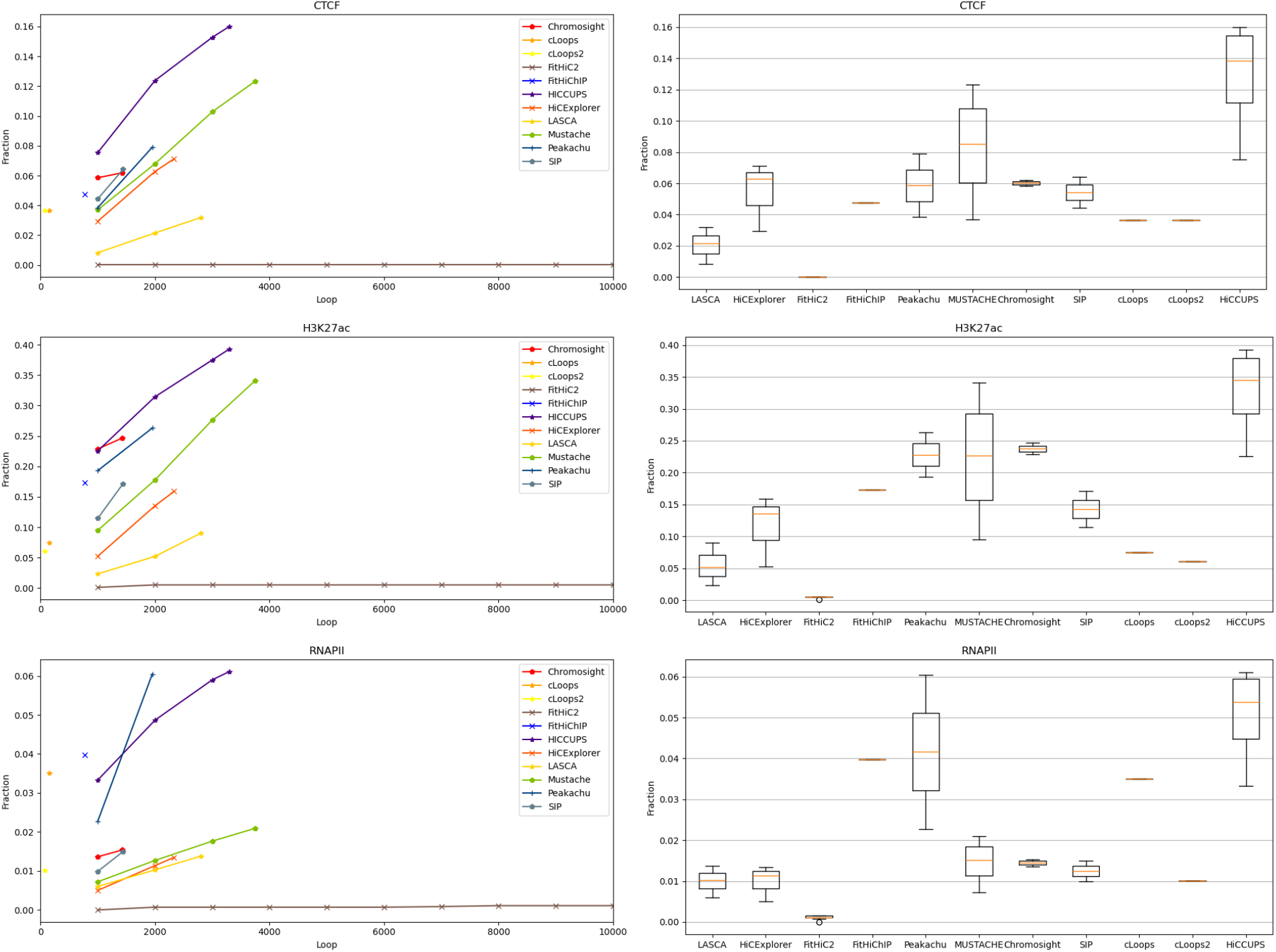
Loop recovery fraction of CTCF, H3K27ac, and RNAPII for chromosome 6 at 5KB using primary GM12878 along with their boxplot representation. HiCCUPS shows the highest fraction of recovery rate in three cases. FitHiC2 produces outliers for H3K27ac and RNAPII and has an improvement for CTCF. Mustache recovers a significant amount of fraction of H3K27ac, FitHiChIP shows a good fraction of recovery at CTCF and Peakachu recovers a good amount of fraction of RNAPII loops.

### 2.3 Peak and APA analysis

Peaks represent regions with the highest observed interactions, contributing to the formation of loops within chromatin. In our analysis, we focused on the 20M to 25M region to visualize peaks using loop lists from various tools. Figure 3-right displays peaks on chromosome 6 at 5KB resolution for primary GM12878 data. FitHiC2, Mustache, and HiCExplorer exhibit the highest number of peaks in this specific region. Chromosight, LASCA, and SIP mark peaks in the upper left and lower right corners near the 20M and 25M regions. FitHiC2 marks the highest number of peaks, forming a diagonal straight line for every dataset combination. We analyzed peak regions for chromosome 1 at 5KB and 10KB, and chromosome 6 at 10KB resolution (Supplemental Figures 5, 6, 7, 8, 9, 10, 11, 12, 13, 14, 15). For chromosome 1 at 5KB and 10KB, all tools preserve almost the same peaks, except for Mustache. For primary and replicate data, Mustache marks more peaks at 5KB compared to 10KB, which is the opposite for KR normalized data.

**Figure 5.**
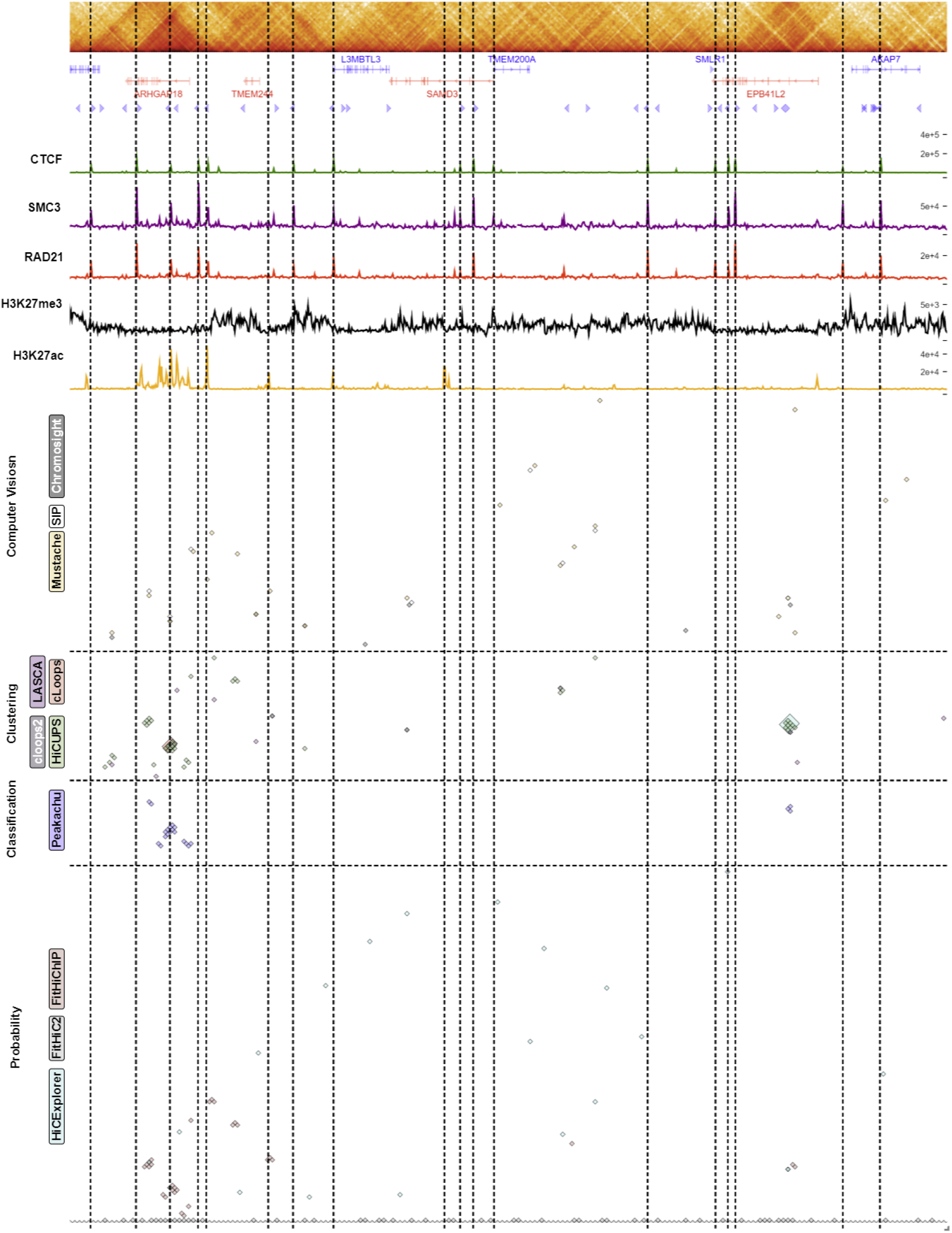
Comparison among four distinct loop caller categories in a random region (129.7M-131.6M and 62.4M - 62.5M) for chromosome 6 using primary GM12878 cell line Hi-C dataset. We depicted gene annotation, CTCF motif orientation, and ChIP signals for CTCF, SMC3, RAD21, H3K27me3, and H3K27ac below the contact map (Plotted using HiGlass). Below this biological features enrichment data, we illustrated the loops identified by each of the algoritms across different categoiries and the vertical line marks the highest enrichment point for the signals identified.

**Figure 6.**
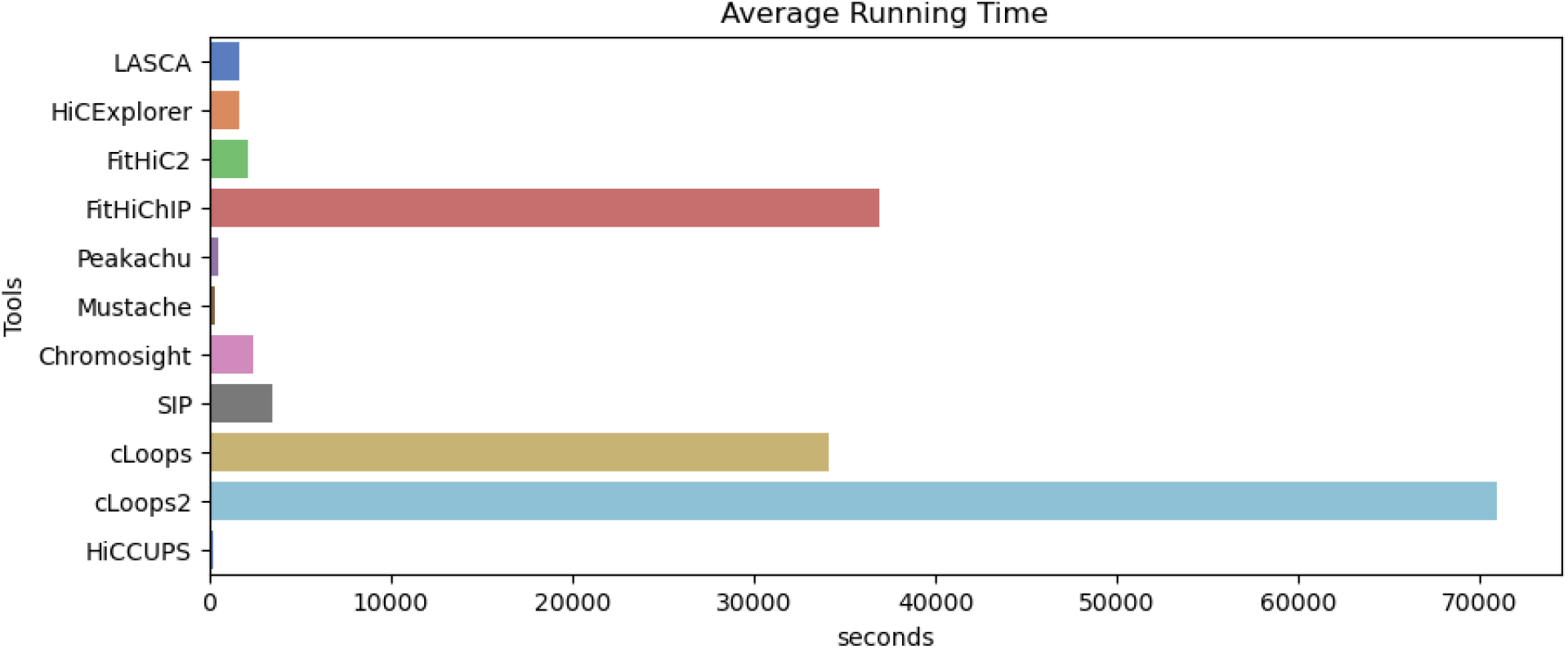
Average running time taken by all loop caller tools. cLoops2 took the highest amount of time (around 70000 seconds) to call loops. cLoops and FitHiChIP took near about 45000 seconds, and HiCCUPs, Mustache and Peakachu took the lowest amount of time (near about 100 seconds) on average. Other tools had a comparable good running time on average.

**Figure 7.**
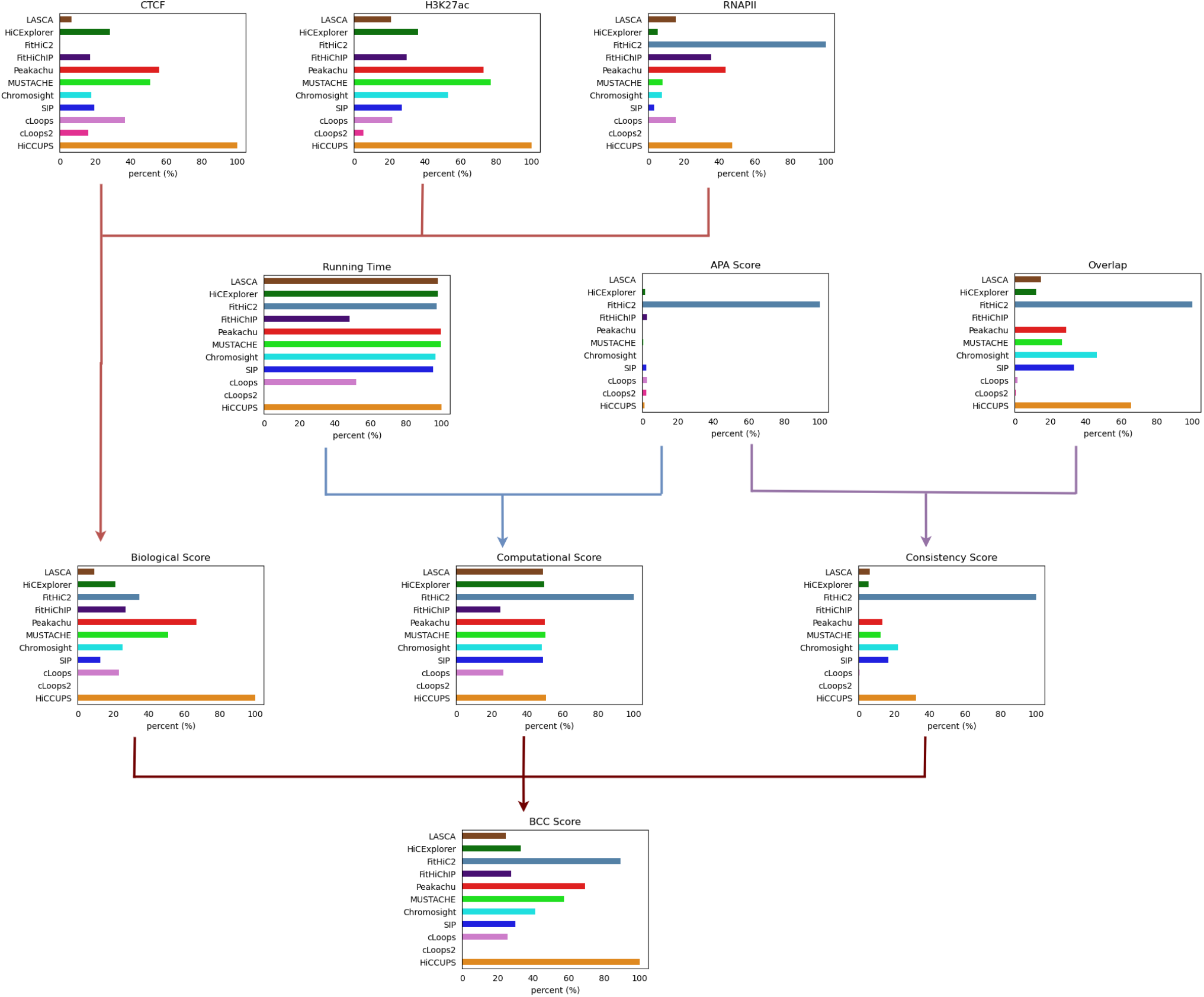
A plot showing the performance of all the algorithms across the Biological, Computational and Consistency score metrics used. The *BCC*_*score*_ representation shows the weighted aggregated performance score for all the loop callers. Every score is calculated according to the description provided in the Methods section for *BCC*_*score*_ calculation, and scaled to 100 for percentage representation, since the scores are the range 0 to 1. The scores represent the domain performance and *BCC*_*score*_ represents the overall aggregated performance of every individual tool.

To quantify the loop prediction quality, we conducted Aggregated Peak Analysis (APA) across the results. APA measures the Hi-C signal enrichment over an entire peak list, providing insights into the quality of loop lists, especially at lower resolutions. Submatrices are calculated from the Hi-C contact map file, and the sum of these submatrices produces an APA matrix. An APA score greater than 1 indicates enrichment, with darker colors in the heatmap indicating higher enrichment. Figure 3-middle shows APA plots for chromosome 6 at 5KB resolution using GM12878 primary data. FitHiC2 shows strong enrichment at different location with APA score 28.1 and cLoops (5.12), cLoops2 (3.98), FitHiChIP (4.89), and SIP (3.7) exhibit the highest enrichment in the center of the plots. Except LASCA (<1), all the tools show enrichment greater than one (Supplemental Table 11). Furthermore, we performed APA analysis at 5KB and 10KB resolution for chromosome 1 (5KB and 10KB) and chromosome 6 (10KB) using primary, KR normalized, and replicated GM12878 data (Supplemental Figure 16, 17, 18, 19, 20, 21, 22, 23, 24, 25, and 26). For normalized data, using chromosome 6 at 5KB, all tools scored greater than one and FitHiC2 (36.9) has the highest score (Supplemental Table 12). HiCCUPS (2.12), SIP (2.72), Mustache (1.52), Chromosight (1.19), and Peakachu (1.26) show gradual enrichment around the center. For 10KB data, SIP (2.77) produces a stronger central pixel color compared to 5KB resolution data, whereas other tools produce prominent plots at 5KB resolution. HiCExplorer produces almost identical visual plots for primary and replicate, KR normalized plots, and they are highly enriched at the central pixel of heatmaps. Throughout the analysis, FitHiC2 (83.73) shows strong and void enrichment at different focal points with robust enrichment at various locations (Supplemental Table 11, 12, 13).

### 2.4 CTCF, H3K27ac and RNAPII Loop Recovery

To assess the robustness of each tool to detecting relevant biological features, we calculated the recovery of specific biological features namely CTCF^39,41^, H3K27ac^24^, and RNAPII^28^ within the loops and scrutinized the results. CCCTC-binding factor (CTCF) is a transcription factor that plays a crucial role in regulating the spatial organization of chromatin. It acts as an insulator protein, helping to define boundaries between different chromatin domains. Histone 3 Lysine 27 Acetylation (H3K27ac) are proteins around which DNA is wound to form nucleosomes and are often found near the promoters of actively transcribed genes. RNA Polymerase II (RNAPII) is an enzyme responsible for transcribing DNA into RNA during the process of transcription. The presence of RNAPII is a key indicator of active transcription. Each of these molecular components serves as distinctive markers or features for the nuanced analysis of chromatin loops. Evaluating the recovery of CTCF, H3K27ac, and RNAPII loops becomes paramount, signifying the proficiency of the analytical tools in precisely identifying or predicting these features within the intricate landscape of chromatin organization. We conducted comparisons both in combination with CTCF, H3K27ac, and RNAPPII, and individually (Supplemental Figures 37, 38, 39, 40, 41, 42, 43, 44, 45). The data distribution is illustrated using boxplots accompanied by line graphs (Figure 4). HiCCUPS exhibits the highest recovery for CTCF (0.16), H3K27ac (0.40), and RNAP (0.06) loops, while FitHiC2 shows near 0.0 fractional loops in all three cases, with an outlier for KR normalized and replicate data. Notably, Mustache, Peakachu, HiCExplorer, and Chromosight also demonstrate substantial recovery of loops.

These findings were further validated with chromosome 1 at 5KB and 10KB and chromosome 6 at 10KB resolution (Supplemental Figures 27, 28, 29, 30, 31, 32, 33, 34, 35, 36). FitHiC2 exhibits an enhancement in RNAPII loops at 10KB resolution (Supplemental Figure 28, 30, 32, 34, 36), which is the highest among other tools for primary datasets. Overall, HiCCUPS recovers the highest number of loops except for RNAPII loops at 10KB resolution. FitHiChIP, using GM12878 primary data, recovers more RNAPII loops at 10KB resolution compared to HiCCUPS. Mustache recovers a substantial amount of loops across the three biological features at 5KB resolution compared to 10KB resolution. Peakachu recovers nearly the same amount of RNAPII loops as HiCCUPS. In the case of KR normalized data, HiCCUPS and Peakachu recover almost the same number of loops, except for RNAPII loops at 10KB resolution where they exhibit similar recovery rates. FitHiC2 recovers the majority of RNAPII loops at 10KB resolution from normalized data, displaying the highest number of outlier points at 10KB resolution. Mustache recovers a noteworthy number of loops, positioning itself between HiCCUPS and Peakachu in most cases. cLoops recovers the majority of CTCF loops at 5KB, and HiCCUPS and Mustache recover most of the H3K27ac loops from replicate data. FitHiC2 excels in recovering most of the RNAPII loops at 10KB resolution, while HiCCUPS recovers more at 5KB resolution. SIP, Chromosight, and HiCExplorer consistently exhibit competence in almost all cases. In summary, HiCCUPS and Peakachu demonstrate the highest recovery of CTCF, H3K27ac, and RNAPII biological markers or features in the loops identified.

We compared CTCF loop recovery and found HiCCUPS recovers most of the loops except for replicate data at 5KB resolution where cLoops recovers more loops than HiCCUPS. FitHiC2 shows zero recovery for 5KB resolution and a little improvement for 10KB resolution. FitHiChIP consistently shows zero recovery rate for KR normalized data. Mustache recovers noticeable loops comparatively at 5KB resolution whereas Peakachu took this position at 10KB resolution. FitHiChIP performs better with primary and replicated data compared to KR normalized data.

We observed HiCCUPS recovers most of the H3K27ac loops, Mustache recovers almost the same loops as HiCCUPS at 5KB resolution and Peakachu performs better than Mustache at 10KB resolution. Chromosight and HiCExplorer show a comparable recovery rate compared to other tools. FitHiChIP recovers a consistent amount of loops from primary and replicate data, but it is down to 0 using KR normalized data. Comparatively, Mustache recovers more loops compared to Peakachu at 5KB resolution and we observed the opposite direction at 10KB resolution.

HiCCUPS recovers the most amount of RNAPII loops from primary and replicates where Peakachu took this place for KR normalized data at 5KB resolution. HiCCUPS and Peakachu perform at the same level for primary data at 5KB resolution. At 10KB resolution, FitHiC2 recovers the highest amount of loops (Supplemental Figure 28, 30, 32, 34, 36). FitHiChIP recovers more loops than HiCCUPS and Peakachu at 10KB resolution from primary and replicate data and it goes down to 0 for KR normalized data. Though HiCCUPS and Peakachu recover almost the same amount of loops, HiCCUPS recovers slightly more loops than Peakachu. Mustache recovers more loops at 5KB resolution compared to 10KB (near to zero) resolution.

To visualize the biological significance within the loop area, we generated a ChIP-seq signal arrangement plot (Figure 5) for each individual category, including the contact map, gene annotation, CTCF motif orientation, ChIP-seq signals of CTCF, SMC3, RAD21, H3K27me3, and H3K27ac. At the bottom, we included loops from different individual categories. Additionally, we incorporated separate categorical ChIP-seq signal plots for four tools in Supplemental Figure 37, 38, 39, and 40. We selected a random region (129.7M-131.6M and 62.4M - 62.5M) for all these plots and marked their biologically significant areas according to their loops. Figure 5 shows that SIP and Chromosight overlap in some areas, while Mustache and Chromosight exhibit a high signal of CTCF, SMC3, H3K27ac, H3K27me3, and RAD21 loops. HiCCUPS, cLoops, cLoops2, and LASCA overlap in some regions. HiCCUPS, LASCA, and cLoops show high signals for CTCF, SMC3, RAD21, H3K27ac, and H3K27me3 in some regions within this randomly selected region, and Peakachu demonstrates signal enrichment. FitHiC2, predicting a large number of contacts across the analysis, shows a high ChIP-seq signal around the selected region. HiCExplorer and FitHiChIP display ChIP-seq signals in some regions, and ChIP-seq signal enrichment from all these tools validates our recovery analysis with a visual representation.

### 2.5 Running time on different resolution within different methods

We compared the average running time (Figure 6) of each individual tool to further assess their robustness. We ran all our tools on an Ubuntu Server operating on Intel Xeon E7-4870 @ 2.40GHz with 160 cores and 1038GB memory. Although each tool accepts different types of parameters, such as threads, chromosomes, and resolution, we attempted to compare them on the same scale using their default settings. The average running time on our server is calculated and the detailed running times are provided in (Supplemental Table 1). We observed that HiCCUPS took the least amount of time, while cLoops2 took the highest amount of time. In comparison, HiCCUPS, Peakachu, and Mustache ran within a shorter period, while LASCA, HiCExplorer, FitHiC2, SIP, and Chromosight ran within a satisfactory amount of time.

**Table 1.** A quick representation of every tool performance in different categories. Star (*) symbol tells the performance category of every individual tools. Tools are categorized in terms of performance in three categories: Excellent (***), Good (**), and Fair (*).

| Tools | Biological Feature (CTCF, H3K27ac, RNAPII) | Computation Efficiency | Consistency | Robustness in Parameters | Running Time |
| --- | --- | --- | --- | --- | --- |
| LASCA | * | ** | * | * | ** |
| cLoops | * | * | * | ** | * |
| cLoops2 | * | * | * | * | * |
| HiCEXplorer | ** | * | * | *** | ** |
| FitHiC2 | * | *** | *** | * | ** |
| FitHiChIP | * | * | * | * | * |
| Peakachu | ** | ** | ** | ** | *** |
| MUSTACHE | ** | ** | ** | *** | *** |
| Chromosight | * | ** | ** | * | ** |
| SIP & SIPMeta | * | ** | ** | ** | ** |
| HiCCUPS | *** | ** | ** | *** | *** |

## 3 Discussion

Recent advancements in 3C-based sequence technology, as highlighted by Han et al.^15^, have significantly expanded genome analysis capabilities. Loop prediction stands out as a pivotal aspect due to its relevance to various biological factors, including histone protein markers, intra and inter-chromosomal contacts, CTCF, and TAD regions. Over the past few years, a plethora of loop prediction tools has emerged, demonstrating proficiency across diverse biological aspects and datasets. In this study, we scrutinized 22 loop-calling tools, categorizing them into five distinct groups. Out of these, we successfully ran 11 tools using the same dataset and environment. Our benchmarking involved a comprehensive evaluation of biological features, encompassing the recovery results of CTCF, H3K27ac, and RNAPII, as well as considerations of running time, computational robustness, and consistency. Each tool’s default parameters were considered within the same dataset to ensure a fair alignment. Every tool has its unique capabilities; hence, we assigned a percentage score of every tool according to their performance during our analysis (Figure 7). The assessment covers three distinct categories:

### Biological Features

This includes the recovery of CTCF, H3K27ac, and RNAPII. The tools were evaluated based on how well they captured these biological features. HiCCUPS exhibited notable results in CTCF and H3K27ac recovery, with Peakachu and Mustache demonstrating considerable rates in CTCF and H3K27ac recovery (Figure 4). Though FitHiC2 excelled in contact predictions and showed superior results in RNAPII recovery, Peakachu and HiCCUPS performing well. Combining these recoveries provide an overall assessment of the biological robustness of the tools, where HiCCUPS displayed versatility across all three biological features (Figure 7).

### Consistency

This is evaluated using APA score and overlap analysis. The APA score is a computational metric that measures the Hi-C signal enrichment over an entire peak list. Tools are assessed for how consistently they perform across different datasets. The APA score revealed that FitHiC2 had the highest score over all of the tools and all the other tools maintained a score grater than one on average. FitHiChIP, cLoops, cLoops2, SIP and HiCExplorer performed well and maintained a score greater than two overall (Figure 3). Overlap analysis across three datasets (primary, replicate, and KR normalized) indicated that HiCCUPS and FitHiC2 exhibited the highest probability of overlap. Incorporating these two categories, we introduced a consistency score which is a equally weighed mean of all the assessment that measured the tools’ APA score and overlap score. Under this metric, HiCCUPS emerged as the most consistent tool. FitHiC2 showed highest consistency, and Chromosights and HiCCUPS demonstrated comparable consistency (Figure 7).

### Computational Efficiency

This category involves two key metrics, the APA score and running time. The running time analysis revealed that HiCCUPS, Mustache, Peakachu, LASCA, and HiCExplorer performed exceptionally well (Figure 6). Combining the APA score and running time, we introduced a computational robustness metric, where FitHiC2 demonstrated prominence. Except cLoops, cLoops2, FitHiChIP, all other tools yielded commendable results in the computational category (Figure 7).

We used the *BCC*_*score*_ to measure the overall performance of the tools. The *BCC*_*score*_ calculates the weighted average among the categories and provides an overall performance assessment covering biological, computational, and consistency metrics. Based on our analysis, HiCCUPS stood out as the most significant tool, closely followed by FitHiC2, Peakachu, and Mustache (Figure 7). Table 1 provides a summary of the top-performing tools across various categories. Our scoring system uses three stars to designate excellent tools, two stars for good tools, and one star for fair tools. The table includes running time as a separate metric to highlight the most efficient tools. Additionally, we benchmarked the tools parameters based on their simplicity and flexibility, noting variations in tool requirements. Some tools supports muilti-threads, normalization, multi-resolution and individual chromosome analaysis. Tools that demonstrated flexibility with a variety of parameters received higher star ratings. Memory usage was not recorded due to varying tool configurations.

## 4 Methods

Many tools and techniques have been developed for loop and peak detection. These algorithms have used different methods and approaches in their implementations based on the underlying objectives and hypotheses. Here, we categorized the tools into five distinct categories (Table 2) according to their base algorithm and we briefly describe them. All the tools are described briefly following their category in the Supplemental Document.

**Table 2.** Tools Categories by Methodology. All the tools are divided into five distinct categories according to their implementation method.

| # | Category | Tool |
| --- | --- | --- |
| A | Clustering Based | i. LOOPbit<sup>22</sup> |
|  |  | ii. LASCA<sup>23</sup> |
|  |  | iii. cLoops<sup>24</sup> |
|  |  | iv. cLoops2<sup>25</sup> |
|  |  | v. HiCCUPS |
| B | Probability-Based | i. HiCEXplorer<sup>26</sup> |
|  |  | ii. HiC-ACT<sup>27</sup> |
|  |  | iii. FitHiC<sup>28</sup> |
|  |  | iv. FitHiC2<sup>29</sup> |
|  |  | v. FitHiChIP<sup>30</sup> |
|  |  | vi. GOTHIC<sup>31</sup> |
|  |  | vii. HiC-DC<sup>32</sup> |
|  |  | viii. ZipHiC<sup>33</sup> |
|  |  | ix. NeoLoopFinder<sup>34</sup> |
|  |  | x. HMRF Bayesian caller<sup>35</sup> |
| C | Classification Based | i. FIREcaller<sup>36</sup> |
|  |  | ii. Peakachu<sup>37</sup> |
| D | Computer Vision Based | i. MUSTACHE<sup>39</sup> |
|  |  | ii. Chromosight<sup>41</sup> |
|  |  | iii. SIP & SIPMeta<sup>40</sup> |
|  |  | iv. DeepLoop<sup>48</sup> |
| E | Pile-up Procedure Based | i. Coolpup.py<sup>38</sup> |

### 4.1 Clustering Based

Clustering algorithms such as DBSCAN^49,50^, derived cDBSCAN and HDBSCAN^51^ have been used as the central algorithm in the development of some loop and peak detection algorithms such as cLoops^24^. DBSCAN algorithm does not consider the spatial organization of input data nor biased with noise data. DBSCAN performs clustering using *α*, a radius from where it will decide its core and border points, and Δ, a threshold value representing a minimum point in a cluster; otherwise, they would be considered noise. It starts scanning considering a point and expands the area considering a radius, *α*, and with this radius, all points are core points and considered to be in the same neighborhood, and if any points are not within this area, those are considered noise. This algorithm has a running time complexity of *O*(*n* log(*n*))^49^ that depends on the distance calculation algorithm and could go up to *O*(*n*^3^)^50^. We described all the clustering-based loop prediction tools in the Supplemental Document (Section 1.1).

### 4.2 Probability Based

Another category that we have identified to which most of the loop and peak detection tools belong is the Probability-based category. Specifically, tools in this category apply the binomial distribution, hidden Markov model (HMM), Cauchy distribution, and others to aid the loop and peak detection. HiCExplorer has many features with loop prediction and it uses binomial distribution, FitHiC uses statistical confidence estimation to calculate midrange intra-chromosomal contacts whereas FitHiC2 is the updated version of FitHiC. We briefly described all the tools in the Supplemental Document (Section 1.2) and in the following, we described different types of distribution algorithms.

Binomial distribution is a success or failure outcome function where the experiments iterate multiple times, and this is similar to the Bernoulli distribution. There are three preconditions for applying binomial distribution: i. observation or trials number is fixed, ii. observation or trials are independent and iii. success probability is the same for all the trials. Formally, we can state the binomial distribution as a function with a coefficient value and parameters, *t* = total number of independent trials, *r* = probability of success, *m*(1 − *r*) = probability of failure, and 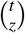= binomial coefficient

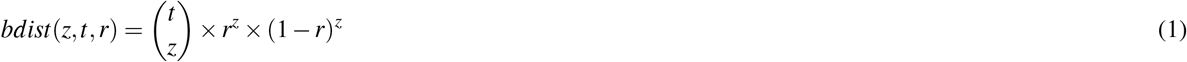

The HMM is a generalized statistical modeling formula for linear problems such as sequence, time series, and computational biology^52^. Mathematically, we can apply the HMM as there is a hidden process *H*_*n/t*_, and emission probability *P*(*S*_*n/t*_ ∈ *B*| *H*_*n*_ = *h*_*n*_*orH*_*t*_ ∈*f B* where *H*_*n*_ is a Markov process, *B* is each Borel set, and *f B* each family of Borel set. For discrete time stochastic processes, *n* ≥ 1, and continuous-time stochastic processes, *t*≤ *t*_0_. It starts from an initial state and continues until the end state generating a sequence of states based on state probabilities. This state sequence is a Markov chain where every next state depends on the current state, observing the symbol sequence hiding the state sequence.

Cauchy distribution is a continuous probability distribution closely related to the Poisson kernel. Cauchy distribution is useful in many domains such as mechanical, electronic fields, and financial analysis^53^. We can describe Cauchy distribution as

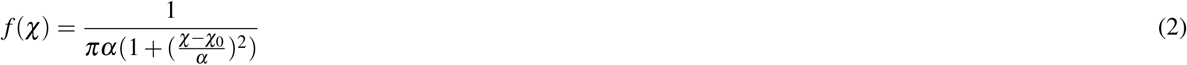

where *χ*_0_ = location parameter and *α* = scale parameter^53^. If *χ*_0_ = 0 and *α* = 1, it is called standard Cauchy distribution.

### 4.3 Classification Based

The third group of loop and peak detection algorithms that we have identified is the classification-based tools. Classification is a supervised machine learning approach that is based on training a classifier or model on labeled examples. This accurately labels unlabeled and unknown datasets introduced to this classifier. Several classification algorithms have been introduced over the years such as Decision Tree, Naive Bayes, and K-Nearest Neighbor, and are used in various domains such as fraud detection and medical diagnostics^54,55^. In bioinformatics, scientists are using classifiers to solve their problems such as cancer cells, and loop and peak detection^56,57^. Peakchu is a random forest classification-based tool to predict loops which are described briefly along with other tools in Supplemental Document (Section 1.3).

### 4.4 Computer Vision Based

Computer Vision (CV) offers access to information such as labels, object structure, shape, and much more meaningful information by analyzing images, video frames, and signals. Over the decades, computer vision algorithms have been used to make notable impacts in image classification, object detection, and recognition in robotics and autonomous vehicles. With the advent of high-resolution microscopes, we have access to biological images that can fit into computer vision algorithms for output. To support this, many CV algorithms have been proposed^39–41^ and there are certain tools for loop calling from Hi-C datasets using CV techniques such as Mustache^39^, DeepLoop^48^. We describe Mustache, SIP and SIPMeta, Chromosight, and DeepLoop in the Supplemental Document (Section 1.4).

### 4.5 Pile-up Procedure Based

Pile-up is a generalized procedure that averages a certain number of data from a given dataset such as averaging 3D points in a specific region from a 3D matrix. It describes the tendency of relation within multiple points/regions. It can be considered to be similar to the normalization technique and quantifies the averaged value with the expected one. We briefly stated Coolpup.py, a pile-up procedure-based loop detection tool in the Supplemental Document (Section 1.5).

### 4.6 Data Formats

Hi-C is a 3C-based sequence technique that facilitates high-resolution conformation capture for chromosome analysis^18,58^. This data can be used to represent and understand genome-wide features in 3D space (e.g. chromatin interaction, genomic structure, TAD, chromatin loops). To efficiently represent the Hi-C data, researchers developed .*hic*^18^, .*cool*^59^, .*mcool*^59^, and other representational formats. Hi-TrAC is another technique for genome-wide interaction profiling at a high resolution^9^. We represented all the input and output formats used in loop and peak calling tools in Table 3. The .*cool* format represents Hi-C data in three columns (bin, chromosome, and pixel) and index^59^. The .*mcool* format is a different representation on the .*cool* format having multiple-resolution data. The .hic is a highly compressed binary file for fast random access containing multiple resolution contact matrix^18^. The .*bed* and .*bedpe* are developed to represent genomic data. The .*bed* (Browser Extensible Data) format contains a maximum of 12 columns (chrom, chromStart, chromEnd, name, score, strand, thickStart, thickEnd, itemRgb, blockCount, blockSizes, and blockStarts) where the first three are required^60^. Another format is .*bedpe* (containing chrom1, start1, end1, chrom2, start2, end2, name, score, strand1, strand2, and user-defined fields) was introduced to represent interchromosomal features for variation analysis of the chromosome structure^61^. The .*sam* is a sequence alignment or a map format developed by Li et al.^62^. It is a tab-separated text format having an optional header section and alignment section. The alignment section has 11 fields (QNAME, FLAG, RNAME, POS, MAPQ, CIGAR, RNEXT, PNEXT, TLEN, SEQ, QUAL) and the @ symbol separates the header section from the alignment section. The .*bam* (binary alignment map) is the binary representation of the .*sam* format^62^. The .*hdf5* (hierarchical data format version 5) is an open-source data format that supports large, complex, and heterogeneous data in a single file and acts like a file system^63^. The .*h5* is developed based on the .*hdf5* container. It has a specific structure describing intervals, matrix, distance count, nan_bins, and correction_factors. The .*rds* (Ray Dream Studio) is a 3D object file extension that is serializable and compressible into a smaller size. The .*bedGraph* is a track format that can hold continuous-valued data such as chromosome name, start, end, and data value^60^. It is similar to the wiggle format and suitable for transnational and probability score data. The .*clpy* is the Coolpup.py defined custom data format for storing pileup results from the method pipeline^38^.

**Table 3.** An overview of the peak and loop calling algorithms from 3C-based Hi-C data. Each column denotes the information about the algorithm in order: the tool name, the year released, the 3C-based sequences data it accepts, the input data file format, the accepted input data resolution, and the output data file format. All the tools have different input and output formats, sequence data, and recommended input resolutions. It is worth noting that often many of the tools accept 3C-based data with resolutions lower than the ones stated in the table. The reported resolution for each tool is based on what was used by the authors in their manuscripts.

| # | Tool | Year | Sequence Data | Input Format | Input Resolution | Output Format |
| --- | --- | --- | --- | --- | --- | --- |
| 1 | FitHiC<sup>28</sup> | 2014 | Hi-C, ChIA-PET | .txt |  | .txt, .gz |
| 2 | HMRFBayesian caller<sup>35</sup> | 2015 | Hi-C | .txt | $\leq 10KB$ | .hdf5, .txt |
| 3 | HiC-DC<sup>32</sup> | 2017 | Hi-C | .bam | $\leq 5KB$ | .rds |
| 4 | GOTHIC<sup>31</sup> | 2017 | Hi-C | .bam, .sam | $\leq 1Mb$ | .gz |
| 5 | cLoops<sup>24</sup> | 2019 | ChIA-PET, Hi-C, HiChIP, TrAC-looping | .bedpe | $\leq 5KB$ | .txt, .pdf |
| 6 | FitHiChIP<sup>30</sup> | 2019 | HiChIP, Hi-C | .txt, .gz, .sizes | $\leq 5KB$ | .bed |
| 7 | FitHiC2<sup>29</sup> | 2020 | Hi-C | .txt | $\leq 40KB$ | .txt |
| 8 | FIREcaller<sup>36</sup> | 2020 | Hi-C | .gz, .hic, .cool | 40KB | .txt |
| 9 | Peakachu<sup>37</sup> | 2020 | Hi-C, Micro-C, ChIA-PET, HiChIP, PLAC-Seq, Capture Hi-C | .cool, .hic, .bedpe | 10KB | .bedpe |
| 10 | Coolpup.py<sup>38</sup> | 2020 | Hi-C | .bed, .cool, .mcool, .hic, .tsv | $\leq 10KB$ | .clpy |
| 11 | MUSTACHE<sup>39</sup> | 2020 | Hi-C, Micro-C | .txt, .hic, .cool, .mcool | $\leq 10KB$ | .tsv |
| 12 | SIP & SIPMeta<sup>40</sup> | 2020 | Hi-C | .hic, .mcool, file | $\leq 5KB$ | file |
| 13 | Chromosight<sup>41</sup> | 2020 | Hi-C, ChIA-PET, DNA SPRITE, HiChIP, Micro-C | .cool | $\leq 10KB$ | .tsv |
| 14 | LASCA<sup>23</sup> | 2021 | Hi-C, ChIP-seq | .cool | $\leq 10KB$ | .bedpe |
| 15 | cLoops2<sup>25</sup> | 2021 | ChIC-seq, Hi-TrAC/TrAC-looping | .bedpe | $\leq 1KB$ | .txt |
| 16 | HiC-ACT<sup>27</sup> | 2021 | Hi-C | .txt | $\leq 10KB$ | file |
| 17 | NeoLoopFinder<sup>34</sup> | 2021 | Hi-C | .mcool | 10KB | .bedGraph |
| 18 | LOOPbit<sup>22</sup> | 2022 | Hi-C | .tsv | 5KB | .tsv |
| 19 | HiCEXplorer<sup>26</sup> | 2022 | Hi-C | .bam, .sam | $\leq 10KB$ | .h5 |
| 20 | ZipHiC<sup>33</sup> | 2022 | Hi-C | .csv | $\leq 10KB$ | data frame |
| 21 | HiCCUPS<sup>47</sup> | 2014 | Hi-C | .hic | $\leq 5KB$ | text |
| 22 | DeepLoop<sup>48</sup> | 2022 | Hi-C | heatmap | $\leq 5KB$ | heatmap |

### 4.7 Analysis Methods

#### 4.7.1 Overlap

Overlap defines the common loops between different loop prediction tools’ results. Here, we used https://github.com/ay-lab/FitHiChIP/tree/master/UtilScript to draw the overlap between primary, replicate, and normalized data for a specific chromosome at a specific resolution. We used 50 window sizes to determine the overlaps. This produces results in two ways, i) comparing with a reference loop file, and ii) producing a master interaction file from the provided files merging them all together. We used the master interaction file generated from our loop files. First, it generates master interaction files from the loop files storing all the loop information and then sorting them. It receives up to 5 interaction files to draw the diagram. Next, it finds the overlap indices between the merged file and the input files and determines the unique overlap indices from the overlap indices.

#### 4.7.2 Recovery

We computed CTCF, H3K27ac, and RNAPII recovery using different loop prediction results. This recovery reports the biological consistency of a tool. The main procedure for recovery analysis is almost same as overlap analysis. Recovery analysis requires two input files i) a reference file to be matched, and ii) a loop file with q-value column. It sorts the input file with q-value and then finds the overlap indices between the loop file and the reference file. It first defines the overlap between files and then only keeps the unique overlap indices to get the overlap statistics. It uses a window size to calculate the overlap and we used 50 window size in our analysis. Then it calculates a fraction of recovery in every thousand count with the reference rows.

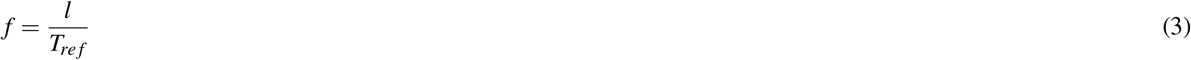

We can write it as where *f* = fraction of recovery in every thousand, *T*_*re f*_ = number of records in reference file, and *l* = length of overlap. To compute this recovery, we used https://github.com/ay-lab/Utilities/tree/main/Recovery_Plot_FitHiChIP script in our manuscript.

### 4.7.3 Peak

We used https://github.com/XiaoTaoWang/HiCPeaks to generate the peak plots. Here, we used 20M to 25M regions to observe the peaks from the loop file. First, it generates the heatmap using the contact matrix file at the given specific regions. After that, it parses the loop file to determine the positions of loops. It creates a loop table with chromosome numbers and, the start and end positions of loops. Using this table, mark the positions in the heatmap to indicate the loops.

#### 4.7.4 APA

To determine the APA score we used https://github.com/XiaoTaoWang/HiCPeaks. First, it determines peak regions from the loop file. After that, with these positions, the interaction matrix file, and the provided window size, it generates an APA submatrix. It takes each value of a square region according to the window size centering the peak position and divides each value with the mean value of those regions generating a submatrix at the end. We write it

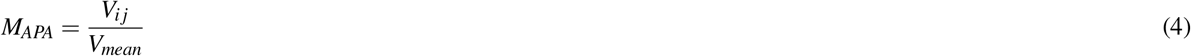

where *M*_*APA*_ = APA submatrix, *V*_*i j*_ = square region from a peak position according to the window size, *w* (we used window size, *w* = 5), *V*_*mean*_ = mean value of the square region *V*_*i j*_, and *i* = (*i*− *w, i* + *w*) and *j* = (*j*− *w, j* + *w*). Then from the submatrix, it creates a mean value list for every row of the submatrix to remove the outliers and determine the percentils. Next, it determines the average value from the submatrix and calculates the lower positions matrix using the average values up to the limit of corner size (we used 3). Finally, it calculates the APA score by dividing the average value within the window by the lower position mean value. We write it

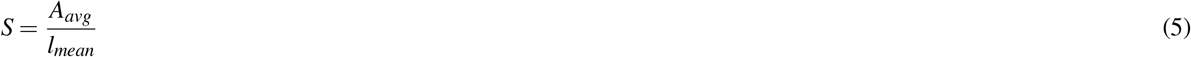

where, *S* = APA score, *A*_*avg*_ = Average APA values, and *l*_*mean*_ = lower position mean value.

#### 4.7.5 BCC Score

To determine the robustness of the tools, we categorize our analysis in three category (Biological, Consistency and Computational) and introduced *BCC*_*score*_ to compute overall score. *BCC*_*score*_ calculates the weighted average score among all the features where users can assign their weights according to their usecase to find the robustness. It is a flexible score function where user can include more categories according to their analysis. We stated *BCC*_*score*_ as

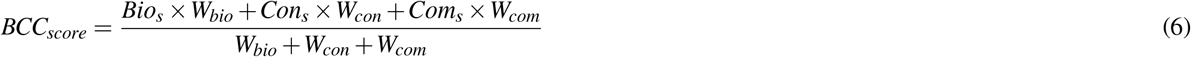

where *Bio*_*s*_ = biological feature score, *W*_*bio*_ = weight for *Bio*_*s*_, *Con*_*s*_ = consistency score, *W*_*con*_ = weight for *Con*_*s*_, *Com*_*s*_ = computational score, and *W*_*com*_ = weight for *Com*_*s*_. In our analysis, we used CTCF, H3K27ac and RNAPII as biological feature score (Equation 3) and assigned *W*_*bio*_ = 3 because the biological correctness of a predicted loop is more valuable and relevant for downstream analysis. Additionally, we assigned this value for *W*_*bio*_ since we had three features making up the *BCC*_*score*_, hence signifying it is three time more important. We anticipate that the users can modify this weight, as needed, in future analysis to signify how important they rate biological correctness among several other features they include or incorporate into the *BCC*_*score*_. We computed the consistency score using APA score (Equation 5) and overlap percentage scores; and computed the computational score using running time and APA scores with *W*_*con*_ = *W*_*com*_ = 1. The *BCC*_*score*_ is computed by normalizing all category scores through Min-max normalization. This transformation ensures that the minimum value becomes 0, the maximum becomes 1, and all other values are expressed as decimals between 0 and 1. Consequently, the *BCC*_*score*_ yields a value between 0 and 1, where higher values indicate better performance.

## 5 Data Availability

The Hi-C contact maps of GSE63525 GM12878 were downloaded from NCBI GEO. We used GSM1872886 as CTCF reference, GSE101498 for H3K27ac reference, and GSM1872886 as RNAPII reference file for biological feature analysis. These files are available in NCBI GEO. We used ChIP-seq signal data from the USCS Genome Browser which is available on HiGlass server. We used HiGLass server’s preloaded CTCF motif orientation file and gene annotation file during our analysis.

## 6 Code Availability

All scripts and programs used to benchmark these loop calling tools are available at https://github.com/OluwadareLab/Comprehensive_Loop-Caller_Benchmark

## 7 Acknowledgements

This work was supported by the National Institutes of General Medical Sciences Grant R35GM150402 to O.O.

## 8 Author contributions

H. M. A. Mohit Chowdhury conducted the analysis and wrote the manuscript, Terrance Boult initially reviewed the manuscript and Oluwatosin Oluwadare concieved, revised the manuscript and supervised this project.

## Notes

### Competing Interest Statement

The authors have declared no competing interest.

