## Supplemental Document for "Comparative study on chromatin loop callers using Hi-C data reveals their effectiveness"

#### 1 Supplemental Document

##### 1.1 Clustering-Based Tools

###### 1.1.1 LOOPbit

Galan et al.[1] developed LOOPbit for classifying CTCF-CTCF pairs. LOOPbit uses HDBSCAN and CNN algorithms in its pipeline and CNN helps to predict loop localization in the Hi-C interaction matrix. They used ChIP-seq and Hi-C data for their experiment and tested the reproducibility with Jaccard Index and calculated biologically relevant benchmarks. LOOPbit prepares its data using MEME, FIMO, and TADbit tools[2–4]. It performs compartmentalization with TADbit tools and uses APA[5] steps for generating the sub-matrices. It defines non-overlapping peaks and performs CTCF peak aggregation with *Meta-Waffle*. After this step, LOOPbit preforms analysis, deconvolution and classifies submatrices into micro-clusters or neurons with a Self-Organizing Feature Map (SOFM)[6] process. It decides the optimal parameters for SOFM considering three evaluation matrices: i. classification submatrices percentage, ii. neuron variability, and iii. compartment segregation score. LOOPbit finds the CTCF-CTCF interaction patterns with the SOFM neuron coordinates, performs clustering with Uniform Manifold Approximation and Projection(UMAP), and then applies the HDBSCAN algorithm for further clustering. It merges all the classes into the four classes of promoter, enhancer, repressed polycomb, and heterochromatin. Next, LOOPbit sections off the genome into $5KB$ bins and accordingly classifies them based on overlap $> 50bp$. One or more categories could have one bin. Finally, LOOPbit classifies these interaction bins as not expected, promoter-enhancer loop, and heterochromatin-heterochromatin.

###### 1.1.2 LASCA

Luzhin et al.[7] introduced the loop and significant contact annotation pipeline (LASCA) for identifying the loop and enhancer-promoter interactions. LASCA is a Weibull distribution-based modeling. It provides users the ability to tune the parameters and steps according to their choice. LASCA follows three basic processes in its entire life cycle. First, each diagonal corrected Hi-C matrix goes through Weibull distribution-based statistical background model. After that, LASCA calculates P-values of each pixel representing probability, and that leads to tracing model pixels in the same or higher intensity and calculating Q-values performing FDR correction in P-values. This Q-value may be corrected again, and user can set the threshold value for the Q-value with a default at 0.1. Next, LASCA clusters all the significant groups having a minimum of three pixels in each cluster by DBSCAN. It assigns pixels in neighbors by calculating the maximum Euclidean distance to be 1 between two pixels related to a particular cluster. The user specifies the cluster size, and it defines each cluster center; these center coordinates are loop coordinates. Finally, the identified loops may go through various filters such as signal over background enrichment, signal intensity, signal decay, and signal enrichment over random signals.

###### 1.1.3 cLoops

Most of the loop calling tools are confined to single data types, depend on contact matrix resolution, or require expensive hardware. Cao et al.[8] proposed cLoops to overcome these issues. They used the cDBSCAN algorithm for loop calling in 2D space and achieved $O(n)$ running time in the most ideal situation. They enhanced the DBSCAN algorithm for loop calling by adding an indexing technique to lessen noise and improve neighbor search. Like the original DBSCAN algorithm, the cDBSCAN algorithm uses two inputs (radius and threshold) with the same meaning. In addition, it marks minimum 2D genomic coordinates, indexes each coordinate, and assigns each point to a square. Then, cDBSCAN determines the noise squares within two steps: i. finding the potential noise indexes, and ii. finding the already detected noise indexes for cross-checking. For noise identification, they used the K-Nearest Neighbor idea and introduced an indexing technique that reduces the search space. The cDBSCAN performs clustering the same as the original DBSCAN algorithm with $3l \times 3l$ square for neighbor search where $l =$ square side length. The main cLoops is a two-step loop calling algorithm and takes mapped PET data formatted in *.bedpe*. It uses the cDBSCAN for loop and peak calling, and the result could be used to find significant combinations and loop boundaries. The cLoops compares the candidate's loops' PETs number with the permuted local backgrounds (PLBs) for determining their significance.

$$Hypergeometric\ Test = 1 - \sum_{n=0}^{P_{i,j}-1} \frac{\binom{Tr_j}{n}\binom{Tp-Tr_j}{Tr_i-n}}{\binom{Tp}{Tr_i}} \tag{1}$$

where $P =$ linking anchor PETs, $Tr =$ specific region total PETs and $Tp =$ total PETs

$$Poisson\ Test = 1 - \sum_{n=0}^{P_{i,j}-1} \frac{\eta^n \varepsilon^{-\eta}}{n!},\ \eta = \sum_{i,j} \frac{P_{i,j}}{\rho_{i,j}} \tag{2}$$

where $\rho =$ permutation region number

$$Binomial\ Test = 1 - \sum_{n=0}^{P_{i,j}-1} \binom{Tp}{n} \psi_{i,j}^n (1-\psi_{i,j})^{Tp-n} \tag{3}$$

where $\psi =$ single PETs link observation possibility. cLoops uses a hypergeometric test, Poisson test, and binomial test for determining statistical significance of candidate loops to increase the precision.

###### *1.1.4 cLoops2*

cLoops2[9] is an updated version of cLoops[8]. It has extra functionalities that can be run separately such as outputting uniquely high-quality mapped PETs from the FASTQ file. Cao et al.[9] improved the DBSCAN algorithm for cLoops2 and named it blockDBSCAN, which starts with marking the smallest and biggest coordinates of PETs. It projects the shorter distance peaks in a 2D diagonal line and long interaction loops in space distance, removes the noisy points, and automatically calculates the cluster numbers. The blockDBSCAN classifies clusters into two groups: candidate peaks and candidate loops. It saves cLoops-specific input for analysis and can be reverted to the PETs-specific data. Here, for peak calling, Cao et al.[9] used blockDBSCAN that performs with a combination of different radii and minimum points in a cluster with $PETs > 1KB$. cLoops2 calculates the statistical significance in the nearby or same region using the Poisson test for all candidate peaks. It describes a candidate's peaks as overlapping with significant Poisson P-value having the highest RPKM value. In the end, cLoops2 rectifies all the Poisson P-values with Bonferroni correction. For calculating the significant peak, Cao et al.[9] took $P < 0.01$ by default. It calculates precision, sensitivity, and F1 scores for analyzing the result. For loop calls, cLoops2 uses blockDBSCAN with different radii and minimum points in a cluster. The blockDBSCAN provides candidates loops after performing numerous clustering rounds, and cLoops2 uses hypergeometric and Poisson tests like as cLoops[8] and an updated binomial test.

###### *1.1.5 HiCCUPS*

HiCCUPS[5] is a tool uses clustering to analyze every pixel in a Hi-C contact matrix and compare it to the pixels around it. It calculates the higher enriched pixels and ensure that this higher enrichment is not from a large structural features. It calculates this enrichment compared with the lower left corner and this specific enrichment should be $\geq 50\%$ from the lower left corner enrichment. HiCCUPS also validate this enrichment with multiple hypothesis testing such as FDR $< 10\%$ and comparing with four neighborhood pixels- i) lower left, ii) left and right, iii) above and below and iv) a donut. This enrichment results in a regions of 5-20 contiguous interactions.

##### 1.2 Probability-Based Tools

###### *1.2.1 HiCExplorer*

Wolff et al.[10] introduced HiCExplorer for loop detection in Hi-C data with a high loop detection percentage using low memory and multicore CPU architecture. HiCExplorer reduces the search space for achieving high throughput and has a good ability to distinguish true interaction and noise. They stated that HiCExplorer can detect almost all the true interactions considering relative genomic distance and noise regions using chromosome 1 on GM12878[10]. At first, HiCExplorer normalizes the the interaction height of every genomic distance by calculating observed over the expected matrix. For expected matrix calculation, the algorithm considers: i. non-zero contacts, ii. all contacts, and iii. different occurring ligation events which are similar to HOMER[11] normalization. Then HiCExplorer applies continuous negative binomial distribution, $\chi_\triangledown \sim CNBD_\triangledown(m_\triangledown, n_\triangledown)$ $\forall \triangledown = |x-y|$ for candidate selection from the normalized data where $\triangledown$ is the genomic distance. They modified the discrete negative binomial distribution function configuring with a modified gamma function[12,13] for making it continuous. It calculates P-values with binomial density function and accepts candidates having a P-value less than a threshold considering only observed versus expected values. The algorithm considers the whole neighborhood of $a^2$ area, pools all candidates from a neighborhood, and divides them into peak and background region. After that, HiCExplorer again divides this neighborhood into vertical and horizontal regions similar to HiCCUPS technique[5]. The peak and neighborhood square size is defined by their peak width and window size. If a candidate satisfying $\bar{P} > \bar{B}$ condition where $\bar{P}$ is mean peak and $\bar{B}$ is mean background, they will be accepted as a loop.

###### 1.2.2 FitHiC

Ay et al.[14] described FitHiC in 2014 by combining the random polymer looping effect and technical biases. FitHiC calculates statistical confidence in midrange intra-chromosomal contacts and requires contact count, number of distinct inter-chromosomal locus pairs, and total number of observed contact counts for this calculation. It considers null probability $= \frac{1}{\sigma}$, where $\sigma =$ number of distinct inter-chromosomal locus pairs to calculate the probability by i. creating a specified equal occupancy bin by dividing the locus pairs, ii. calculating average contact count per locus pair, prior contact probability, and the average interaction distance over all locus pairs, and iii. applying a univariate spline to the derived points. From this point, FitHiC calculates the P-values of each bin and inter-chromosomal contact. It synthesizes the derived P-values to get a signal ranking for the observed contacts set. Finally, FitHiC corrects the combined P-values with the Benjamini Hochberg procedure and produces a Q-value. This Q-value defines the statistical confidence of contacts and reveals the target loops.

###### 1.2.3 FitHiC2

FitHiC2 is a versatile tool for analyzing intra-chromosome, inter-chromosome, and all-region interaction from Hi-C data. FitHiC2 is not limited to a minimum sequencing depth or any contact-maps resolution. It is capable of computing the statistical confidence of Hi-C datasets with random polymer looping effects and potential technical biases. Kaul et al.[15] introduced this improved version of FitHiC for detecting chromatin contacts without any parametric assumption. First, FitHiC2 reads user-defined ranged non-zero interaction counts and builds a genomic distance index and contact counts. It records specified ranges and places the contact counts into equal occupancy bins. Next, FitHiC2 reads the fragments file for computing equal occupancy bin by enumerating all possible fragment pairs for non-fixed-size data or looping through the possible fragment pairs for fixed-size data. This ultimately results in an average contact count and average distance. Then, FitHiC2 reads and stores bias value files according to their availability. After that, the algorithm applies null model univariate cubic spline function for each bin to the derived average distance over average contact count. It ensures that this spline is non-increasing by applying antitonic regression. Finally, FitHiC2 calculates the P-value. For this P-value, it reads each entry from the contact count file, calculates the prior contact probability and multiplies this with the bias value, and finally fits this corrected P-value into a binomial distribution. It outputs these derived P-values and corresponding Q-values by appending them to the contact counts file. If a user defines a specific number of pass parameters, then an extra spline fit process is performed into the null models.

###### 1.2.4 HiC-ACT

Lagler et al.[16] created HiC-ACT with the aim to improve the chromatin interaction result by postprocessing the results from other methods (e.g., FitHiC2). These methods should consider interaction as independent to be postprocessed by HiC-ACT. For all sequencing depths, HiC-ACT shows a greater improvement in maintaining sensitivity and insignificant loss in precision. It uses aggregated Cauchy combination test (ACT)[17] and uses non-negative weights P-values from linear combination transformation. Under null hypothesis, Liu and Xie[17] proved that ACT is a standard Cauchy distribution, and using Cauchy distribution, test statistic and P-value could be transformed by each other. For test statistic, it conditions the smoothing window using the HiCRep[18] method considering the resolution and uses Gaussian Kernel weight function. First, HiC-ACT gathers results from a standard peak caller without spatial dependency and requires identifiers of bin pair and P-values. Next, the algorithm sets the distance value depending on the resolution. Then it selects interest pairs considering a threshold greater than the P-value between two bins. Finally, HiC-ACT determines all pairs and calculates weights, test statistics, and P-value for derived bin pairs.

###### 1.2.5 FitHiChIP

Bhattacharyya et al.[19] introduced the FitHiChIP method in 2019 for loop detection. FitHiChIP models non-uniform coverage and genomic distance scaling at the same time and has reproducibility. It calculates the contact probability within two steps: i. monotonic spline fitting[14] for modeling the contact probability decay with increasing genomic distance and ii. applying a regression on observed contact and the bias values. First, FitHiChIP sorts all the locus pairs and applies an equal occupancy binning across the total contacts. It finds the sum of contact counts and average contacts per locus pair. Next, the algorithm calculates a prior contact probability using these average contacts and finds the average interaction distance for all possible pairs. Finally, FitHiChIP uses a univariate spline filtering for expected over prior contact probability. For this spline fitting and equal occupancy binning, FitHiChIP applies a background model (e.g., peak to all locus pairs, peak-to-peak loops). It determines statistical significance without bias regression and with bias regression. The algorithm calculates the contact probability with binomial distribution[14] without applying the bias regression and uses the prior contact probability directly from the spline fit. From this procedure, FitHiChIP results in P-values and derives Q-values with the Benjamini–Hochberg procedure[20]. If it considers the bias (e.g., coverage, ICE), it applies bias regression in equal occupancy bin for getting bias values. It either applies HiChIP coverage to the mean coverage ratio or a matrix balancing method such as ICE[21] for bias values. FitHiChIP uses a linear regression model for all equal occupancy bins, calculates regression coefficients, and uses these in a smoothing spline. Then it determines expected contact counts and expected contact probability. Finally, it calculates the statistical significance with the binomial distribution.

##### 1.2.6 GOTHiC

Mifsud et al.[22] introduced GOTHiC in 2017 considering known and unknown biases such as density of restriction sites, cleavage efficiency, mappability and believed these biases had an independent effect in each pair. GOTHiC uses a binomial probabilistic model for achieving precision across the loop detection procedure. It determines the probability of given read pairs using random ligation and observed over expected ratio. First, within a specific distance (default $10KB$) and in the same fragment, GOTHiC applies filtering read pairs for removing self-legations, dangling ends, re-legations, and incomplete digestion products. Next, it calculates the spurious read pair probability using two-locus relative coverage and read pairs fraction. Then the algorithm derives probability of read pairs with binomial cumulative density and results in P-value. Finally, GOTHiC results in Q-value applying Benjamini-Hochberg multiple-testing and defines statistically significant interactions.

##### 1.2.7 HiC-DC

Carty et al.[23] introduced HiC-DC in 2017. They used hurdle regression (zero truncated negative binomial regression)[24] for Hi-C contact count resulting in a null or background model. HiC-DC can handle systematic biases such as GC content and mappability. The algorithm takes Hi-C intra-chromosomal interactions contact map as input where every bin has its associated covariates vector. First, HiC-DC takes a random variable that follows a zero-truncated negative binomial distribution. Next, it defines the regression model with negative binomial distribution having a dispersion parameter and negative binomial mean parameter. It considers GC content and mappability features while calculating negative binomial mean parameters and this is a log-linear function. Then, HiC-DC calculates a third-order B-spline by establishing a relation between contact significance and genomic distance. After this, it refits the model for increasing statistical power after removing the bin counts greater than 97.5% null distribution percentile. Finally, HiC-DC calculates the P-value for each bin and then retrieves the adjusted P-value with the Benjamini-Hochberg procedure. It also performs known genomic and epigenomic labelling. For labelling, it performs IDR for getting reproducible peaks of each dataset and creates a peaks catalog by merging the GRanges objects. HiC-DC uses this catalog to label the bins as promoter, exonic, intronic, and distal intergenic. While labelling, it retrieves the GRanges object from the catalog.

##### 1.2.8 ZipHiC

ZipHiC is a HMM and Approximation Bayesian Computation (ABC)[25] based algorithm for identifying interaction in 2D space. Osuntoki et al.[26] introduced this algorithm in 2022. ZipHiC uses a mixture model with K-components mixture density. Here, zero-inflated Poisson (ZIP) distribution is the first component and Poisson distribution the second. The algorithm describes this mixture model with a latent variable, $L_{mn}$. They used three components in their experiment, but it can accept two or more components. They also assumed that noise follows ZIP distribution in most of the cases, and this noise could be corrected with a log function. ZipHiC uses the hidden Markov random field (HMRF) based on HMM for hidden components and implements Potts model[27] for spatially dependent data as its prior. It models the latent variables as a 2D HMRF, and this latent variable depends on the neighbors' status. It defines the neighboring as an influence sum and uses a partition function as normalization constant. For this normalization constant, ZipHiC uses the ABC model. It implements the Empirical Bayes approach and conventional Bayesian approach, and those are based on posterior distribution proportional to the product of the prior and likelihood.

##### 1.2.9 NeoLoopFinder

In 2021, Wang et al.[28] published the NeoLoopFinder for detecting chromosome loops induced with structural variations (SVs) along with interchromosomal translocations, large deletions, and inversions. NeoLoopFinder uses a modified matrix balancing technique and can handle signal distortion, mappability, GC content, and restriction fragment size. It can draw heatmaps, genes, epigenomic loops and tracks. It takes the Hi-C contact matrix and SVs, and produces genome-wide CNV profile and segments, linked SV events chain, corrected Hi-C matrix, chromatin loops and enhancer-hijacking events. They developed HMM-based CNV segmentation module with a better performance. NeoLoopFinder starts with CNV profiles and segments inference in the Hi-C contact matrix, and goes through matrix balancing process. It uses HiNT[29] and HiCnv[30] for CNV profile computation. In the next step, sub-matrices are flipped or rotated according to breakpoints type and orientation. They initiated a graph-based algorithm for correctly combining the order of each SV that mitigates the complexity of SVs. They developed a linear regression model for balancing signals and bringing them into a similar range caused by the allelic effect. After this, NeoLoopFinder identifies loops, and starts with integrating Peakachu[31] and includes a pre-trained model. It trains this model considering resolutions, window sizes, and in-situ and dilution protocols.

##### 1.2.10 HMRF Bayesian caller

Zheng Xu et al.[32] introduced this algorithm in 2015 with a basis of finding intra-domain interactions. They showed this algorithm has great power to find peaks accurately along with reproducibility and statistical power. This algorithm uses Hi-C generated contact frequency matrix between fragments pairs where they considered observed $= O_{mn}$ and expected $= E_{mn}$ contact frequencies between $m$ and $n$ fragment, and $1 \leq m < n \leq T$ and $T =$ total number of fragments. It has a binary indicator variable, $X_{mn}$ and this has two values. Value 1 represents a peak and -1 represents a non-peak. Zheng Xu et al. conjectured that observed contact frequencies use a negative binomial distribution, $O_{mn} \sim NBD(M_{mn}, \alpha)$ where $\alpha$ is over-dispersion parameter, $M_{mn}$ is mean of $O_{mn}$ with a variance $M_{mn} + M_{mn}^2/\alpha$. They also believed that there is an effect of the status of $X_{mn}$, and this is derived with a log function of $O_{mn}$ and $E_{mn}$ where they used FitHiC or ICE[21] methods. This algorithm uses the HMRF model for identifying local spatial dependency and finds out all 2D peaks. It stretches the HMRF model from 1D space to 2D space maintaining the concept of the original HMM or Bayesian hidden Ising model. It follows Ising prior[33] model for binary indicator variable, $X_{mn}$ which depends on four neighbor fragment pairs, $\{(i-1, j), (i+1, j), (i, j-1), (i, j+1)\}$. This algorithm measures the cluster level among binary variables for inverse temperature parameter[34], and this parameter's large value indicates a tightly clustered configuration of $X_{mn}$, and zero values indicate independent uniform prior. If there is no cluster, this algorithm calibrates the inverse temperature parameter close to zero without losing any power. This algorithm uses the Metropolis-Hastings algorithm for deriving all the parameters except the inverse temperature. It uses the Bayesian approach[35] for posterior distribution-based parameter inference. First, this algorithm specifies the prior and re-parameterizes by doing inversion of over-dispersion parameters, which provides convenience and computational efficiency. This algorithm employs weak prior with large variance by default, and there is little impact of priors in the final peak calls. It combines the binomial distribution mixture with conditional independence assumption and introduces the posterior probability function.

#### 1.3 Classification-Based Tools

##### 1.3.1 FIREcaller

In 2020, Crowley et al.[36] introduced FIREcaller to find the frequently interacting regions (FIREs) from Hi-C contact matrix. This is a classifier tool developed with *R* and provides continuous FIRE scores, dichotomous FIREs, and super-FIREs. The input matrix is prepared by dividing the genome into non-overlapping consecutive b-sized bins[37]. At first, in each bin, FIREcaller determines the total local[37] cis-interactions number and local threshold value, and is mostly dominated by contact domain exert influences. This algorithm provides an option to set user-specific cis-interacting region's upper bound. Next, bins go through a filtering process that creates some systematic bias (e.g., GC content, mappability). It removes 0 GC content, mappability, and effective fragment length along with $> 25\%$ neighborhood bins, $< 90\%$ mappability, and overlapping bins. After that, FIREcaller performs within-sample normalization using HiCNormCis[37] that uses Poisson regression process to synthesize the systematic bias. It allows negative binomial regression model for bias removal and accepts normalized data as input. If this algorithm finds multiple datasets, it performs quantile normalization across samples of the normalized cis-interactions. Later, FIREcaller computes the Z-score from normalized cis-interactions, and P-values using the standard normal distribution and classify the bins as FIREs having P-value $> 0.05$. It can identify super-FIREs by adding all $-\ln(P - values)$ and ranks least to most interactive FIREs inspired by ROSE[38]. It identifies different FIREs according to fold change $> 2$ and P-value $< 0.05$ and visualizes FIREs and super-FIREs.

##### 1.3.2 Peakachu

In 2020, Salameh et al.[31] published supervised learning-based loop detection tool, Peakachu. It is a random forest classification algorithm. According to Salameh et al., Peakachu achieved good results in their analysis and required less computational power. This algorithm gives the facility of tuning the pixel probability value that generates the higher or lower loop count. It requires $10KB$ binned Hi-C matrix and positive training set interaction list as input. It applies an $11 \times 11$ window for collecting interaction from the Hi-C matrix and discards those training lists whose Hi-C value is $< 10\%$. Peakachu derives the positive classes and defines negative classes by collecting nonzero centers and windows with random coordinates. It decomposes each sample into $2n+1$ feature vector, where $n =$ sample feature space radius and appends P2LL with each feature vector. It uses the *sci-kit* tool and fits training datasets in random forests. It divides these datasets into separate test and training datasets for avoiding over-fitting. After this, Peakachu derives the optimal hyperparameters and uses Matthew's Correlation Coefficient for picking the best model. It calculates a score of each feature vector and applies a greedy algorithm. This greedy algorithm first defines the anchor regions according to P-value and applies the DBSCAN algorithm. After running DBSCAN within any two connected regions, Peakachu counts the candidate pixels and finds peaks.

#### 1.4 Computer Vision-Based Tools

##### 1.4.1 MUSTACHE

In the real world, it is difficult to determine an object without its scaler position. Ardakany et al.[39] introduced MUSTACHE tools to overcome this problem. They presumed chromatin loops could be found in blob-shaped object interaction regions and developed this tool followed by a Scale-space framework with multiple parameters such as genomic region, CTCF-binding, size, and regulatory elements. MUSTACHE tries to identify regions of interactions with high statistical significance. It applies a local Z-normalization technique and re-scales interactions with a logarithm function. As a result, this algorithm gets a scale-space representation of normalized contact maps. MUSTACHE convolves normalized contact map with Gaussian to get scale-space representation, which ultimately produces smoothed contact map and then computes difference-of-Gaussians (DoGs). This tool employs two space-space octaves and computes P-value for each pixel with Lapasian distribution on each DoGs scales. After calculating DoGs, MUSTACHE looks for local maxima in 3D space; these maxima are considered candidate loops, and others are discarded. Then, candidate loops go through different types of filters to get high confidence and locally enriched loops. Finally, MUSTACHE i. removes candidate loops that do not belong to at least two consecutive scales, ii. discovers connected components using 8-connectivity and removes those candidate loops in sparse regions, and iii. discards candidates having contact counts $< 2\times$ expected counts and applies Benjamin-Hochberg[20] for P-value correction.

###### *1.4.2 SIP & SIPMeta*

Rowley et al.[40] developed the Significant Interaction Peak caller (SIP) and SIPMeta for identifying and characterizing loops. These two tools are platform independent and have efficiency in terms of time and memory consumption. They are not biased with noise and sequence depth. The SIP method reclaims raw Hi-C signals from the *.hic* (Juicer[41]) file, user-defined resolution, and normalization technique. It uses the sliding window technique for genome analysis, and user defines resolution and matrix size, which leads to the size of the window. It creates images using observed-expected value and applies a distance-normalization formula for calculating the central loop value. This mainly uses an image processing algorithm for candidate loop listing and applies filters to this list. First, SIP applies Gaussian blurring for making the signals smooth and then applies contrast enhancement[42] method such as White top-hat. Lastly, SIP applies a minimization and maximization filter[42] and homogenizes the background. These processes provide corrected images, and with these images, this algorithm creates a candidate loop list with the maximum detection algorithm offered by ImageJ[42]. These loops go through several filters before image processing (e.g., eliminating pixels having insufficient data, removal of pixels that do not have an increasing interaction). SIP filters the candidates' loops considering various parameters (e.g., KR value, Poisson CDF function, PA score). SIPMeta first infers bin size from loop file or creates image using the *.bedpe* file from the SIP procedure or user can specify the *.hic* file. After this step, SIPMeta tests all signals within a specified distance, calculates the APA[5] score, and produces an average matrix. With this matrix, SIPMeta generates square and bullseye plots using *bullseye.py*. In bullseye plots, SIPMeta has a ring segment representing $4\times N$ bins where $N$ is Manhattan distance and these rings' Z-score is calculated separately along with ADA score having Z-score $> 1$.

###### *1.4.3 Chromosight*

Matthey et al.[43] introduced Chromosight to detect all types of patterns in various genome-wide contact maps such as hairpin-like configuration in bacteria and yeast. They developed Chromosight based on CV having higher sensitivity in synthesis data and working well with data from different protocols without prior training data. Before going to the detection algorithm, it uses the ICE[21] algorithm for balancing the whole matrix, and this accounts for the associated biases. Then Chromosight divides each pixel with the mean of its diagonal and calculates observed over expected contact ratio. It discards the contacts that are above the user-defined threshold distance. Next, Chromosight calculates the Pearson coefficients with a Convolution algorithm. This algorithm convolves the template over the contact map and calculates the correlation coefficients. Finally, Chromosight separates the high correlation foci by i. discarding the correlation coefficient images less than a correlation coefficient threshold value, ii. generating adjacency graph with the remaining points, iii. labelling the nonzero pixels as contiguous foci with the Connected Component Labelling (CCL)[44], and with greater or equal to two pixels focus are kept, iv. detecting the patterns having the highest coefficient pixel and filters the overlapped or closed patterns, and v. sorting out candidate local maxima and filters out neighboring candidates.

###### *1.4.4 DeepLoop*

DeepLoop[45] is a deep learning based multidimensional tool which can predict chromatin interaction using low depth Hi-C heatmap. It can correct bias data and enables loop resolution. They introduced LoopDenoise, a convolutional autoencoder architecture having five convolutional layer. LoopDenoise has two encoding layer with, two decoding layer and one final layer having eight $13\times 13$, eight $2\times 2$ and one $13\times 13$ filters respectively. They used the MaxPooling layer to pull up the main features and applied ReLU activation function. DeepLoop accepts the HiCorr heatmap as input and pass it to the LoopDenoise autoencoder architecture. DeepLoop applied in different Hi-C data sets such as micro-C and have a comparable convergence and could discover loops that are escaped for X-inactivation.

##### 1.5 Pile-up Procedure-Based Tools

###### *1.5.1 Coolpup.py*

Sequencing depth is a major limitation of 3D genome organization analysis. Coolpup.py uses the pile-up procedure to mitigate this limitation. Flyamer et al.[46] introduced this tool available as a Python package. Coolpup.py removes technical biases or uninteresting biological signals. It has an isolated CLI for visualization which makes it an adaptable tool. First, Coolpup.py initializes a sparse matrix from input, defines regions, and tracks the number of summed-up regions during the process. It records the window coverage according to the specification and can rescale the window in the required shape. Next, Coolpup.py divides the total sum of the results by the total number of windows, applies coverage normalization, and removes the contact probability distance dependency according to specification. After this, it averages the expected matrices for getting the normalized matrix or uses randomly shifted control regions. Finally, Coolpup.py divides this matrix with normalized matrix for distance effect removal.

#### 2 Supplemental Tables

**Table 1.** Running time of every individual loop caller using primary GM12878 dataset.

| Tools | 10KB (Full) | 5KB (Full) | 10KB (chr1) | 5KB (chr1) | 10KB (chr6) | 5KB (chr6) |
|---|---|---|---|---|---|---|
| LASCA | | | 1014.061633 | 3218.792016 | 571.7460418 | 1722.742106 |
| HiCExplorer | | | 1602 | 4415 | 1198 | 3503 |
| FitHiC2 | | | 1072 | 1464 | 764 | 1015 |
| FitHiChIP | 30693 | 44030 | | | | |
| Peakachu | | | 673 | 1195 | 501 | 892 |
| MUSTACHE | | | 254 | 507 | 176 | 334 |
| Chromosight | 6440 | 7322 | | | | |
| SIP | 353 | 358 | | | | |
| cloops | | | 1510 | | 1196 | |
| cloops2 | | | 2338 | | 1030 | |
| HiCCUPS | | | 192 | 380 | 128 | 235 |

**Table 2.** Running time of every individual loop caller using KR normalized GM12878 dataset.

| Tools | 10KB (Full) | 5KB (Full) | 10KB (chr1) | 5KB (chr1) | 10KB (chr6) | 5KB (chr6) |
|---|---|---|---|---|---|---|
| LASCA | | | 930.989547 | 3393.544896 | 583.7422869 | 1694.471702 |
| HiCExplorer | | | 378 | 1741 | 318 | 1646 |
| FitHiC2 | | | 2514 | 3619 | 1709 | 2423 |
| FitHiChIP | 32792 | 45553 | | | | |
| Peakachu | | | 270 | 431 | 189 | 280 |
| MUSTACHE | | | 238 | 533 | 195 | 347 |
| Chromosight | 160 | 333 | | | | |
| SIP | 3515 | 6338 | | | | |
| HiCCUPS | | | 196 | 393 | 128 | 239 |

**Table 3.** Running time of every individual loop caller using replicate GM12878 dataset.

| Tools | 10KB (Full) | 5KB (Full) | 10KB (chr1) | 5KB (chr1) | 10KB (chr6) | 5KB (chr6) |
|---|---|---|---|---|---|---|
| LASCA | | | 955.6488478 | 3408.026108 | 583.619509 | 1683.847833 |
| HiCExplorer | | | 526 | 2589 | 338 | 1690 |
| FitHiC2 | | | 2235 | 3011 | 2532 | 3158 |
| FitHiChIP | 26565 | 41551 | | | | |
| Peakachu | | | 254 | 378 | 183 | 252 |
| MUSTACHE | | | 190 | 438 | 123 | 270 |
| Chromosight | 162 | 325 | | | | |
| SIP | 3755 | 6407 | | | | |
| cloops | | | 80137 | | 53667 | |
| cloops2 | | | 196596 | | 83716 | |
| HiCCUPS | | | 194 | 392 | 129 | 241 |

**Table 4.** Average running time of every individual loop caller using GM12878 dataset.

| Tools | Average |
|---|---|
| LASCA | 1646.77 |
| HiCExplorer | 1662 |
| FitHiC2 | 2126.33 |
| FitHiChIP | 36864 |
| Peakachu | 458.17 |
| MUSTACHE | 300.42 |
| Chromosight | 2457 |
| SIP | 3454.33 |
| cloops | 34127.5 |
| cloops2 | 70920 |
| HiCCUPS | 237.25 |

**Table 5.** Loop count of every individual loop caller using primary GM12878 dataset.

| Tools | 10KB (chr1) | 5KB (chr1) | 10KB (chr6) | 5KB (chr6) |
|---|---|---|---|---|
| LASCA | 1332 | 2801 | 962 | 1814 |
| HiCExplorer | 1970 | 2334 | 1284 | 1663 |
| FitHiC2 | 63150 | 66440 | 46812 | 50260 |
| FitHiChIP | 3693 | 770 | 3175 | 952 |
| Peakachu | 2312 | 1953 | 1705 | 1633 |
| MUSTACHE | 2018 | 3749 | 1570 | 2490 |
| Chromosight | 951 | 1426 | 614 | 866 |
| SIP | 249 | 1442 | 310 | 1040 |
| cloops | 154 | | 35 | |
| cloops2 | 76 | | 62 | |
| HiCCUPS | 3890 | 3302 | 2367 | 1878 |

**Table 6.** Loop count of every individual loop caller using KR normalized GM12878 dataset.

| Tools | 10KB (chr1) | 5KB (chr1) | 10KB (chr6) | 5KB (chr6) |
|---|---|---|---|---|
| LASCA | 1503 | 3146 | 1077 | 2095 |
| HiCExplorer | 1471 | 1757 | 1071 | 1319 |
| FitHiC2 | 63150 | 66440 | 46812 | 50260 |
| FitHiChIP | 3678 | 769 | 3165 | 951 |
| Peakachu | 7607 | 3867 | 2668 | 1027 |
| MUSTACHE | 5844 | 2809 | 3900 | 1946 |
| Chromosight | 958 | 1414 | 625 | 888 |
| SIP | 315 | 747 | 413 | 589 |
| HiCCUPS | 3890 | 3302 | 2367 | 1878 |

**Table 7.** Loop count of every individual loop caller using replicate GM12878 dataset.

| Tools | 10KB (chr1) | 5KB (chr1) | 10KB (chr6) | 5KB (chr6) |
|---|---|---|---|---|
| LASCA | 1543 | 3249 | 1045 | 2197 |
| HiCExplorer | 1732 | 1923 | 1119 | 1344 |
| FitHiC2 | 50638 | 67576 | 36962 | 53178 |
| FitHiChIP | 1864 | 225 | 1669 | 380 |
| Peakachu | 2638 | 905 | 855 | 281 |
| MUSTACHE | 4805 | 2237 | 3393 | 1563 |
| Chromosight | 928 | 1184 | 607 | 788 |
| SIP | 361 | 608 | 421 | 403 |
| cloops | 1658 | | 1140 | |
| cloops2 | 2492 | | 1696 | |
| HiCCUPS | 2930 | 1945 | 1586 | 952 |

**Table 8.** Average loop count of every individual loop caller using primary GM12878 dataset.

| Tools | Average |
|---|---|
| LASCA | 1897 |
| HiCExplorer | 1582 |
| FitHiC2 | 55140 |
| FitHiChIP | 1774 |
| Peakachu | 2288 |
| MUSTACHE | 3027 |
| Chromosight | 937 |
| SIP | 575 |
| cloops | 747 |
| cloops2 | 1082 |
| HiCCUPS | 2524 |

**Table 9.** Average loop size in KB.

| Tools | 5KB (chr1) | 10KB (chr1) | 5KB (chr6) | 10KB (chr6) |
|---|---|---|---|---|
| LASCA | 87317 | 160901 | 88294 | 169335 |
| HiCExplorer | 326665 | 736447 | 329744 | 760818 |
| FitHiC2 | 1351696 | 1114492 | 131833 | 76612 |
| FitHiChIP | 226494 | 287834 | 461077 | 407102 |
| Peakachu | 72711 | 134957 | 80184 | 176604 |
| MUSTACHE | 356814 | 513525 | 392389 | 556635 |
| Chromosight | 102752 | 123386 | 101524 | 126205 |
| SIP | 352191 | 435181 | 425981 | 536290 |
| cloops | 78584 | 78584 | 182691 | 182691 |
| cloops2 | 308986 | 308986 | 338860 | 338860 |
| HiCCUPs | 146645 | 219439 | 168374 | 258051 |

**Table 10.** Average loop size in terms of number of bin.

| Tools | 5KB (chr1) | 10KB (chr1) | 5KB (chr6) | 10KB (chr6) |
|---|---|---|---|---|
| LASCA | 17 | 16 | 18 | 17 |
| HiCExplorer | 65 | 74 | 66 | 76 |
| FitHiC2 | 270 | 111 | 26 | 8 |
| FitHiChIP | 45 | 29 | 92 | 41 |
| Peakachu | 15 | 13 | 16 | 18 |
| MUSTACHE | 71 | 51 | 78 | 56 |
| Chromosight | 21 | 12 | 20 | 13 |
| SIP | 70 | 44 | 85 | 54 |
| cloops | 16 | 8 | 37 | 19 |
| cloops2 | 62 | 31 | 68 | 34 |
| HiCCUPs | 29 | 22 | 34 | 26 |

**Table 11.** APA score using GM12878 primary data.

| Tools | 5KB (chr1) | 10KB (chr1) | 5KB (chr6) | 10KB (chr6) |
|---|---|---|---|---|
| LASCA | 1 | 1.2 | 0.953 | 1.21 |
| HiCExplorer | 2.87 | 2.28 | 2.98 | 2.27 |
| FitHiC2 | 179 | 83.1 | 28.1 | 18.6 |
| FitHiChIP | 2.52 | 1.75 | 4.89 | 1.89 |
| Peakachu | 1.33 | 1.3 | 1.34 | 1.55 |
| MUSTACHE | 1.4 | 1.56 | 1.46 | 1.57 |
| Chromosight | 1.11 | 0.904 | 1.13 | 0.91 |
| SIP | 3.4 | 3.59 | 3.7 | 3.33 |
| cloops | 2.73 | | 5.12 | |
| cloops2 | 3.61 | | 3.98 | |
| HiCCUPs | 2.04 | 1.81 | 2.1 | 1.82 |

**Table 12.** APA score using GM12878 KR normalized data.

| Tools | 5KB (chr1) | 10KB (chr1) | 5KB (chr6) | 10KB (chr6) |
|---|---|---|---|---|
| LASCA | 1.05 | 1.13 | 1.1 | 1.3 |
| HiCExplorer | 3.45 | 1.72 | 1.67 | 1.27 |
| FitHiC2 | 212 | 76.5 | 36.9 | 16.7 |
| FitHiChIP | 2.54 | 1.68 | 4.86 | 1.92 |
| Peakachu | 1.13 | 0.933 | 1.26 | 1.04 |
| MUSTACHE | 1.45 | 1.39 | 1.52 | 1.44 |
| Chromosight | 1.18 | 0.903 | 1.19 | 0.902 |
| SIP | 2.7 | 3.03 | 2.72 | 2.77 |
| HiCCUPs | 2.08 | 1.78 | 2.12 | 1.86 |

**Table 13.** APA score using GM12878 replicate data.

| Tools | 5KB (chr1) | 10KB (chr1) | 5KB (chr6) | 10KB (chr6) |
|---|---|---|---|---|
| LASCA | 1.1 | 1.17 | 1.05 | 1.21 |
| HiCExplorer | 2.64 | 1.72 | 2.75 | 1.97 |
| FitHiC2 | 174 | 88.6 | 67.8 | 23.4 |
| FitHiChIP | 2.8 | 1.91 | 9.43 | 2.34 |
| Peakachu | 1.24 | 1.05 | 1.28 | 1.15 |
| MUSTACHE | 1.36 | 1.37 | 1.39 | 1.4 |
| Chromosight | 1.12 | 0.907 | 1.12 | 0.849 |
| SIP | 2.36 | 2.57 | 2.65 | 2.45 |
| cloops | 2.28 | | 2.52 | |
| cloops2 | 2.25 | | 2.11 | |
| HiCCUPs | 2.08 | 1.72 | 2.19 | 1.82 |

### 3 Supplemental Figures

#### 3.1 Loop Count

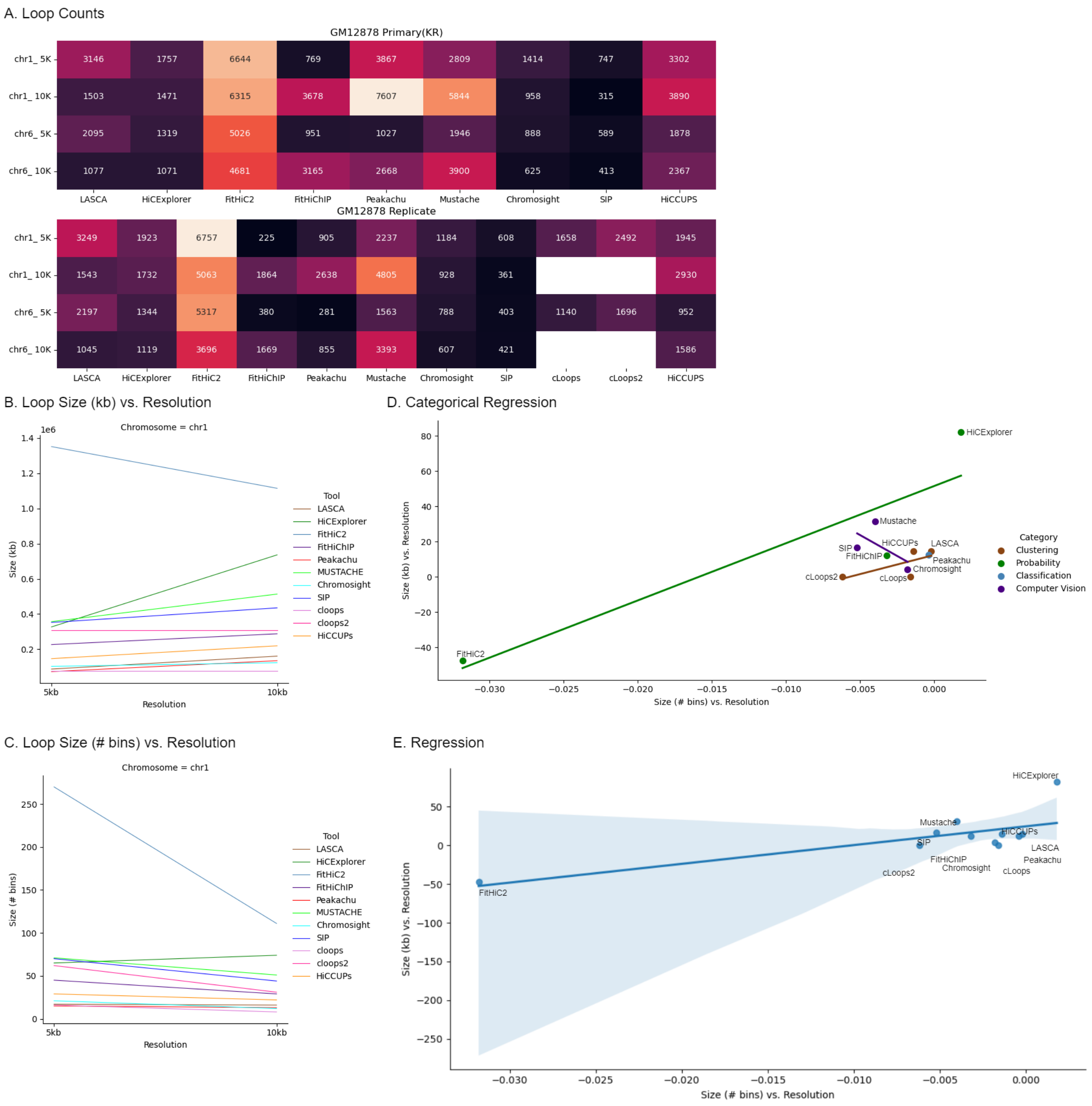


**Figure 1.** A. Chromatin loop counts for KR normalized GM12878 cell at the top and replicate data at the bottom. For all combinations, FitHiC2 predicts the highest number of loops and we represented this tools loop count multiple of 1/10. Apart from that, Peakachu predicts the highest (7607) number of loop and SIP finds the lowest (315) number of loops at 10KB resolution from chromosome 1, and at 5KB resolution, Peakachu (3867) finds the highest and SIP (747) finds the lowest. Using chromosome 6, Mustache (3900) predicts the highest number of loops and SIP (413) predicts lowest number of loops at 10KB resolution, and at 5KB resolution, LASCA (2095) predicts the most amount loops and SIP (589) predicts the least amount of loops. For chromosome 1 at 5KB resolution, LASCA (3146), Mustache (2809) and HiCCUPS (3302) predicts significant amount of loops, and at 10KB resolution, FitHiChIP (3678), Mustache (5844) and HiCCUPS (3890) predicts significant loops. Considering chromosome 6 at 5KB, Mustache (1946), HiCCUPS (1878), HiCExplorer (1319), Peakachu (1027) predicts a shear amount of loops, and at 10KB resolution, FitHiChIP (3165), Peakachu (2668), HiCCUPS (2367) predicts a considerable amount of loops. Again, we computed loop count using replicate data. Mustache predicts 4805 loops at 10KB using chromosome 1 and SIP predicts 361 loops. For chromosome 6 at 10KB resolution, Mustache predicts 3393 loops and SIP predicts 403 loops at 5KB resolution. cLoops and cLoops2 do not associate with resolution, rather they produce chromosome wise loops. They accepts bedpe format data and does not have an option to analyze normalize data.

##### 3.2 Overlap Analysis

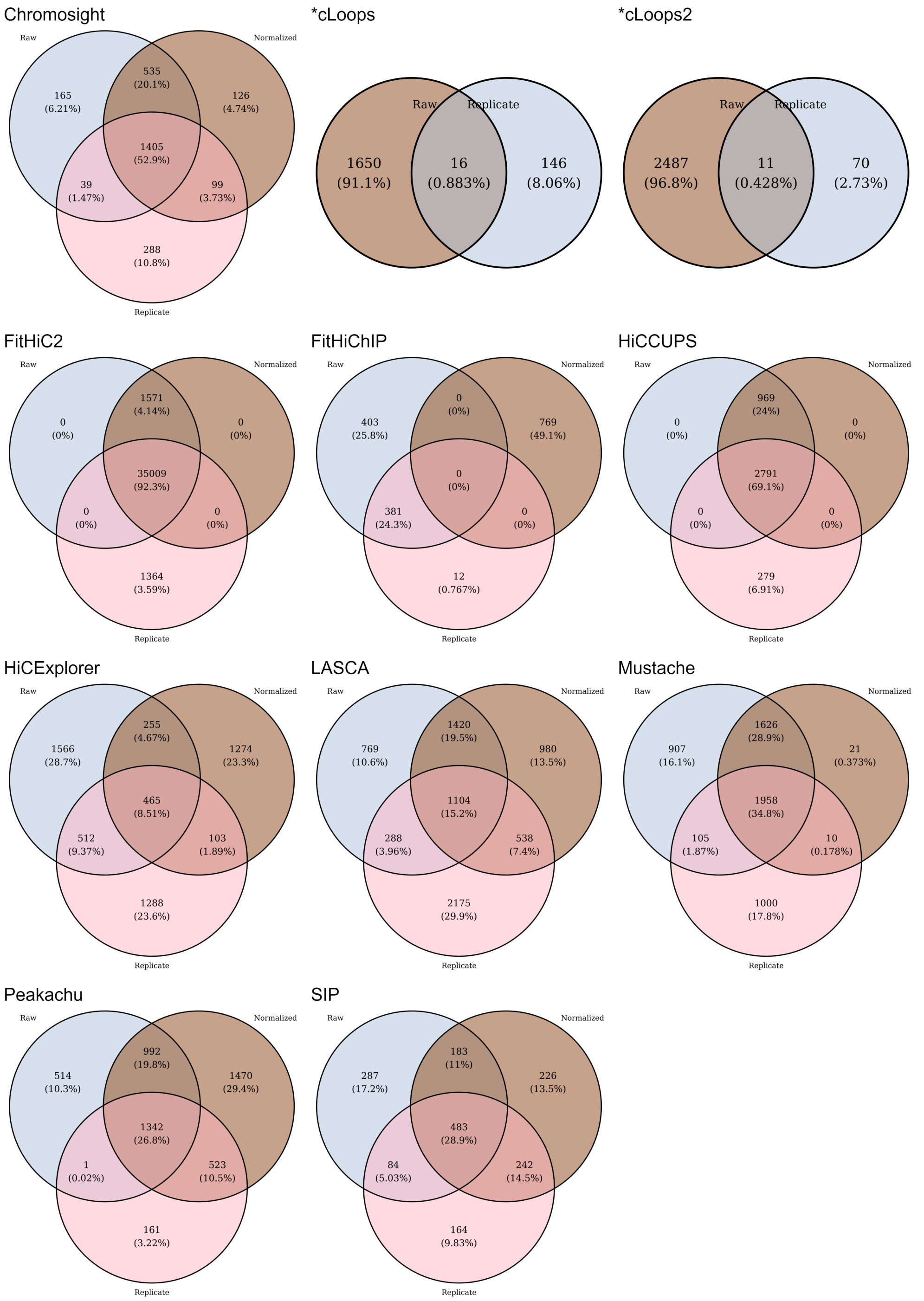


**Figure 2.** Overlap across three (primary, replicate and KR normalized) dataset of GM12878 using chromosome 1 at 5kb resolution. As FitHiC2 predicts a highest amount of loops, it overlap ratio is highest. Considering other tools, HiCCUPS scores 69.1%, Chromosight 52.9%, Mustache 34.8% and FitHiChIP does not overlap any loops. Other tools overlaps below 30% loops. cLoops and cLoops overlaps near 0% loops and HiCExplorer scores 8.51%.

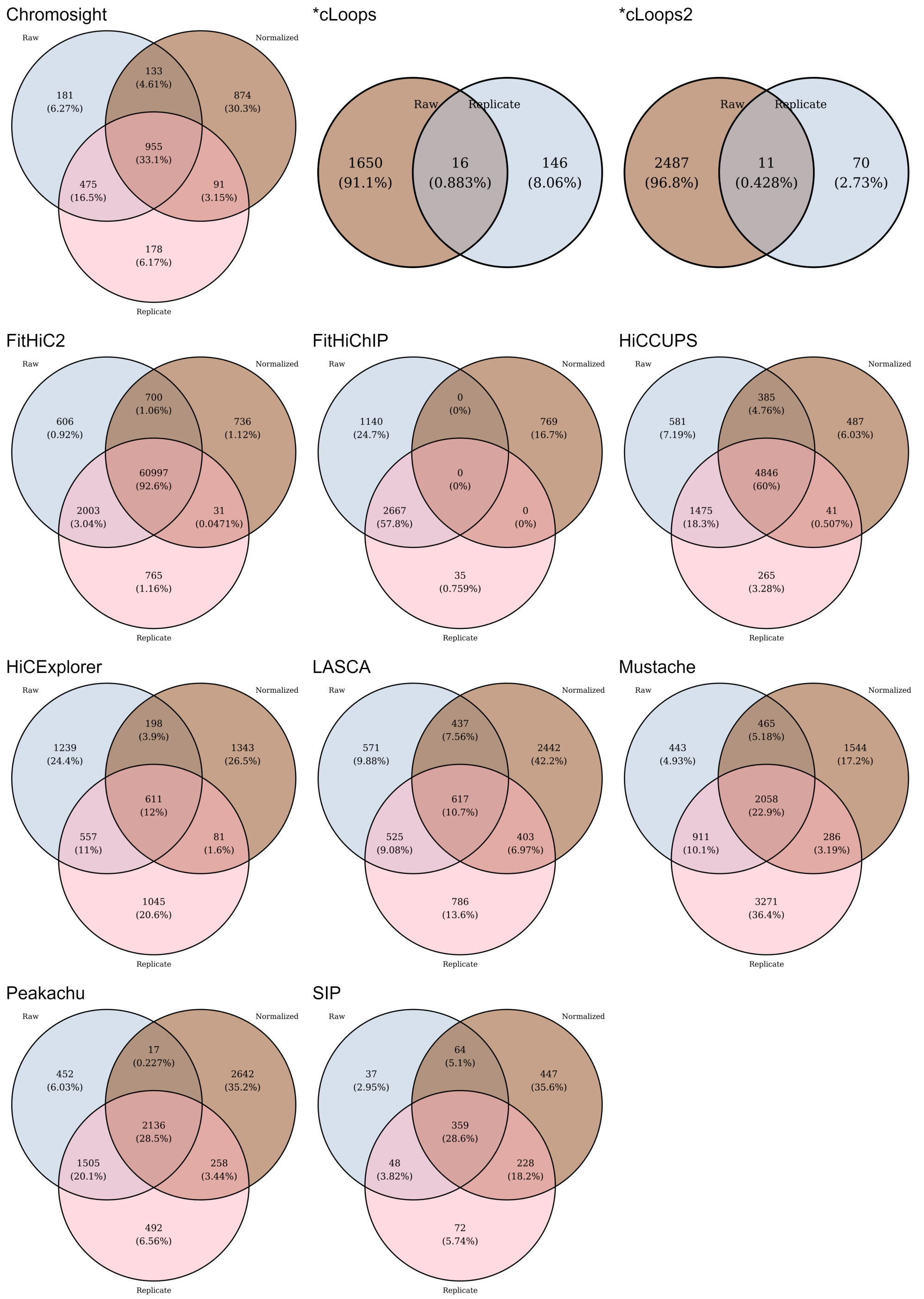


**Figure 3.** Overlap across three (primary, replicate and KR normalized) dataset of GM12878 using chromosome 1 at 10kb resolution. FitHiC2 overlaps 92.6% loops across three datasets. cLoops and cLoops2 scores near to zero and FitHiChIP does not overlap any loops. HiCExplorer, LASCA, Mustache, Peakachu and SIP overlap below 30% loops. HiCCUPS overlaps 60% loops and Chromosight scores around 33%.

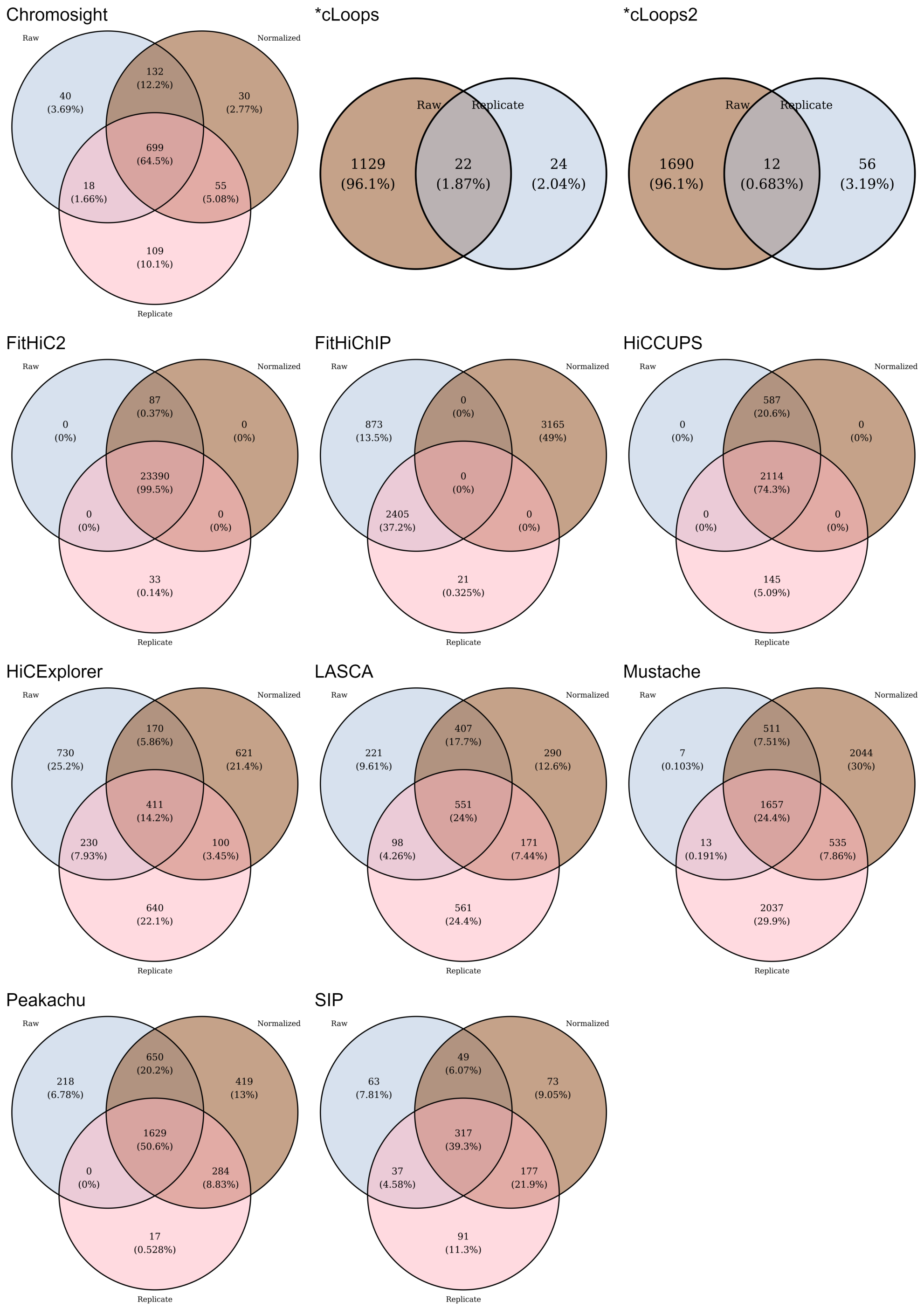


**Figure 4.** Overlap across three (primary, replicate and KR normalized) dataset of GM12878 using chromosome 6 at 10kb resolution. FitHiC2 overlaps allmost all the loops, and Chromosight overlaps 64.5%, Peakachu overlaps 50.6% and HiCCUPS overlaps 74.3% loops. SIP scores around 40%, and HiCExplorer, LASCA and Mustache overlaps below 30% loops. cLoops and cLoops2 overlaps around 1% loops across primary and replicate data.

##### 3.3 Peak Analysis

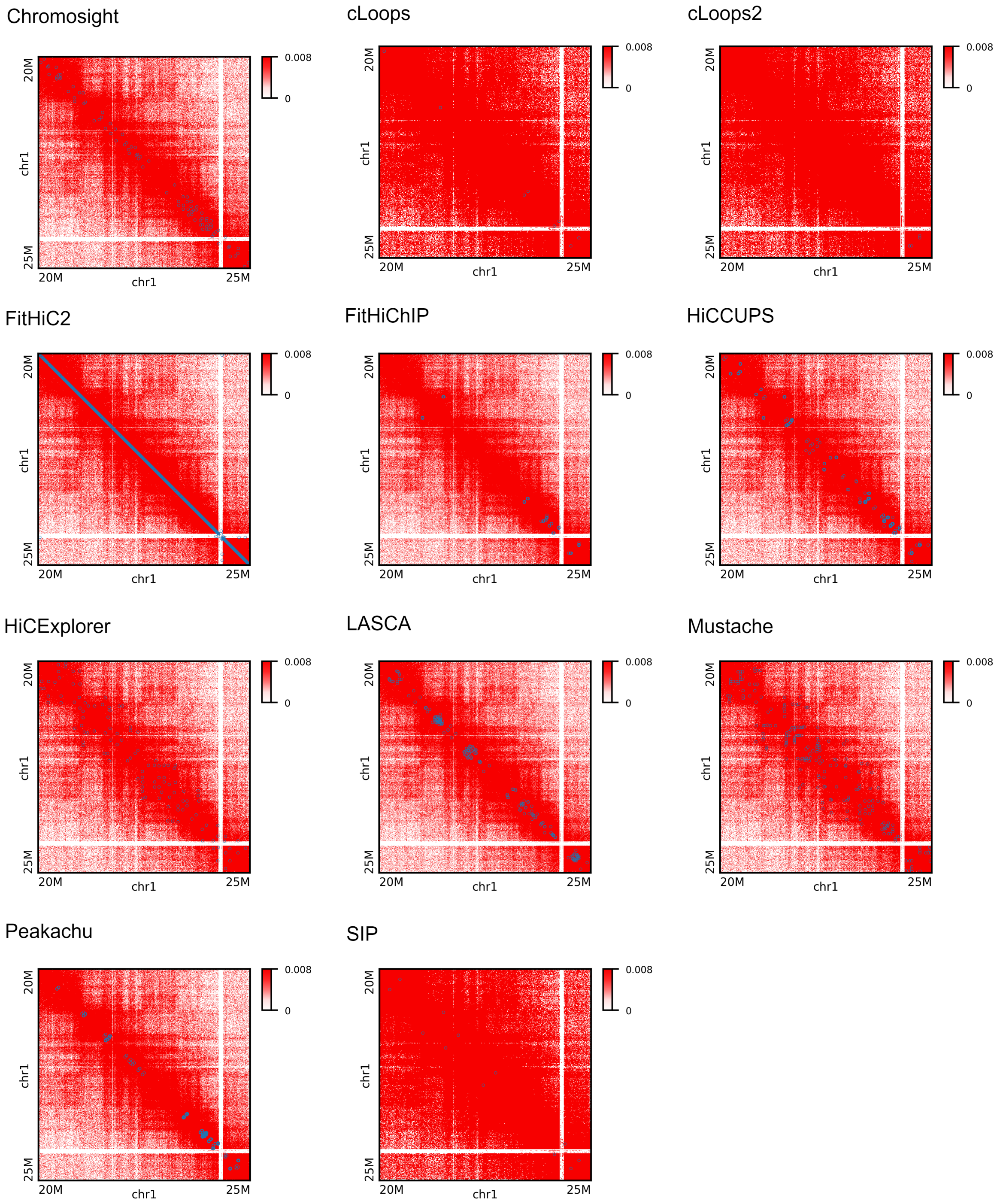


**Figure 5.** Peak plots for chromosome 1 at 5KB using primary GM12878 data from 20M to 25M genomic range. FitHiC2 marks peaks through the diagonal from left to right creating a diagonal strong straight line. Chromosight, HiCCUPS, HiCExplorer, LASCA, Mustache, and Peakachu mark significant number of the peaks in this region. cLoops, cLoops2, FitHiChIP, and SIP finds least amount of peaks in this region.

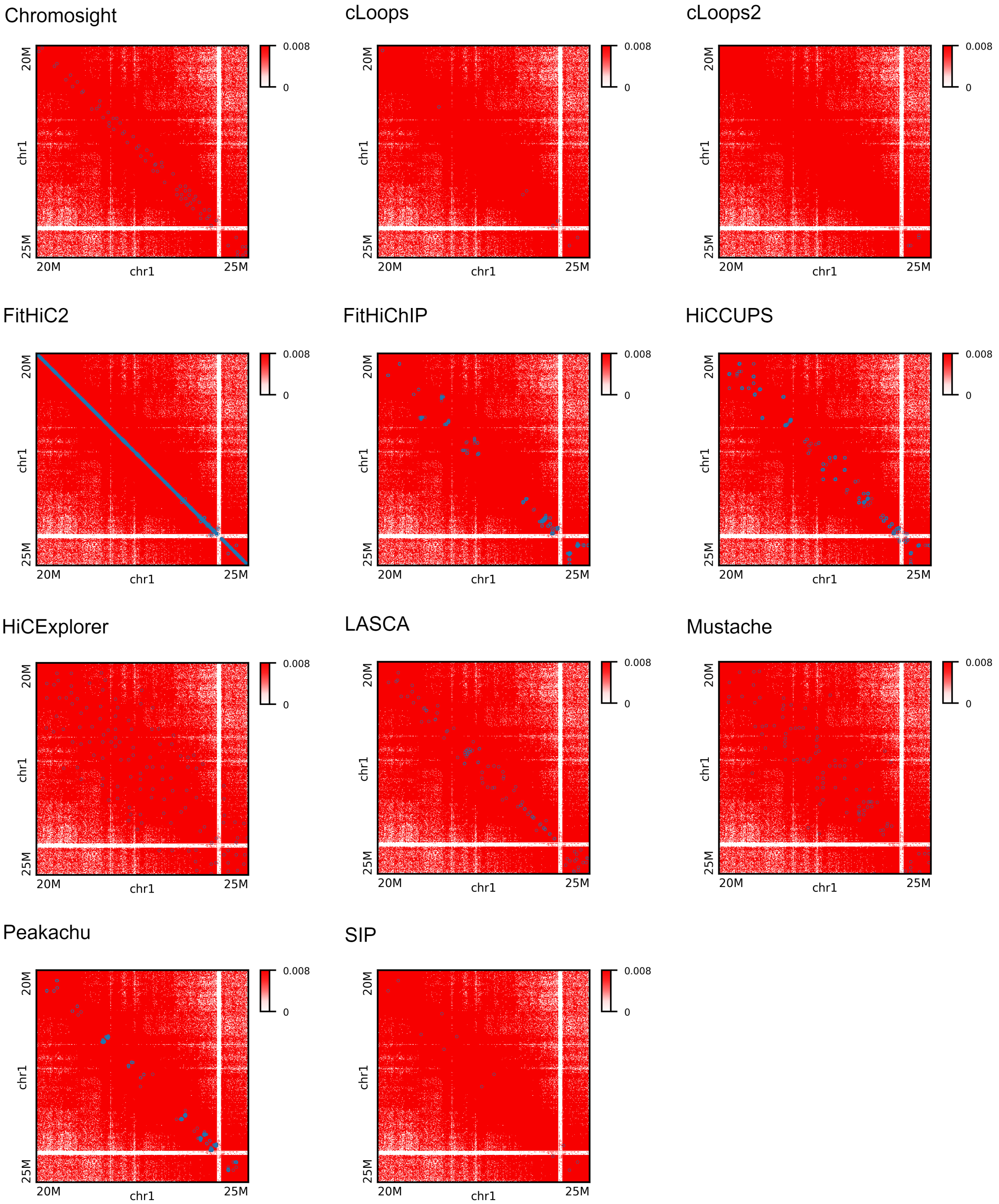


**Figure 6.** Peak plots for chromosome 1 at 10KB using primary GM12878 data from 20M to 25M genomic range. SIP, cLoops and cLoops2 finds the least amount of peaks whereas FitHiChIP, Chromosight, LASCA and Peakachu more dense compared to them. HiCExplorer and Mustache mark peaks covering a wide range of area, and HiCCUPS points significant amount of peaks. FitHiChIP marks more peaks compared to 5KB data. HiCExplorer marks peaks within a large diagonal region in the heatmap compared to 5KB data. FitHiC2 marks peaks through the diagonal from left to right creating a diagonal strong straight line.

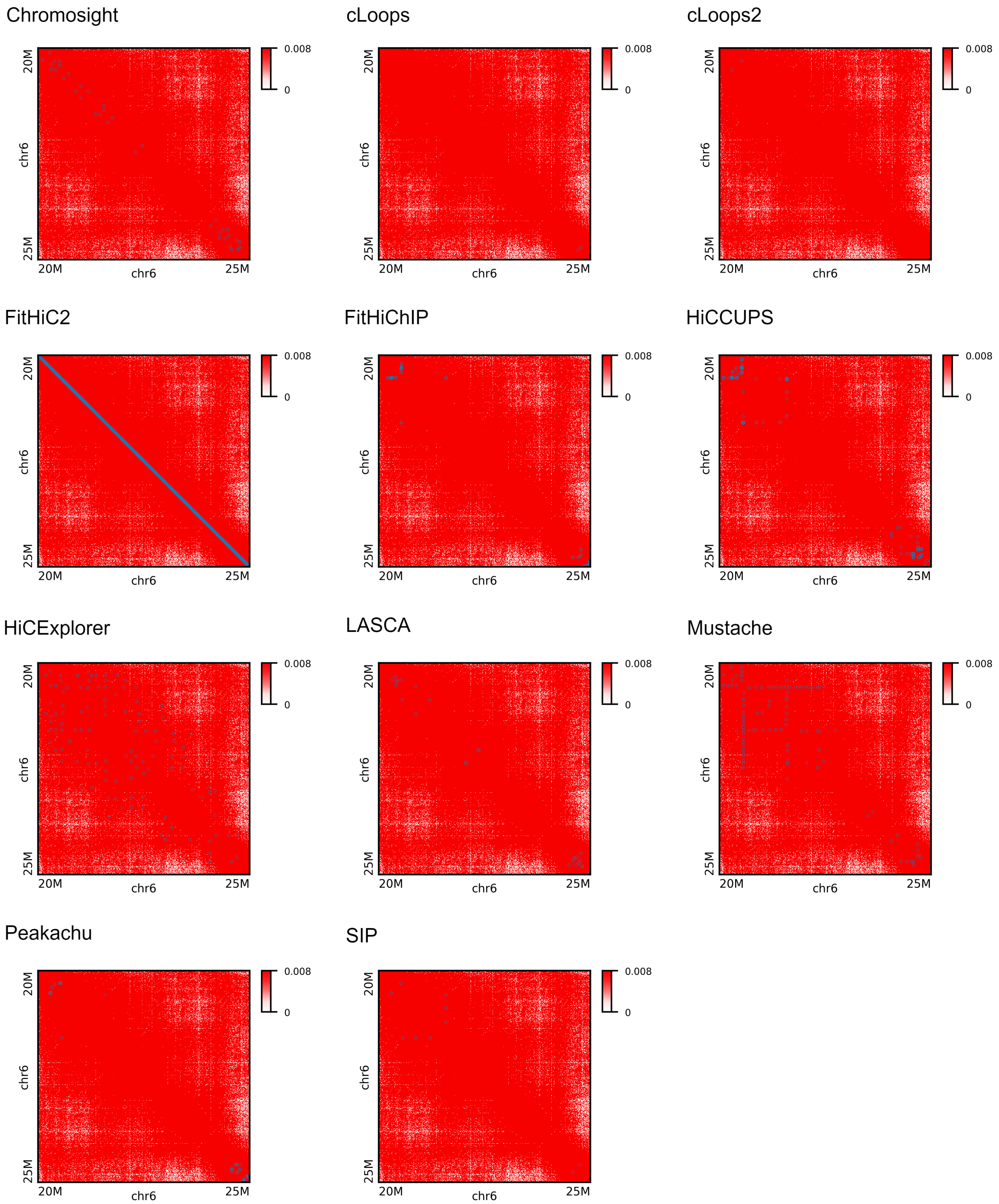


**Figure 7.** Peak plots for chromosome 6 at 10KB using primary GM12878 data from 20M to 25M genomic range. cLoops and cLoops2 marks almost zero amount of peaks. FitHiC2 marks peaks through the diagonal from left to right creating a diagonal strong straight line. HiCCUPS, Mustache and SIP mark peaks in the upper left diagonal corner. HiCExplorer and Mustache covers a wide area predicting peaks. HiCExplorer, Mustache, Chromosighr, and HiCCUPS mark more peaks compared to other tools at this region.

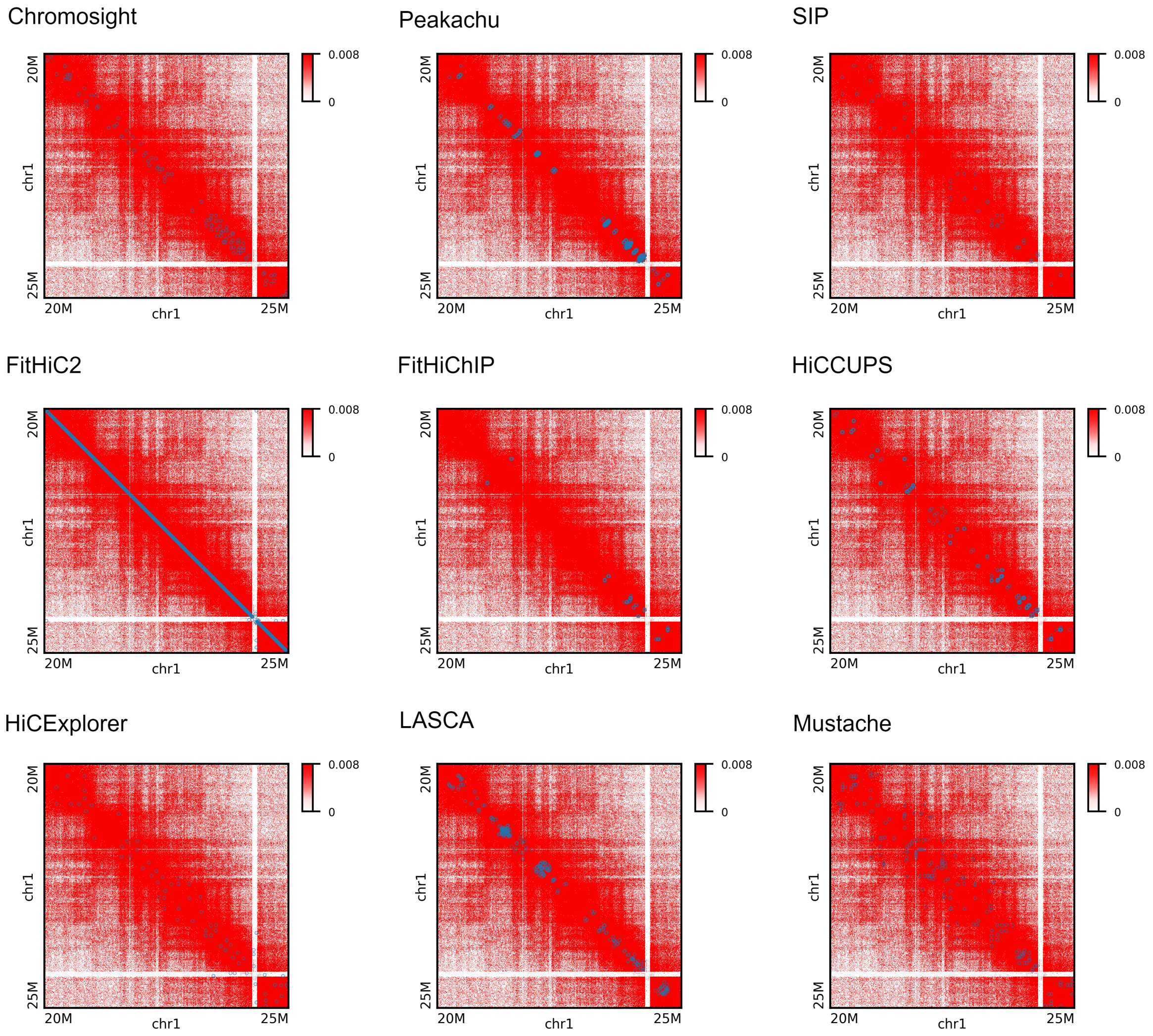


**Figure 8.** Peak plots for chromosome 1 at 5KB using KR normalized GM12878 data from 20M to 25M genomic range. FitHiChIP marks less peaks compared to other tools, and LASCA and Peakachu marks saturated loops at some points. FitHiC2 marks peaks through the diagonal from left to right creating a diagonal strong straight line. Mustache, LASCA, HiCCUPS, Peakachu, Chromosight and HiCExplorer marks a visible amount of peaks. Mustache marks peaks within a wide range of diameter from the left to right diagonal line.

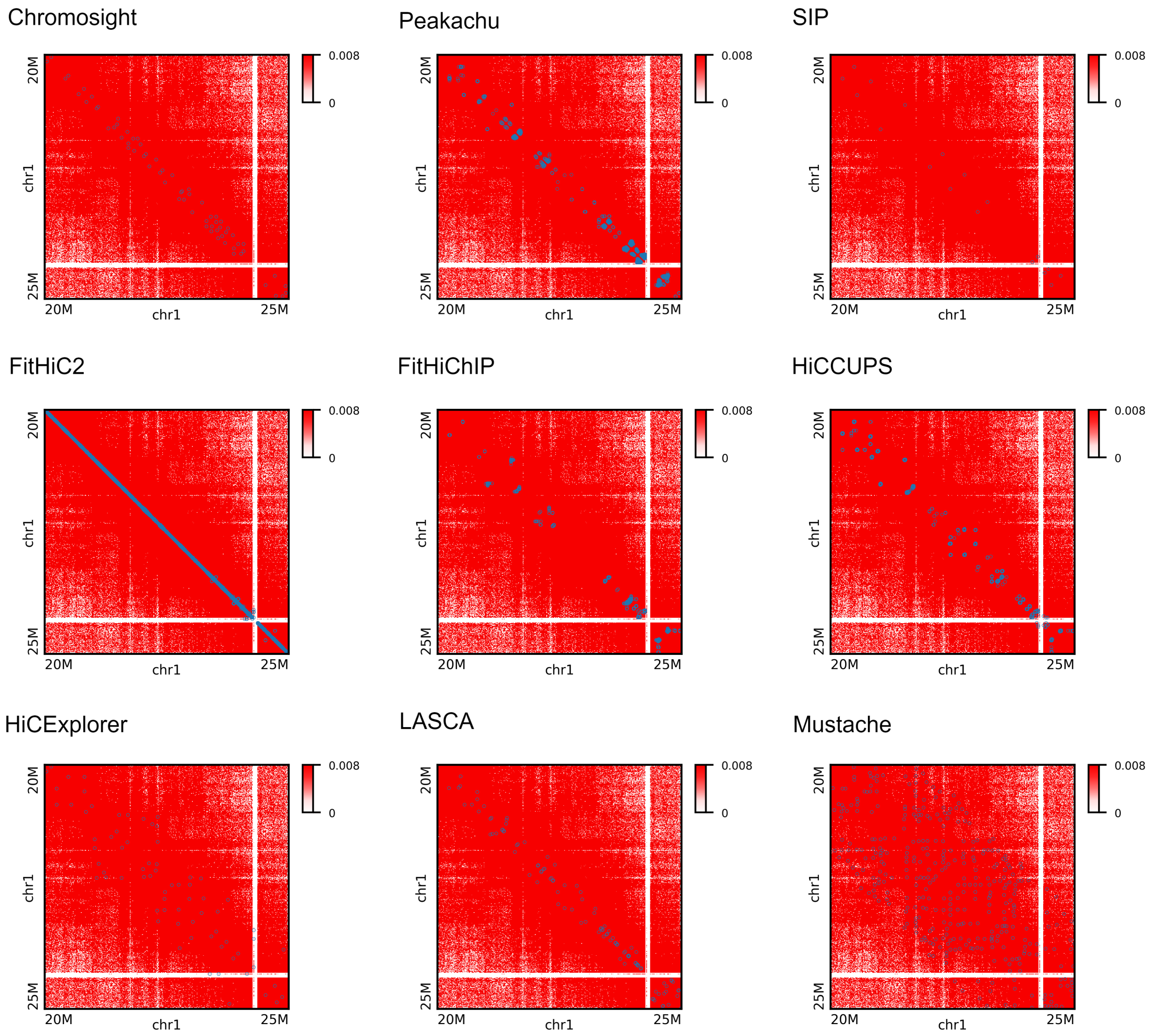


**Figure 9.** Peak plots for chromosome 1 at 10KB using KR normalized GM12878 data from 20M to 25M genomic range. Mustache marks peaks over a diagonal region covering most of the region. SIP marks less peaks, and Peakachu, FitHiChIP, HiCCUPS and LASCA create some saturated peaks points.

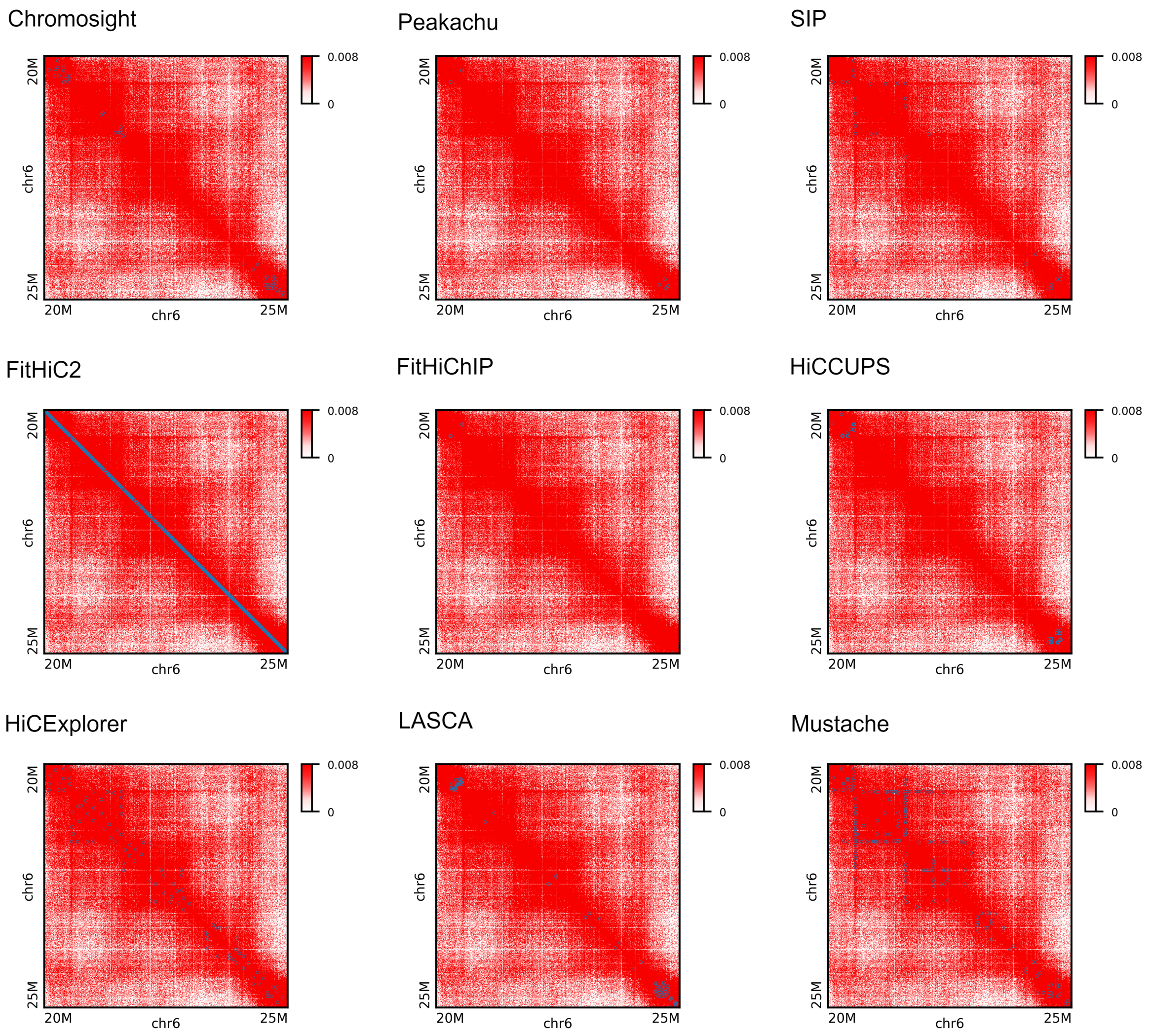


**Figure 10.** Peak plots for chromosome 6 at 5KB using KR normalized GM12878 data from 20M to 25M genomic range. Mustache and HiCExplorer marks more loops compared to other tools.

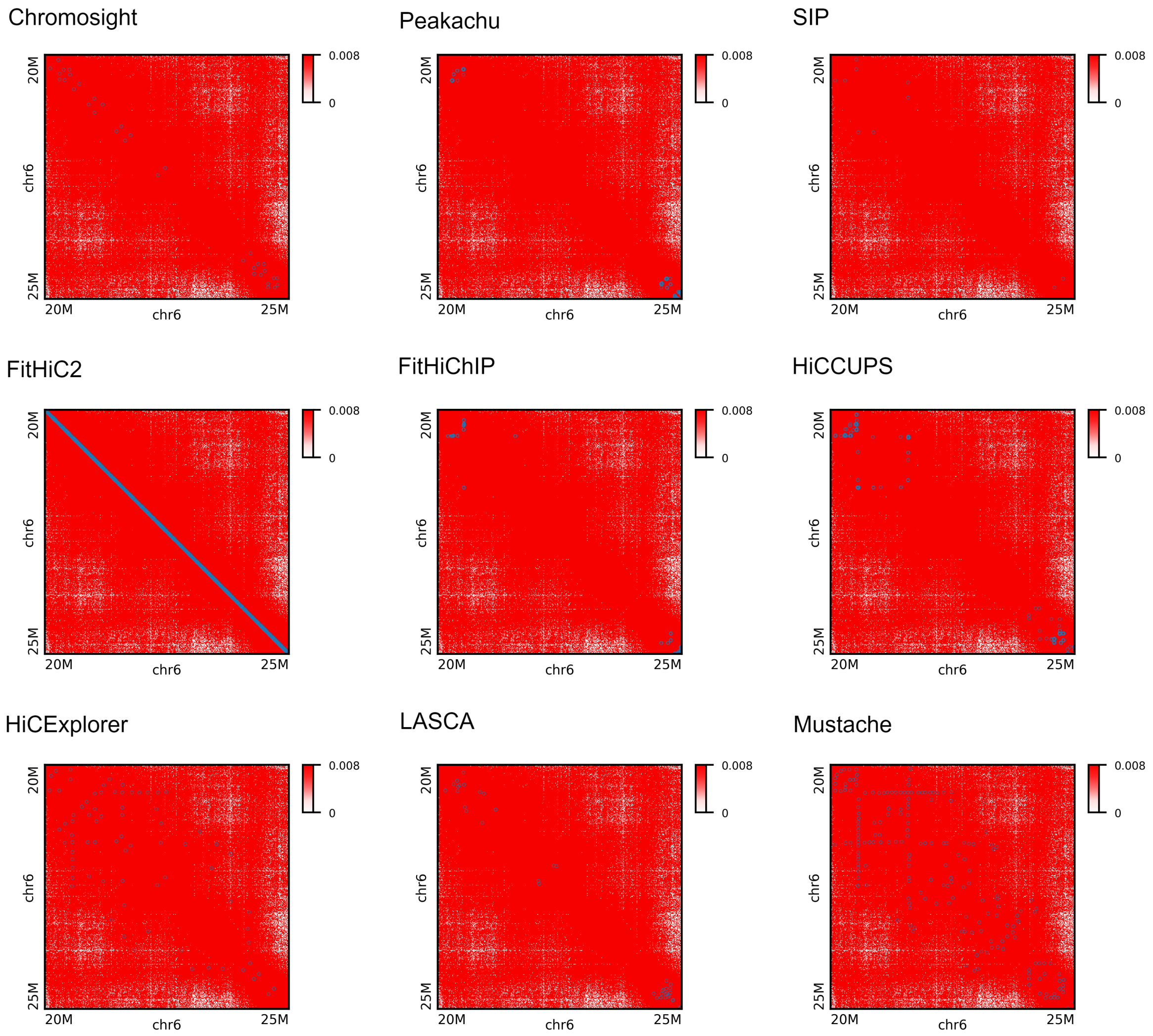


**Figure 11.** Peak plots for chromosome 6 at 10KB using KR normalized GM12878 data from 20M to 25M genomic range. Mustache continues the same pattern as 5KB with some less peaks marked where HiCExplorer covers a large diagonal area. HiCCUPS marks more loops compared to 5KB data.

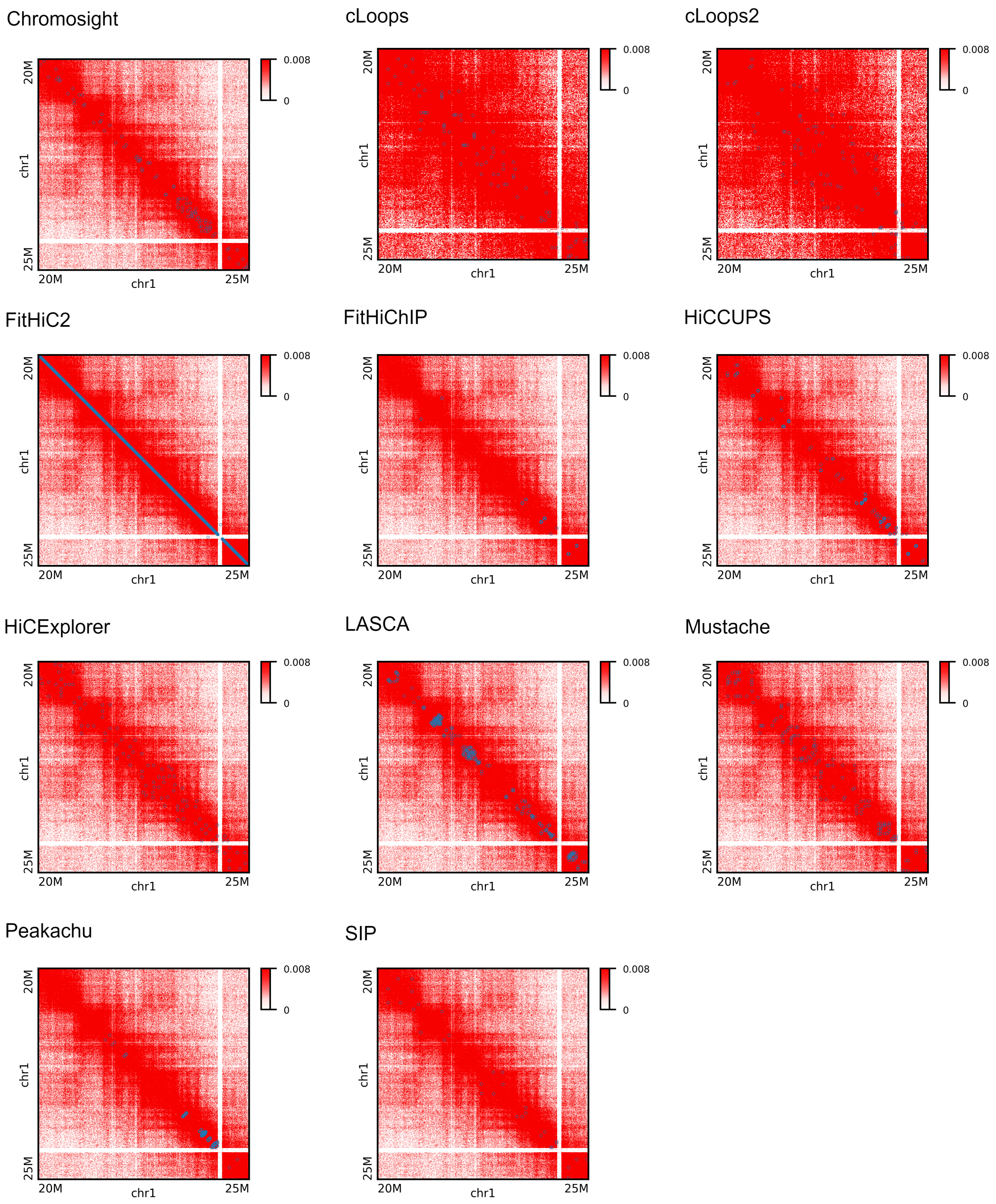


**Figure 12.** Peak plots for chromosome 1 at 5KB using replicate GM12878 data from 20M to 25M genomic range. FitHiChIP, Peakachu, and SIP mark the least amount of peaks at this region.

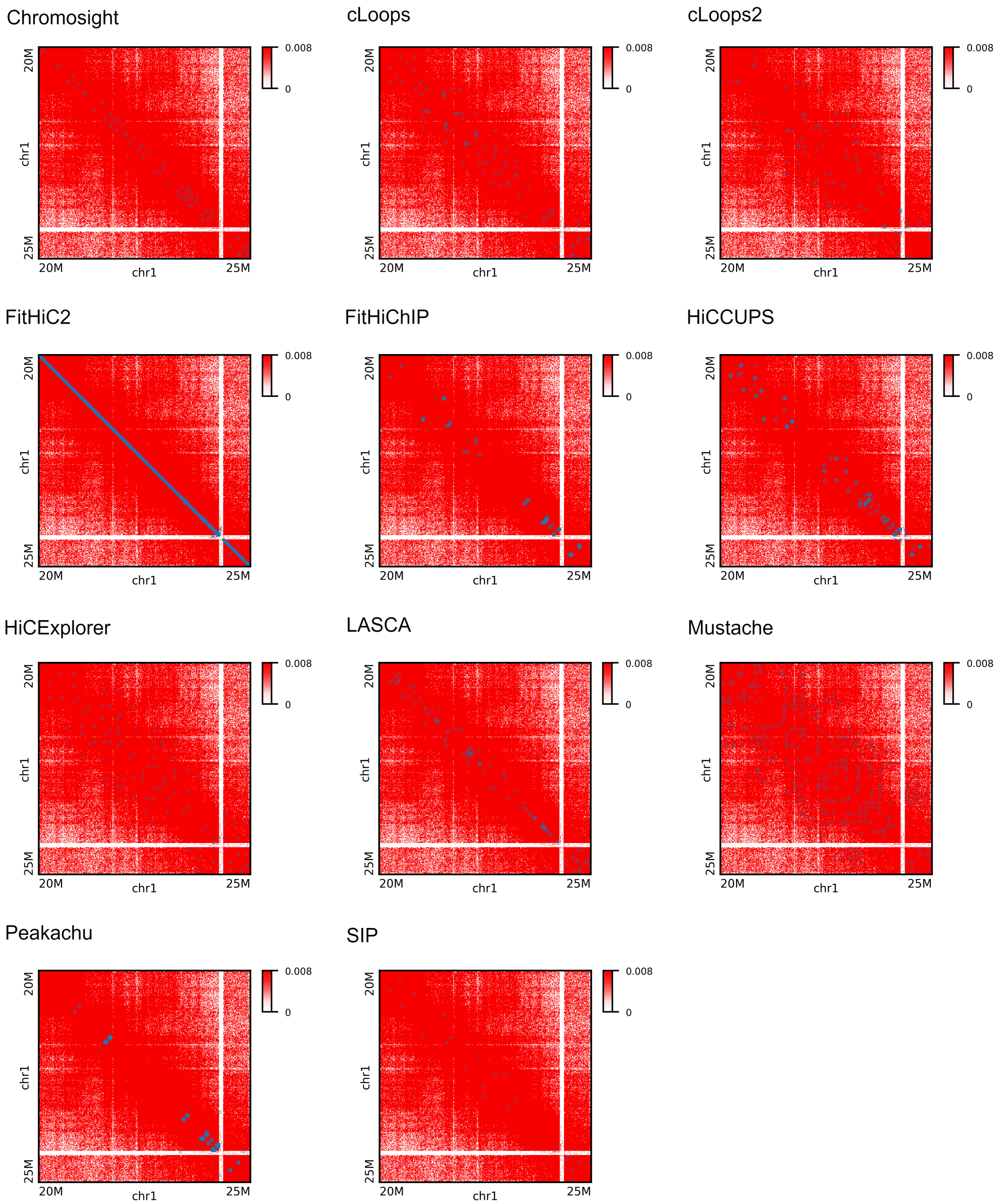


**Figure 13.** Peak plots for chromosome 1 at 10KB using replicate GM12878 data from 20M to 25M genomic range. Mustache and HiCExplorer covers a large area diagonally whrer as Chromosight, cLoops, and cLoops continue the same pattern as 5KB. FitHiChIP shows some saturated peaks at some points.

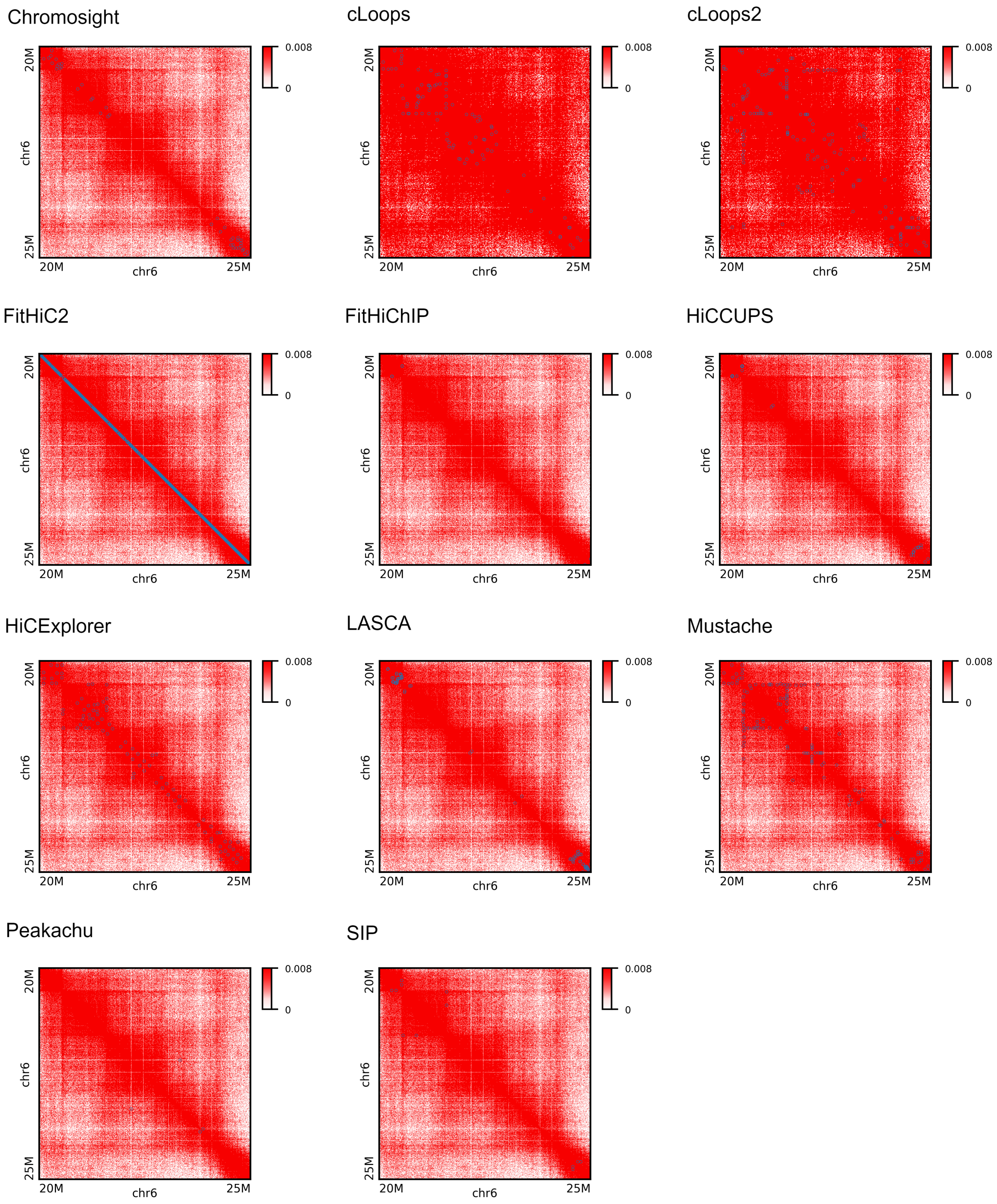


**Figure 14.** Peak plots for chromosome 6 at 5KB using replicate GM12878 data from 20M to 25M genomic range. cLoops, cLoops2, HiCExplorer, and Mustache marks most of the peaks at this regions. LASCA shows saturation points at corner sides.

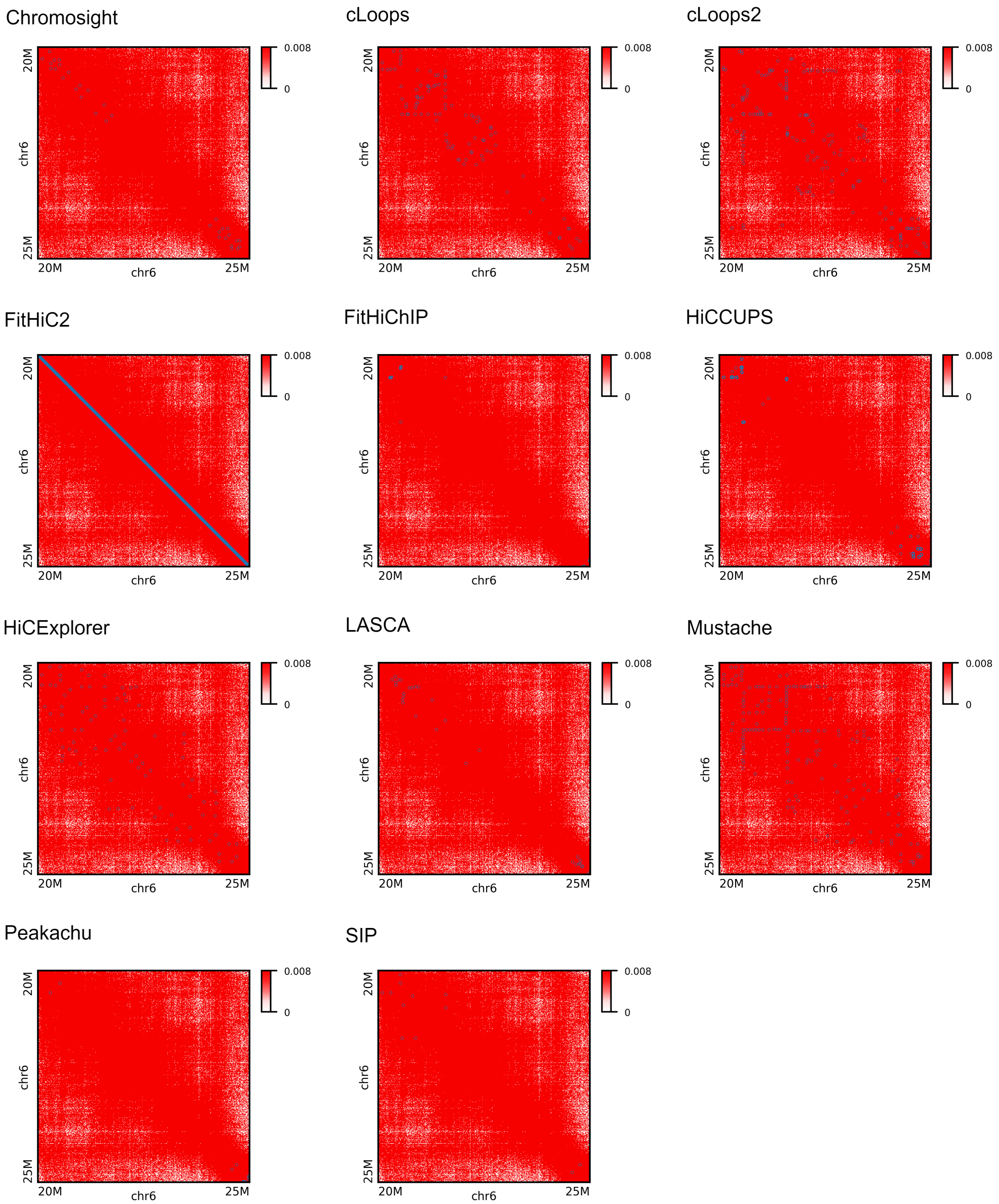


**Figure 15.** Peak plots for chromosome 6 at 10KB using replicate GM12878 data from 20M to 25M genomic range. HiCCUPS marks more peaks at 10KB compared to 5KB resolution data. HiCExplorer, Mustache, and cLoops2 cover a large area through the diagonal line. cLoops2 shows almost same peaks as in 5KB data.

##### 3.4 APA Analysis

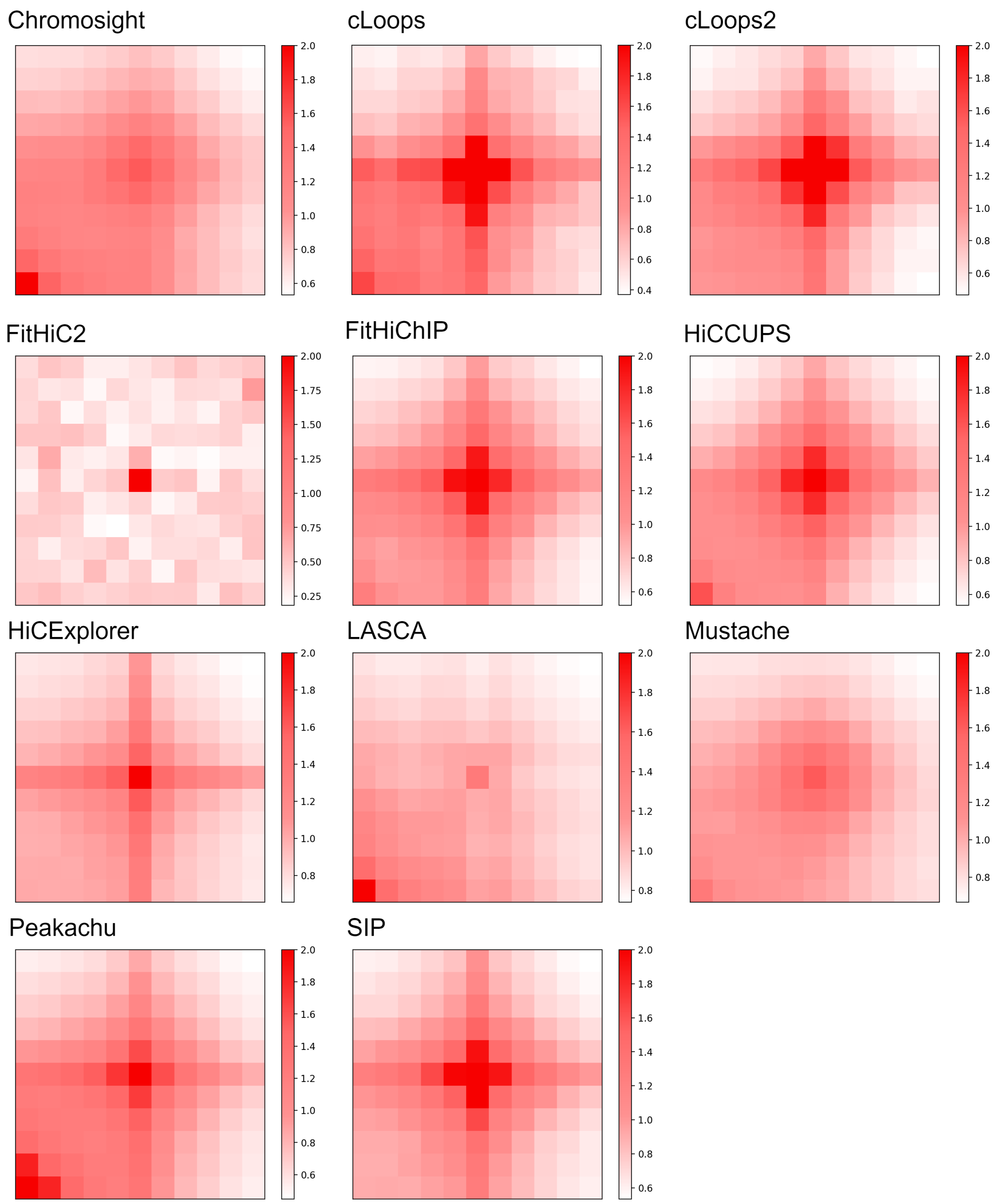


**Figure 16.** APA plot for chromosome 1 at 5KB using GM12878 primary data. FitHiC2 has the highest enrichment. cLoops, cLoops2, FitHiChIP, HiCCUPS, HiCExplorer, SIP have strong enrichment at the center. LASCA show strong enrichment at the lower left corner of the center.

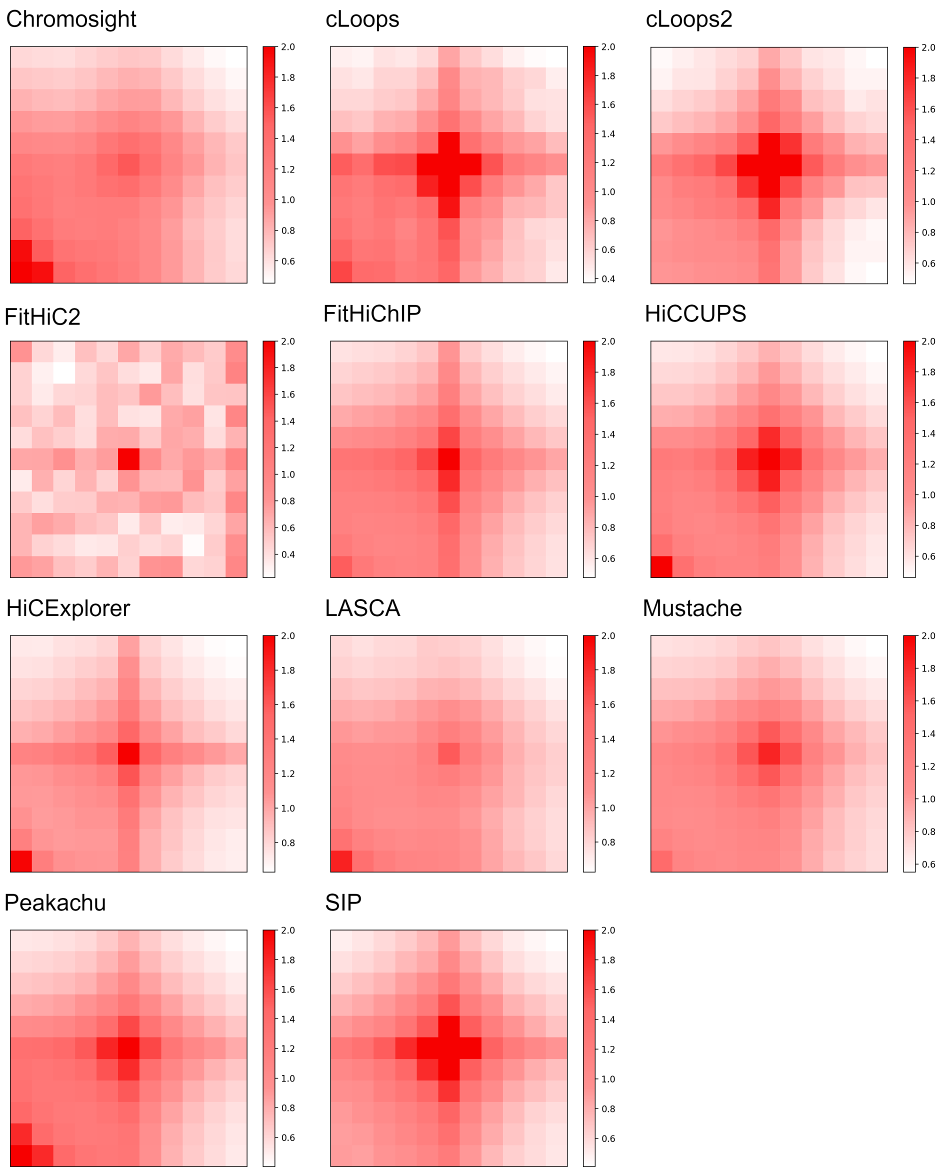


**Figure 17.** APA plot for chromosome 1 at 10KB using GM12878 primary data. FitHiC2 shows the highest APA score at the center, and cLoops, cLoops2, HiCExplorer, and SIP have strong enrichment at the center. Chromosight shows APA score less than 1 and other tools have enrichment greater than 1.

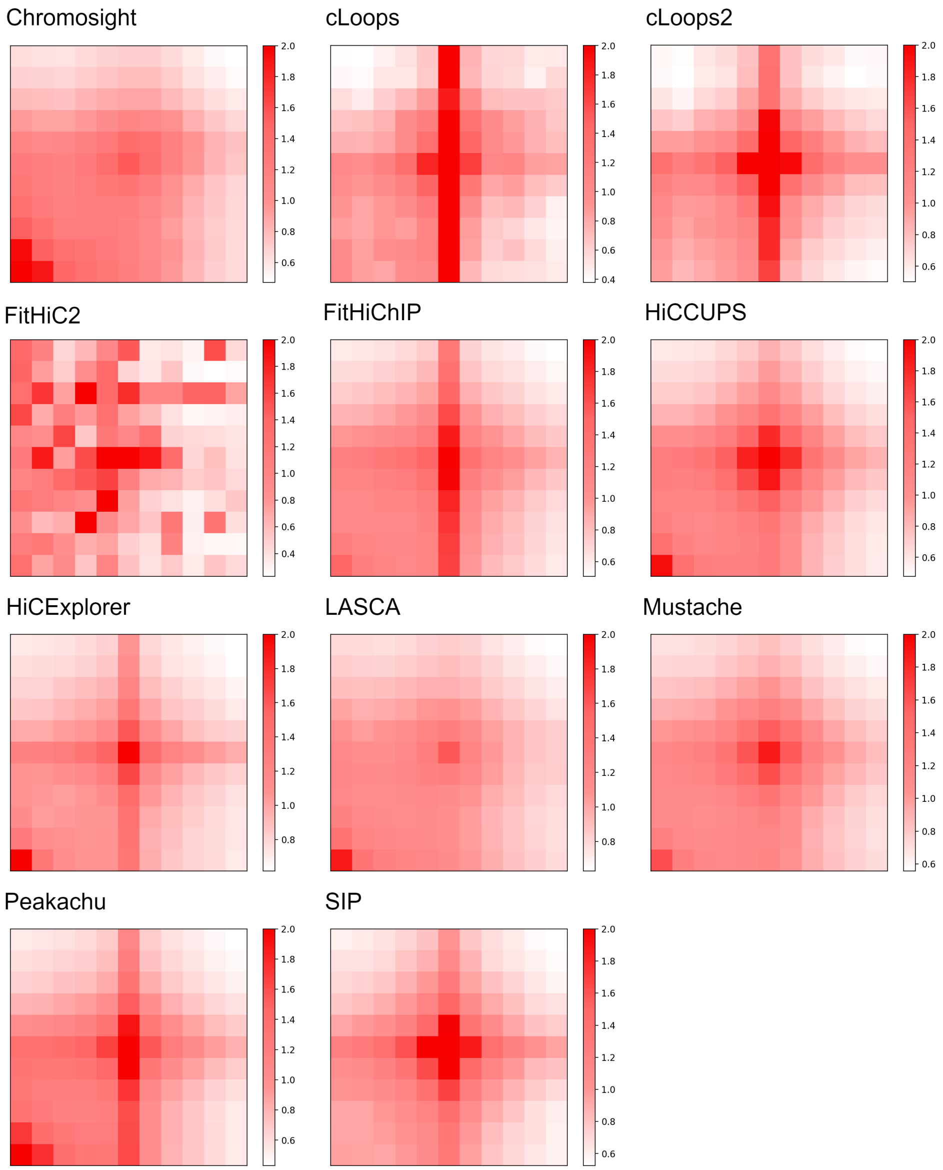


**Figure 18.** APA plot for chromosome 6 at 10KB using GM12878 primary data. cLoops, cLoops2, FitHiChIP, and Peakachu have strong enrichment vertically through the center. FitHiC2 shows highest enrichment in multiple focal points and LASCA produced lowest APA score. Chromosight, Peakachu, HiCCUPS, HiCExplorer, and LASCA show strong enrichment at the lower left corner of the center.

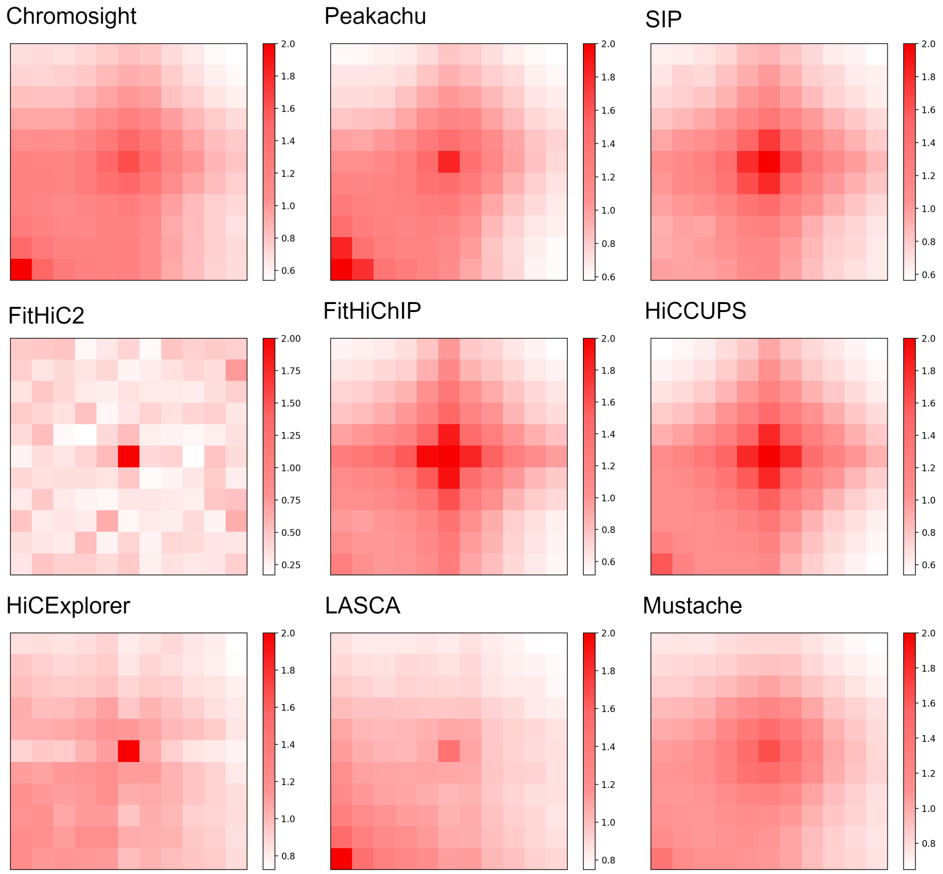


**Figure 19.** APA plot for chromosome 1 at 5KB using GM12878 KR normalized data. Though Chromosight, Peakachu, and LASCA have strong enrichment at the lower left corner, all the tools shows strong enrichment in the central focal point and score greater than 1.

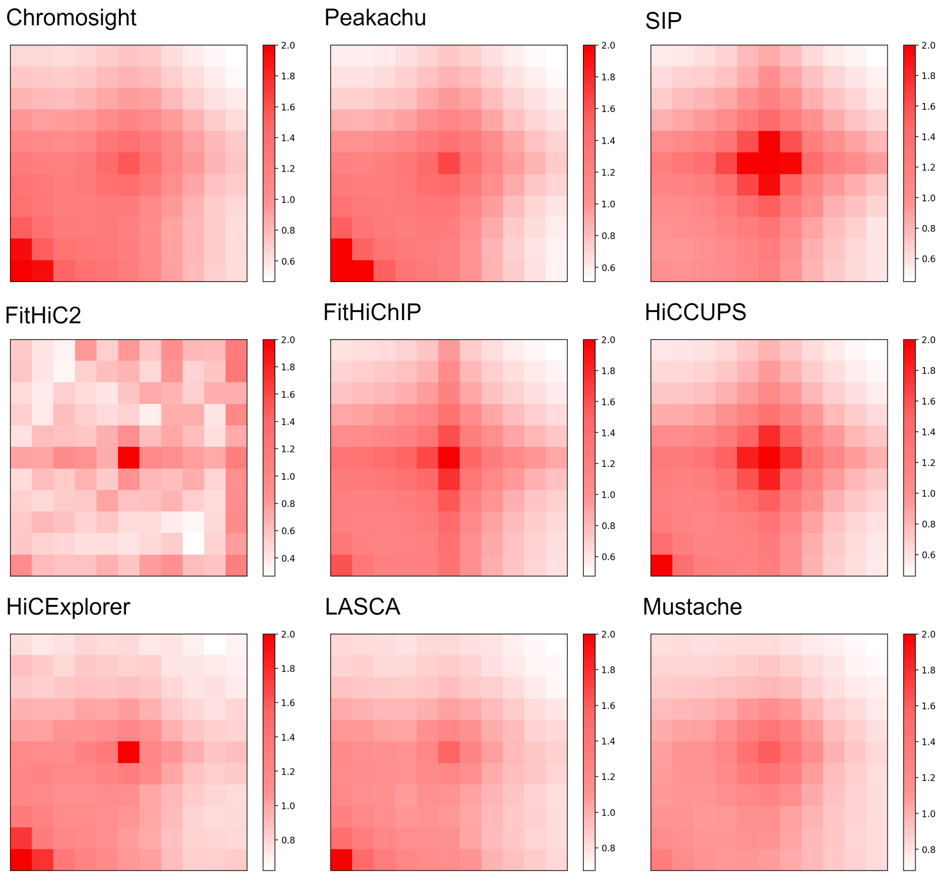


**Figure 20.** APA plot for chromosome 1 at 10KB using GM12878 KR normalized data. Here, all the tools have strong color in the center and dimmed symmetrically from the center except FitHiC2, and Peakachu and Chromosight score less than 1.

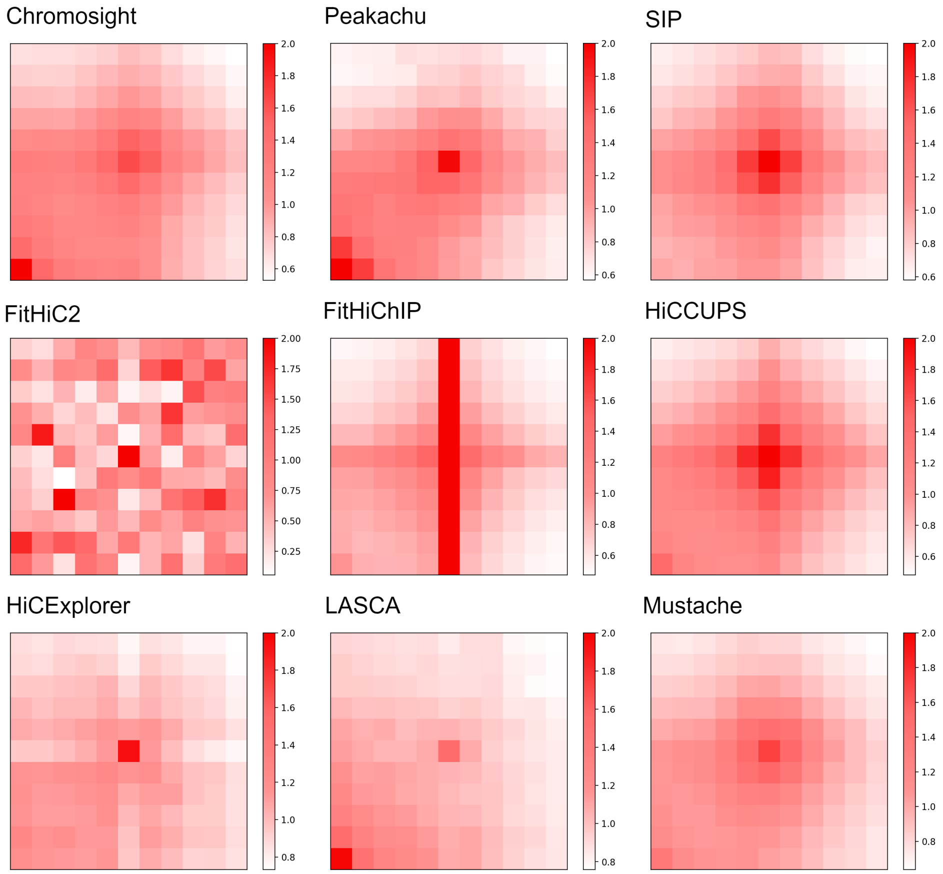


**Figure 21.** APA plot for chromosome 6 at 5KB using GM12878 KR normalized data. FitHiChIP (4.86) has a vertical strong color through the center and FitHiC2 shows asymmetric color with a high APA score.

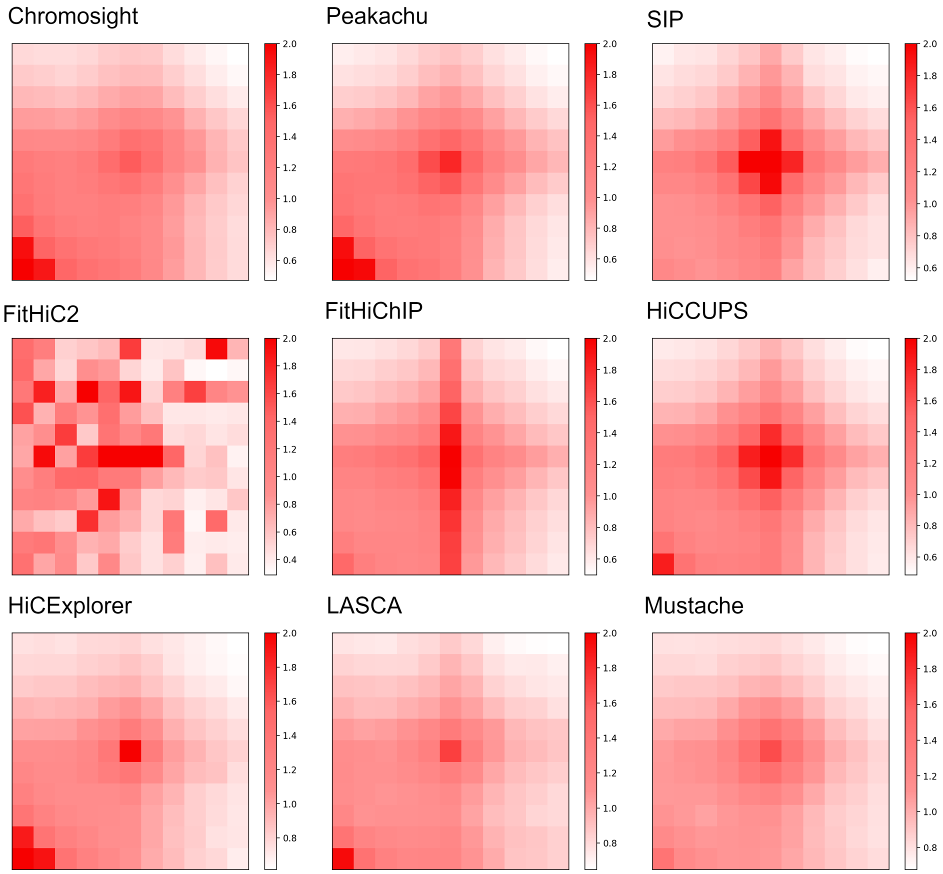


**Figure 22.** APA plot for chromosome 6 at 10KB using GM12878 KR normalized data. FitHiChIP shows the similar enrichment pattern as 5KB but a lower APA score. Chromosight, Peakachu, and HiCExplorer shows a similar color contrast at the lower left corner, and Chromosight scored less than 1.

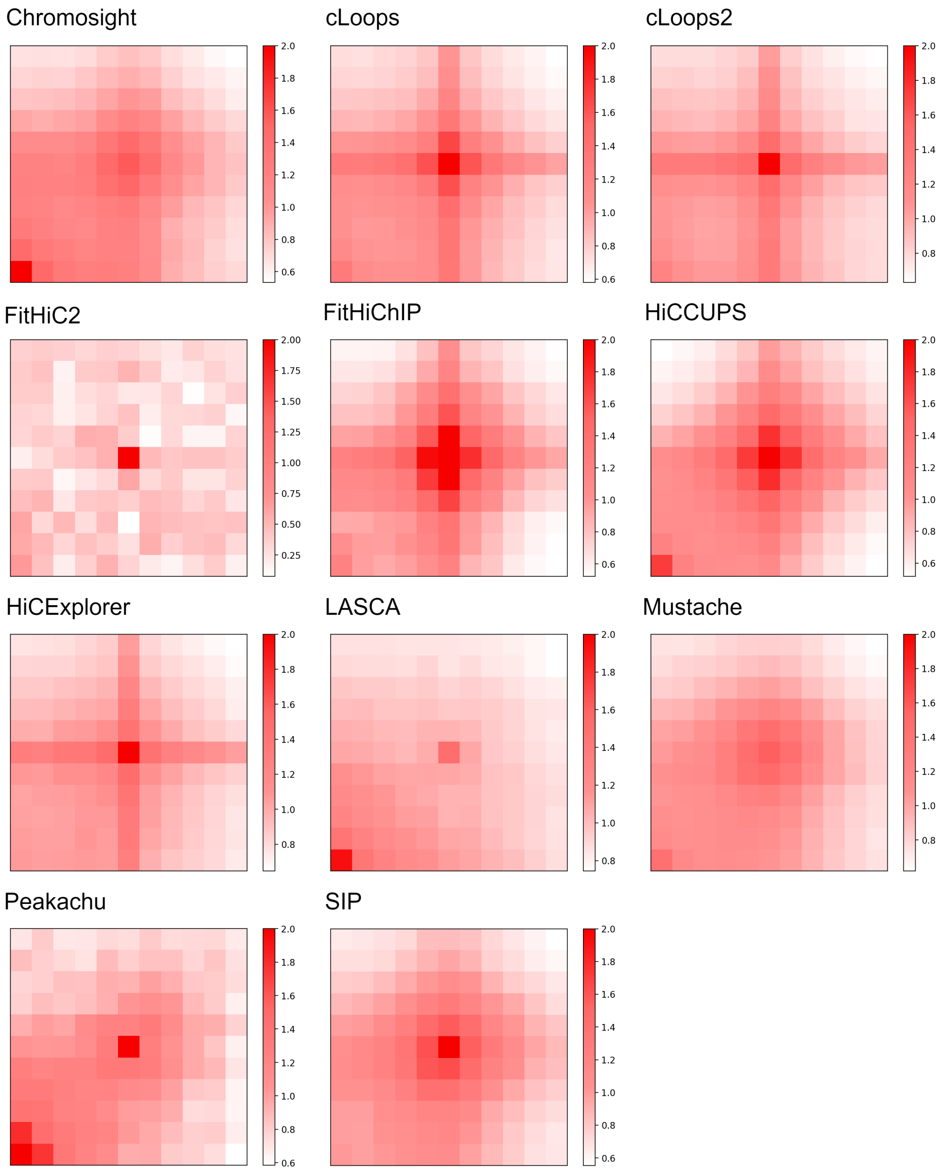


**Figure 23.** APA plot for chromosome 1 at 5KB using GM12878 replicate data. Peakachu, SIP, cLoops, cLoops2, HiCExplorer, FitHiChIP, HiCCUPS, FitHiC2, and LASCA have strong focal point in the center. Except FitHiC2 with the highest APA score, all of them have comparable enrichment around the center.

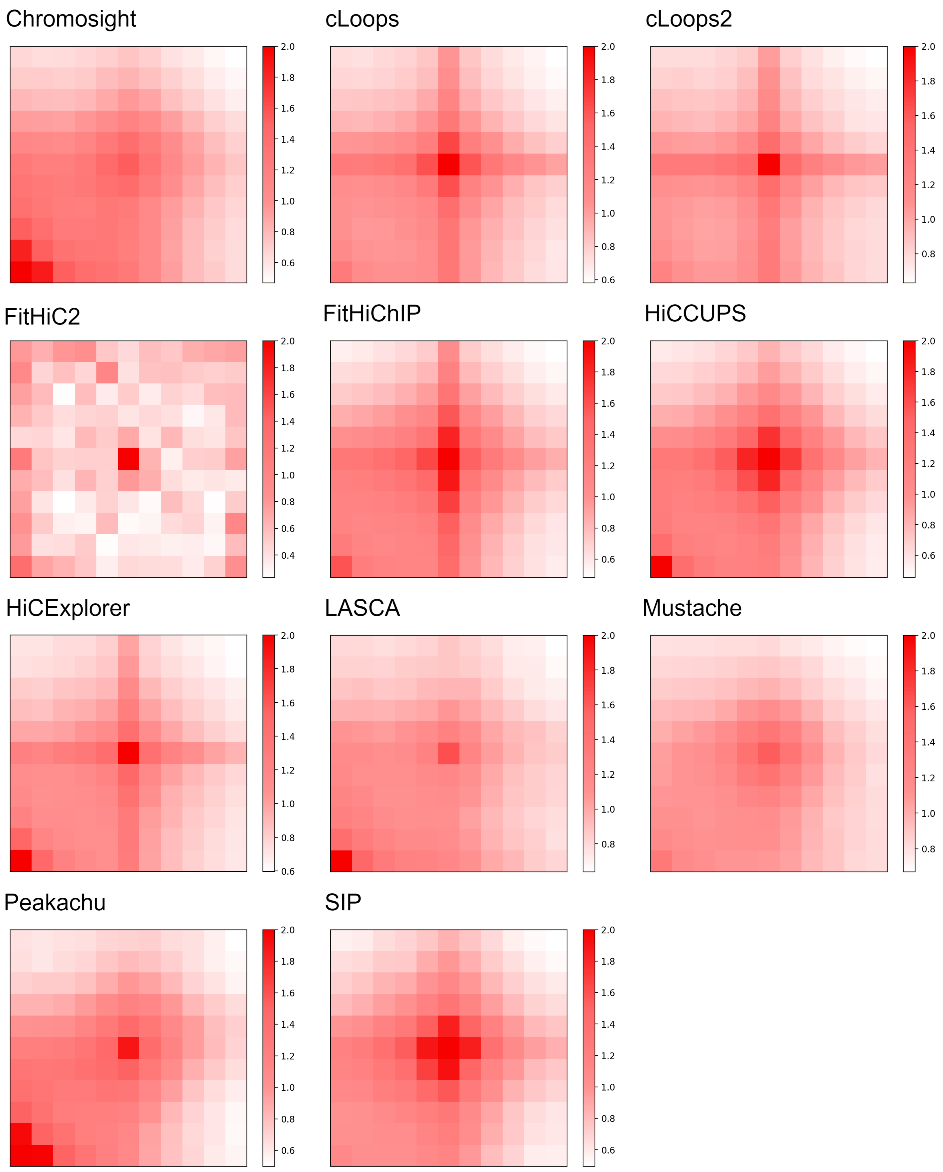


**Figure 24.** APA plot for chromosome 1 at 10KB using GM12878 replicate data. SIP and HiCCUPS show almost similar plot. Chromosight, HiCCUPS, HiCExplorer, LASCA, and Peakachu shows enrichment at the lower left corner with the center where Chromosight APA score is less than 1.

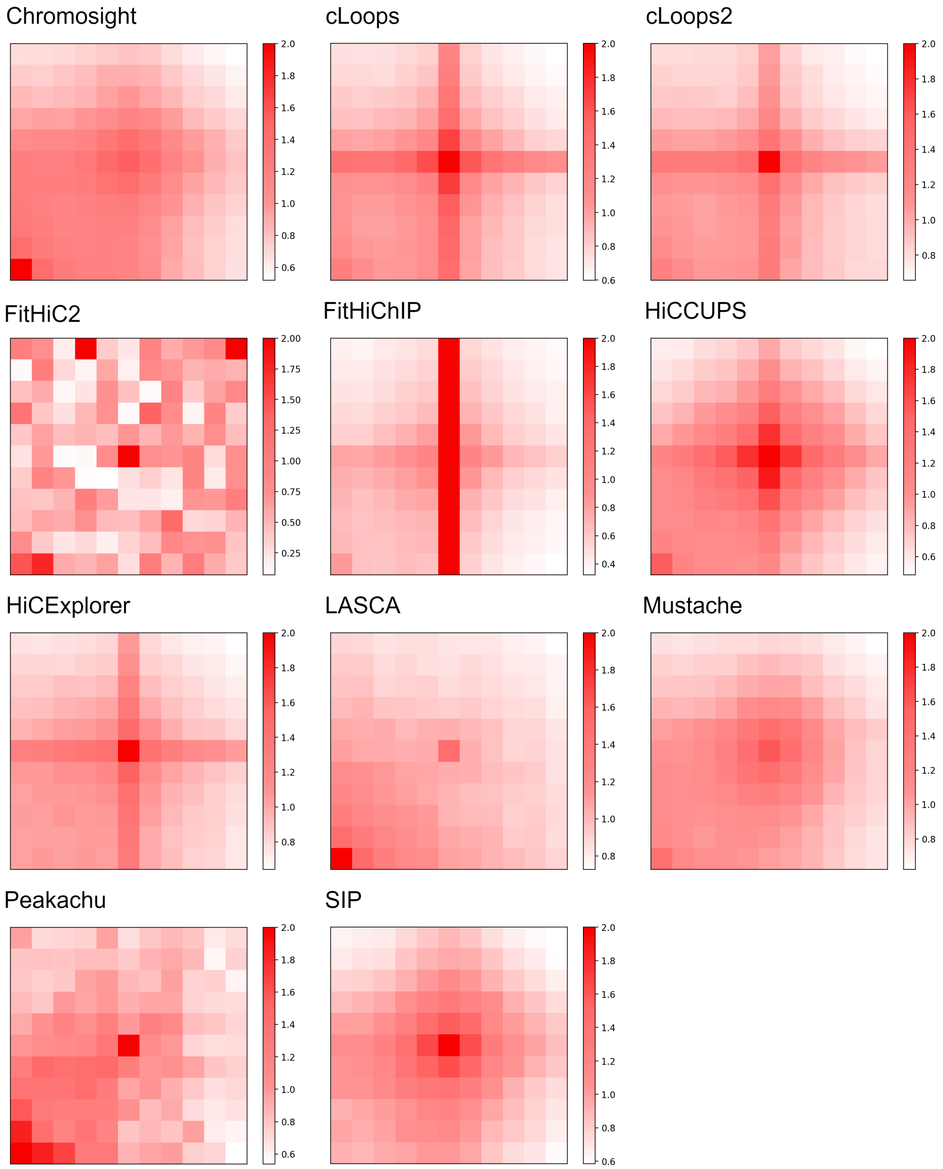


**Figure 25.** APA plot for chromosome 6 at 5KB using GM12878 replicate data. FitHiChIP has vertically strongest color through the center. FitHiC2 (67.8) and Peakachu (1.28) have almost similar pattern in their plots.

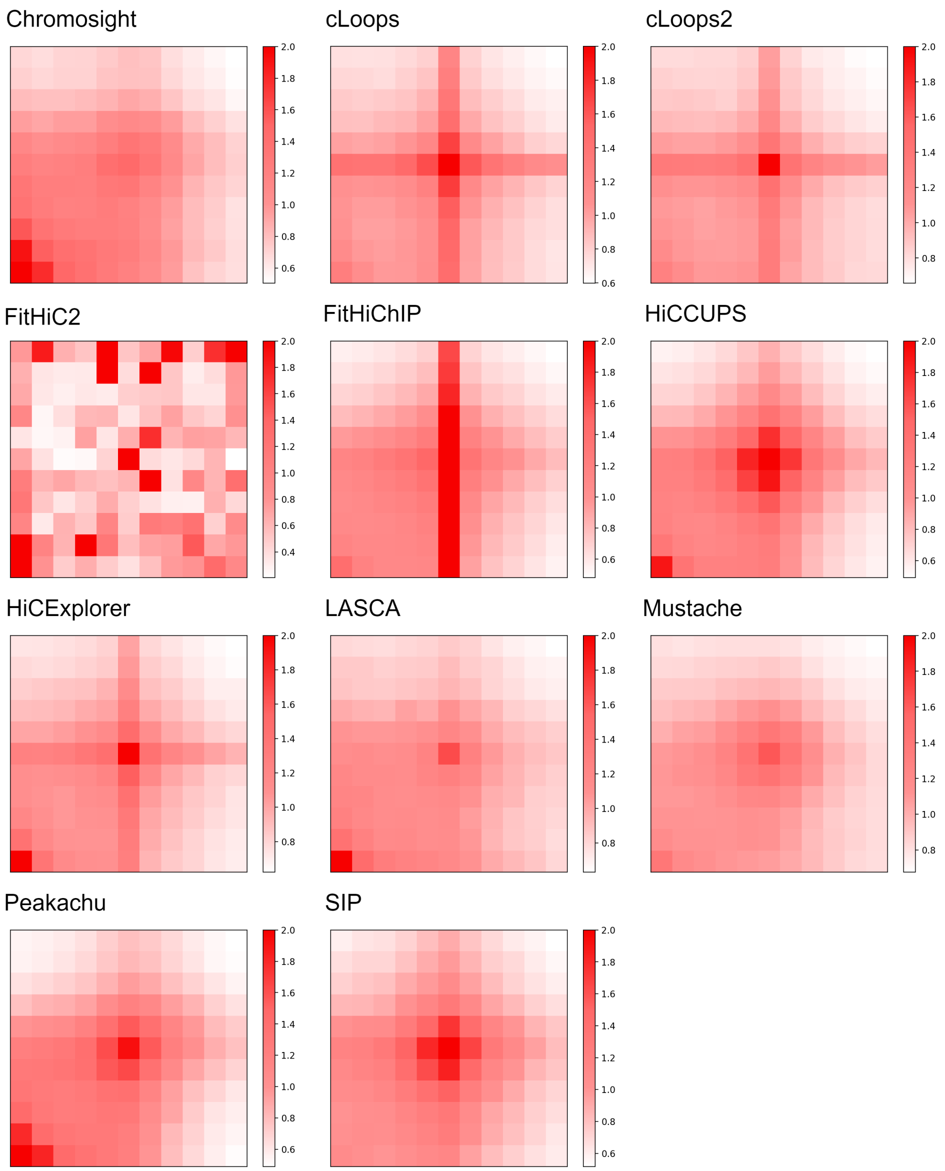


**Figure 26.** APA plot for chromosome 6 at 10KB using GM12878 replicate data. FitHiChIP continued the pattern from 5KB data and Peakachu has smoother heatmap compared to 5KB data. FitHiC2 scores 23.4 which is the highest and Chromosight (0.849) the lowest.

#### 3.5 Recovery Analysis

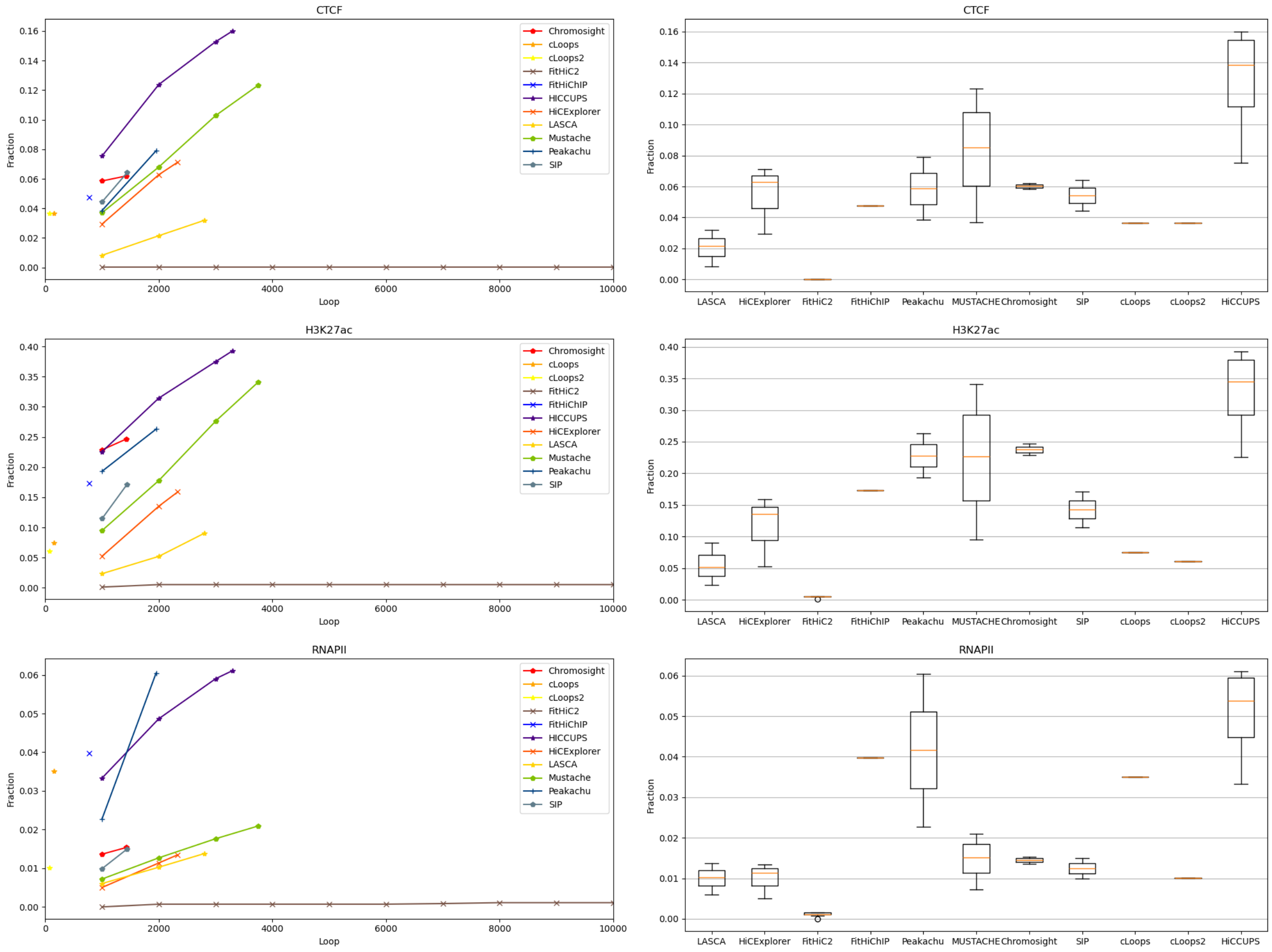


**Figure 27.** Loop recovery fraction of CTCF, H3K27ac, and RNAPII for chromosome 1 at 5KB using primary GM12878. FitHiC2 recovers a constant amount of CTCF, H3K27ac, and RNAPII loops and this is the lowest amount. Peakachu and HiCCUPs has the highest amount of recovery rate for RNAPII. For CTCT and H3K27ac, HiCCUPS holds the highest fraction of amount. On average, Mustache recovers a significant amount of loops. FitHiC2 shows an outlier data at the lower portion for H3K27ac3 and RNAPII.

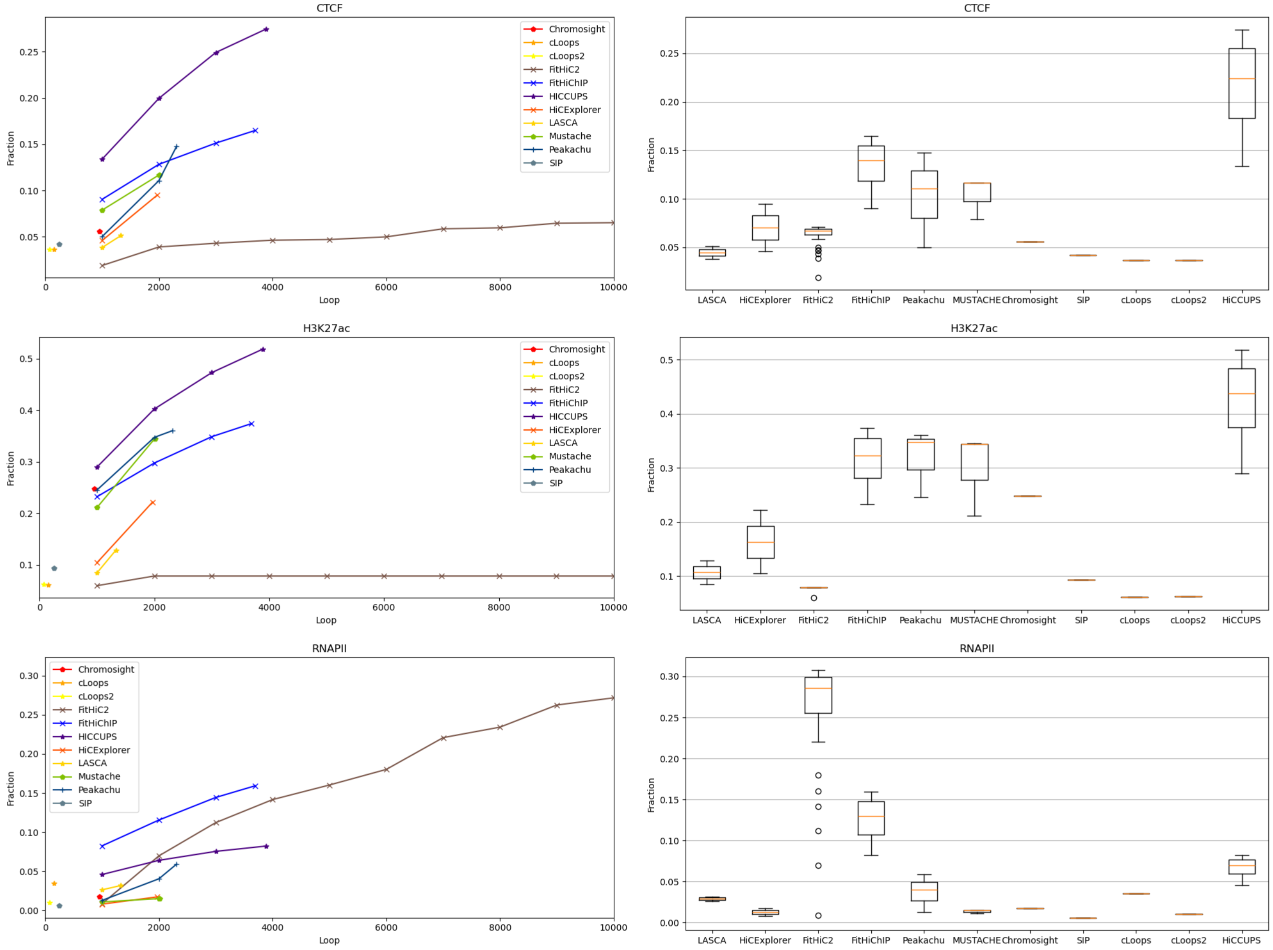


**Figure 28.** Loop recovery fraction of CTCF, H3K27ac, and RNAPII for chromosome 1 at 10KB using primary GM12878. Though FitHiC2 has a wide range of outliers, it shows an improvement in recovery rate for CTCF and RNAPII, and recovers the highest amount of RNAPII loops compared to other tools. HiCCUPS recovers most amount of CTCF and H3K27ac loops.

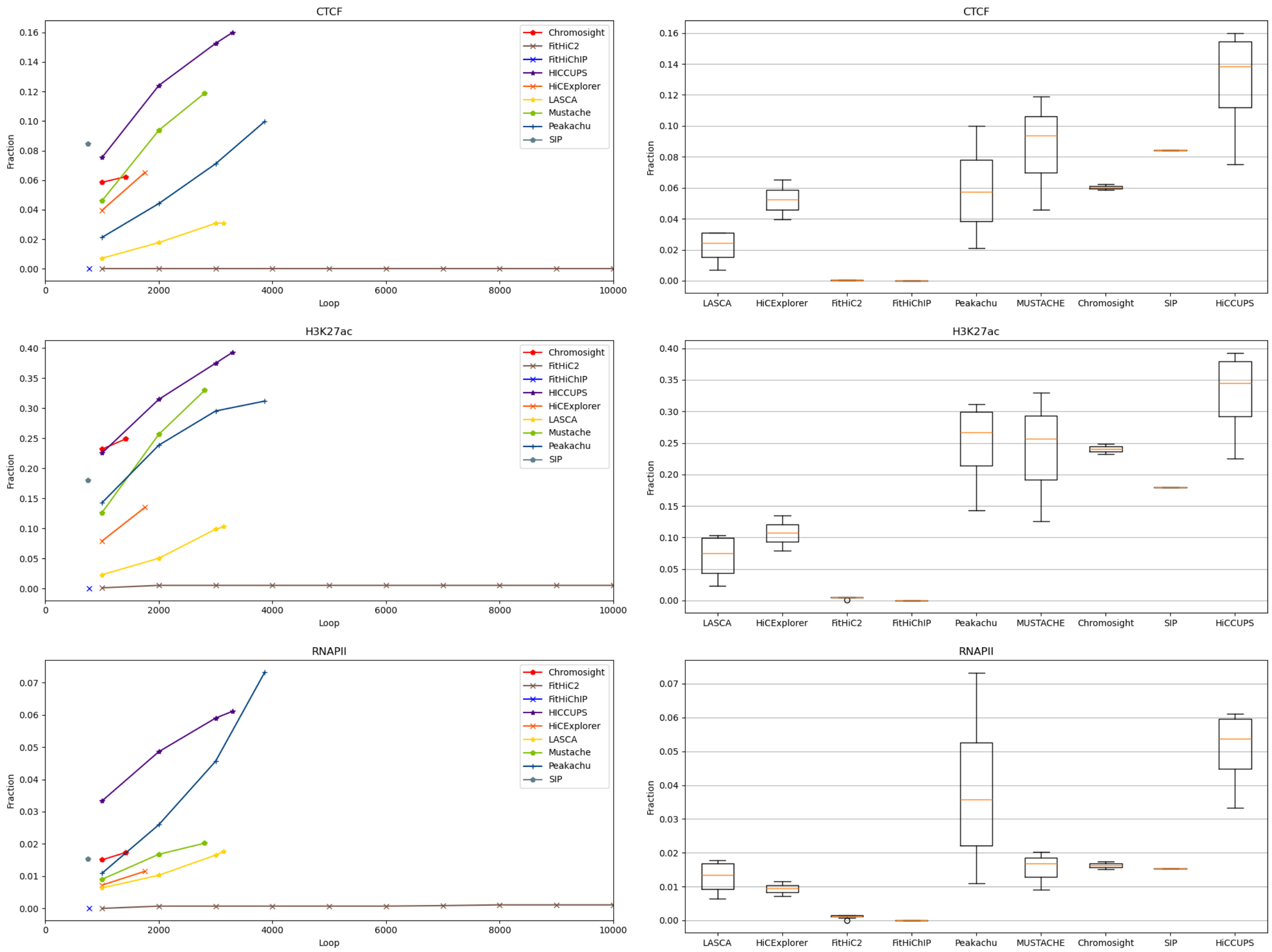


**Figure 29.** Loop recovery fraction of CTCF, H3K27ac, and RNAPII for chromosome 1 at 5KB using KR normalized GM12878. HiCCUPS recovers highest number of fraction CTCF and H3K27ac loops, and Peakachu recovers RNAPII loops. FitHiC2 remain constant which is almost near to 0. On average, Mustache recovers a good number of loops in these three category. FitHiC2 has outliers in H3K27ac and RNAPII.

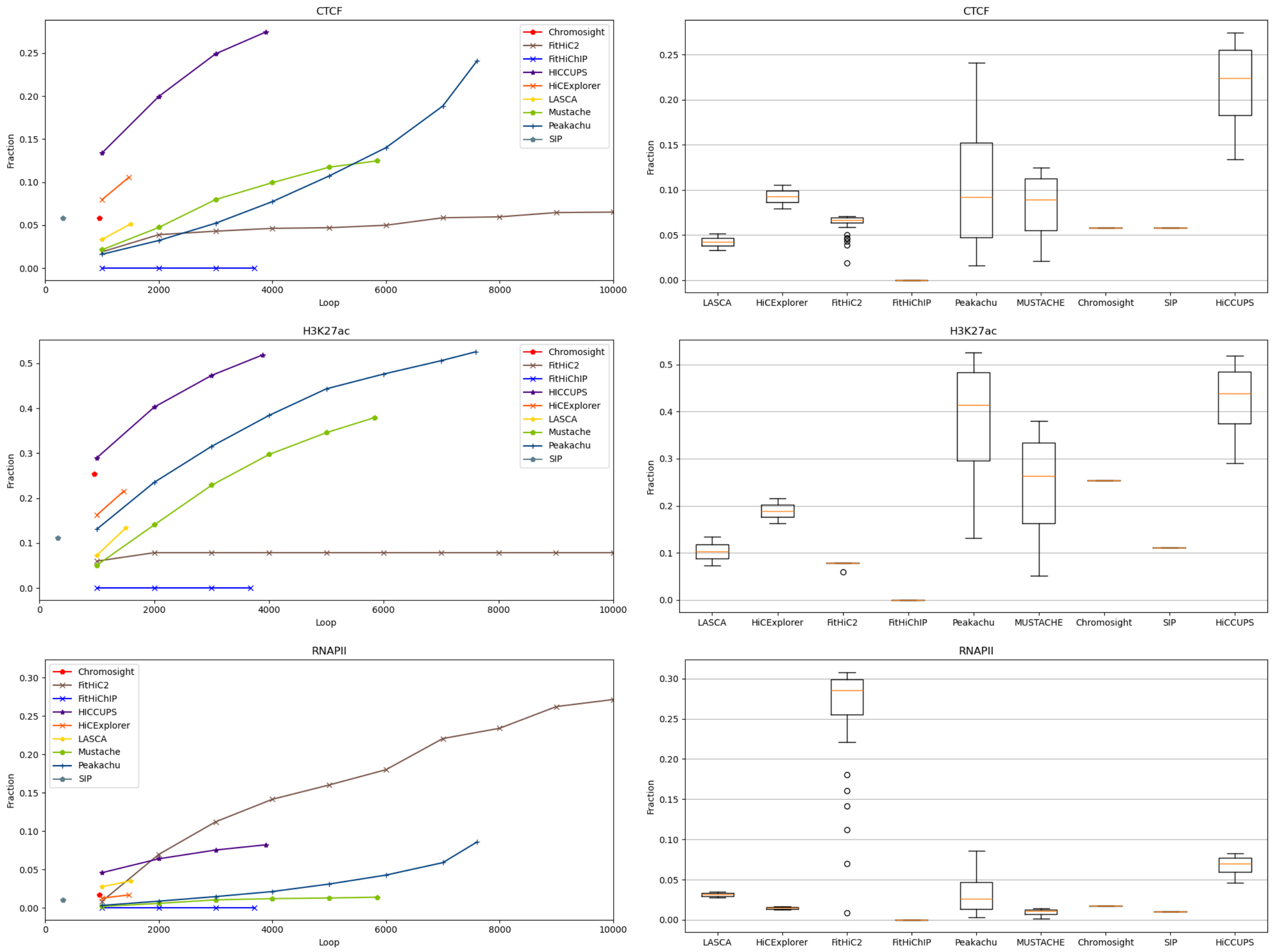


**Figure 30.** Loop recovery fraction of CTCF, H3K27ac, and RNAPII for chromosome 1 at 10KB using KR normalized GM12878. HiCCUPS and Peakachu recovers significant amount of CTCF and H3K27ac loops and Mustache recovers an average number of fraction. In contrast, RNAPII loops are mostly recovered by FitHiC2 with outliers in the three cases.

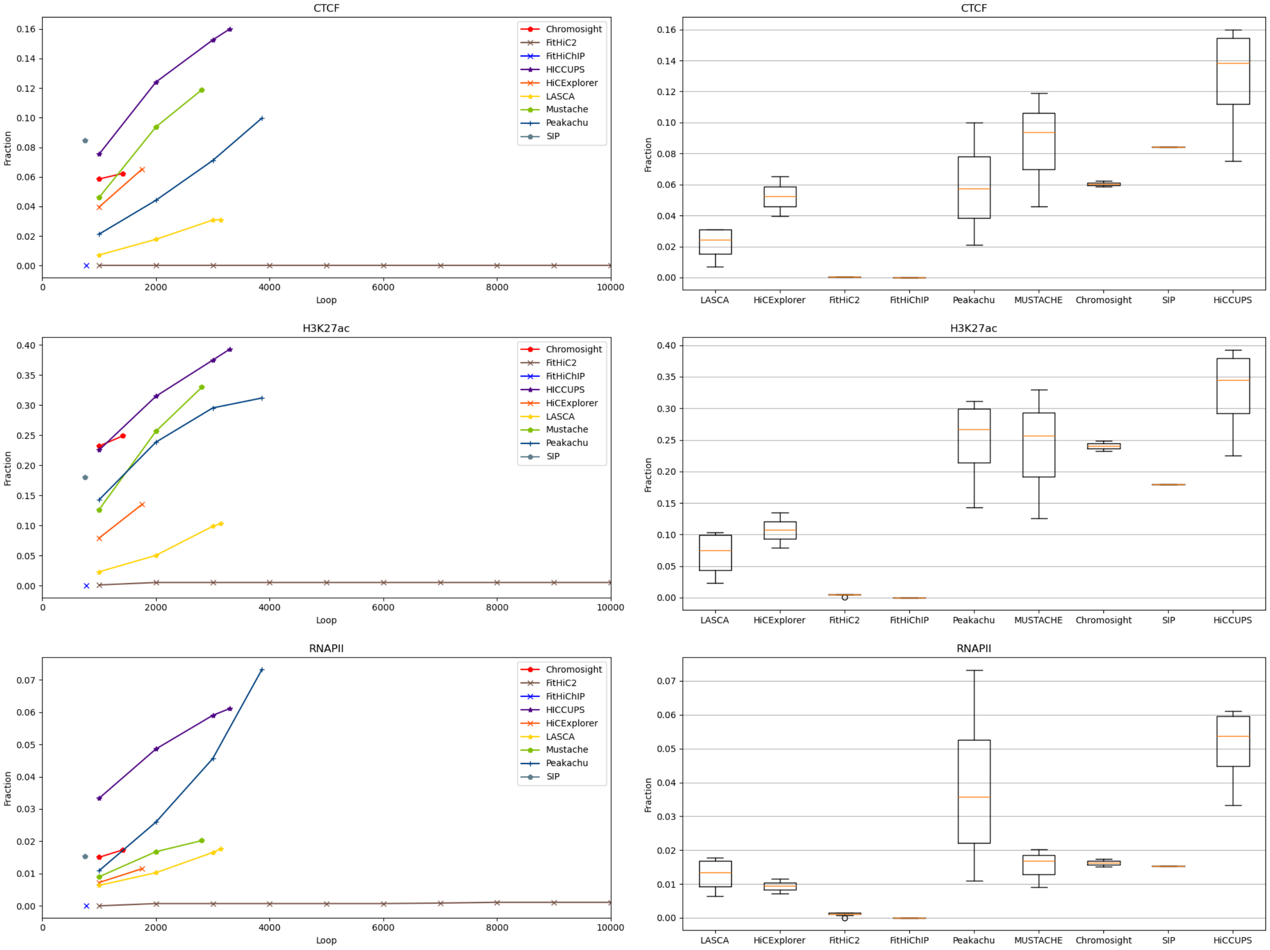


**Figure 31.** Loop recovery fraction of CTCF, H3K27ac, and RNAPII for chromosome 6 at 5KB using KR normalized GM12878. HiCCUPS and Mustache recovers shear amount of CTCF and H3K27ac loops compared to other tools. Peakachu and HiCCUPS recovers highest fraction of RNAPII loops. FitHiC2 shows a constant recovery rate with some outliers.

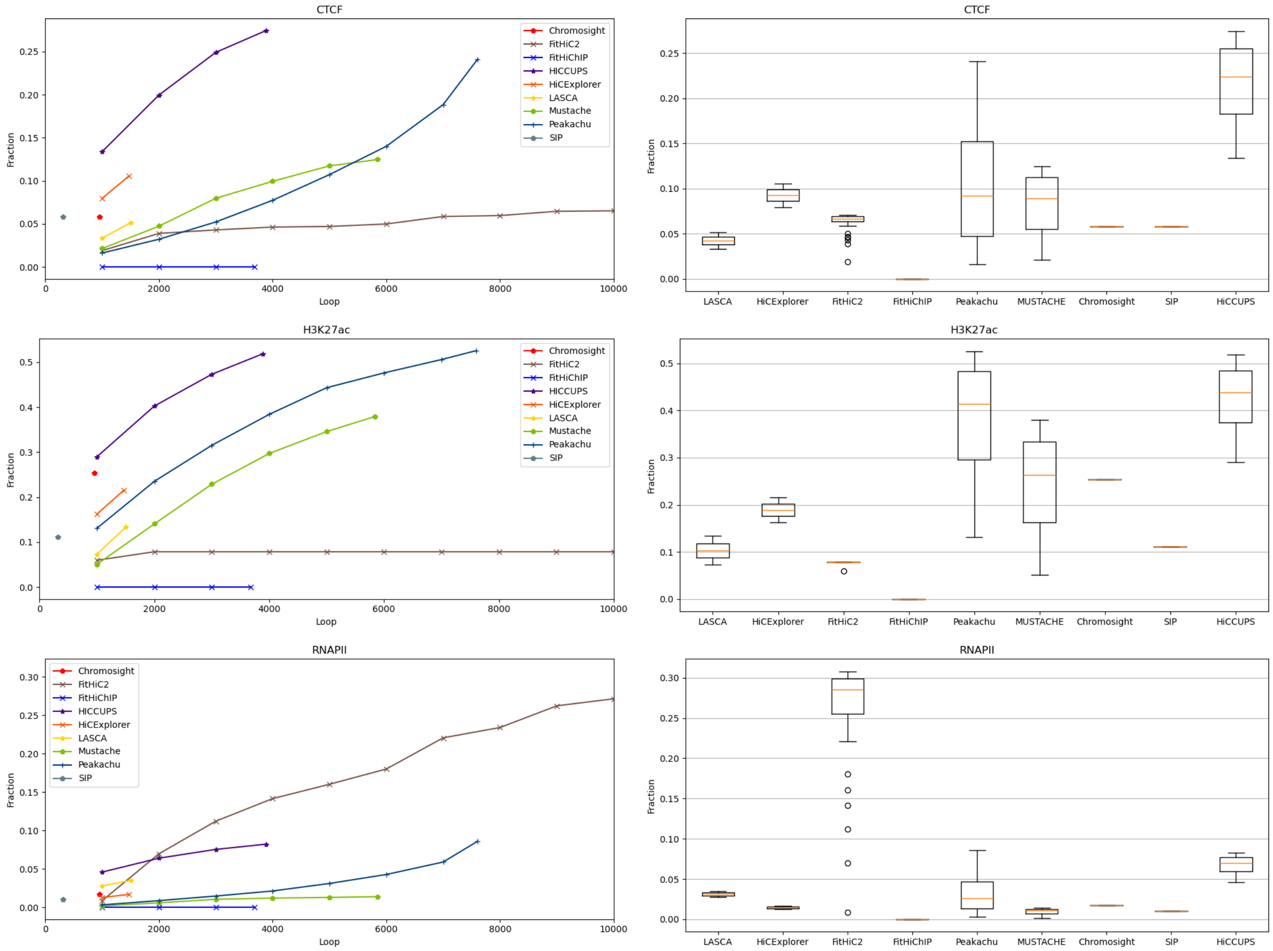


**Figure 32.** Loop recovery fraction of CTCF, H3K27ac, and RNAPII for chromosome 6 at 10KB using KR normalized GM12878. A large amount of CTCF and H3K27ac loops are recovered by HiCCUPS and Peakachu whereas Mustache recover an average number of loops among the other tools. FitHiChIP recovers near 0 loops. FitHiC2 recovers above 25% RNAPII loops and it has outliers in the three cases.

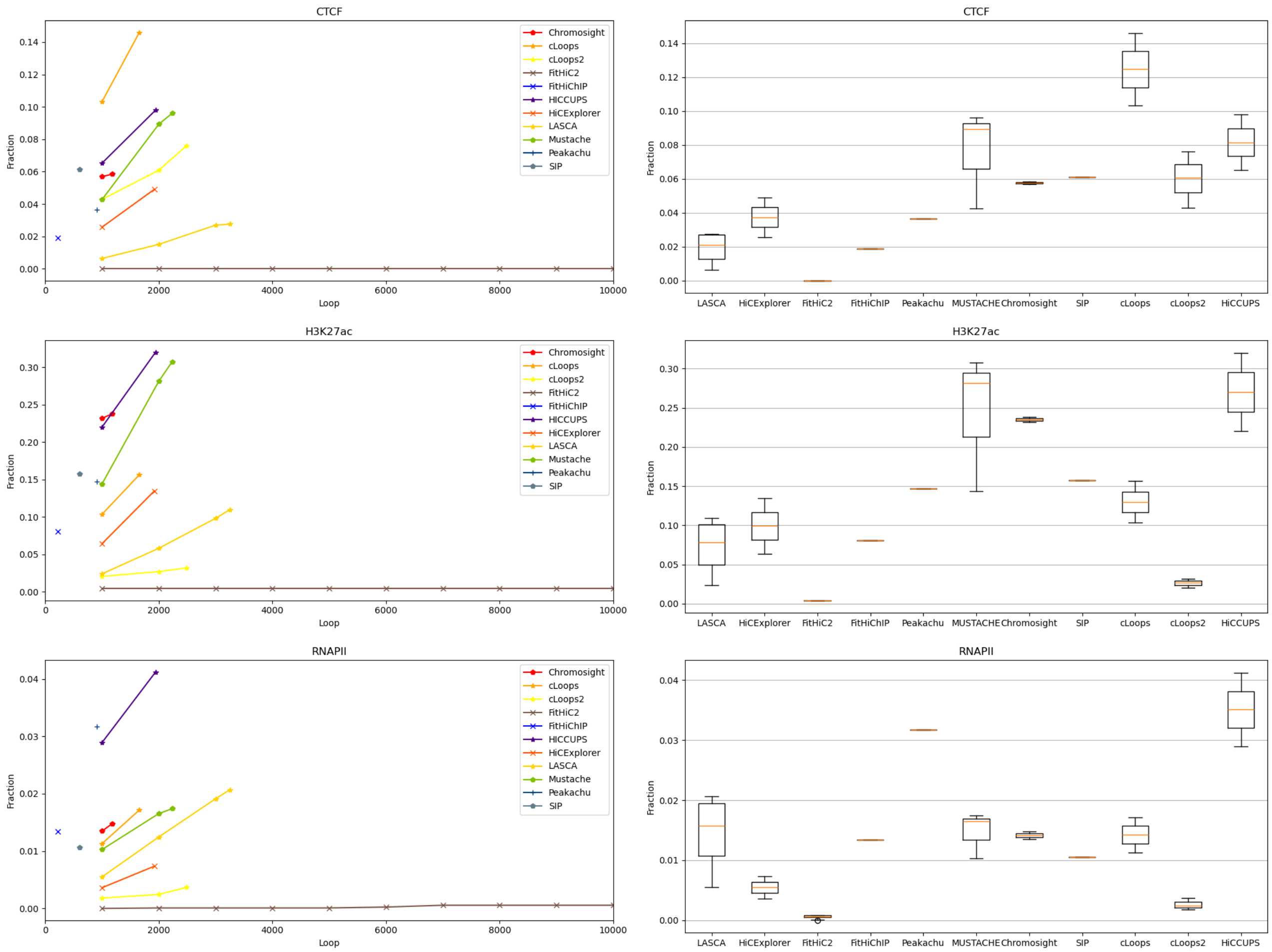


**Figure 33.** Loop recovery fraction of CTCF, H3K27ac, and RNAPII for chromosome 1 at 5KB using replicate GM12878. cLoops recovers a shear amount of CTCF loops, Mustache and HiCCUPS recovers near about 10% CTCF loops. Mustache and HiCCUPS recovers above 0.30 fraction of H3K27ac loops. Chromosight recovers near about 0.25 fraction of H3K27ac loops. RNAPII loops mostly rocovered by HiCCUPS. FitHiC2 shows a outlier point for RNAPII recovery.

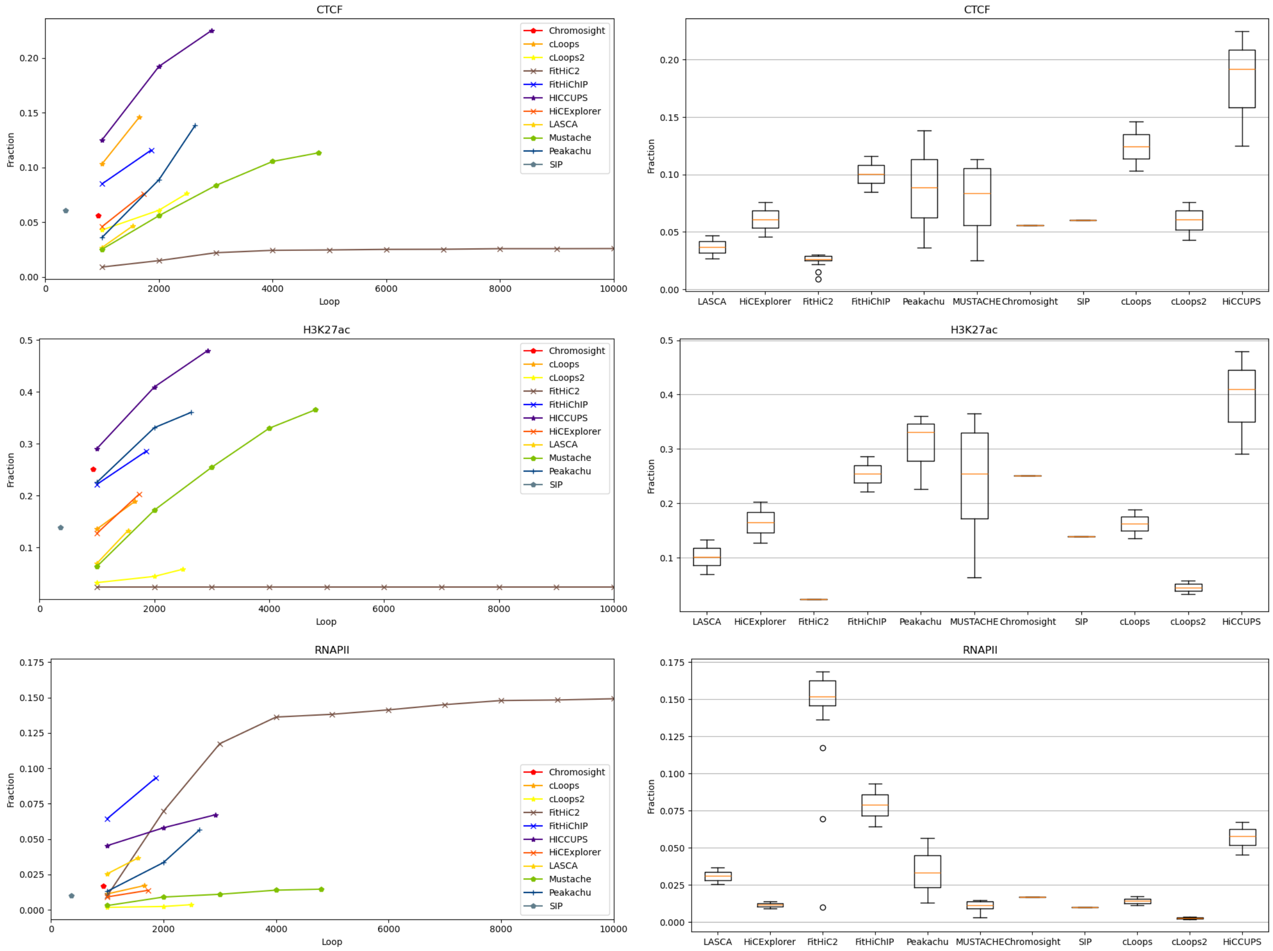


**Figure 34.** Loop recovery fraction of CTCF, H3K27ac, and RNAPII for chromosome 1 at 10KB using replicate GM12878. HiCCUPS and Peakachu recovers a large fractional amount of CTCF and H3K27ac loops and FitHiC2 recovers most RNAPII loops compared to other tools and has outlier points for CTCF and RNAPII recovery. Mustache holds the middle position for CTCF and H3K27ac, and HiCCUPS and FitHiChIP for RNAPII.

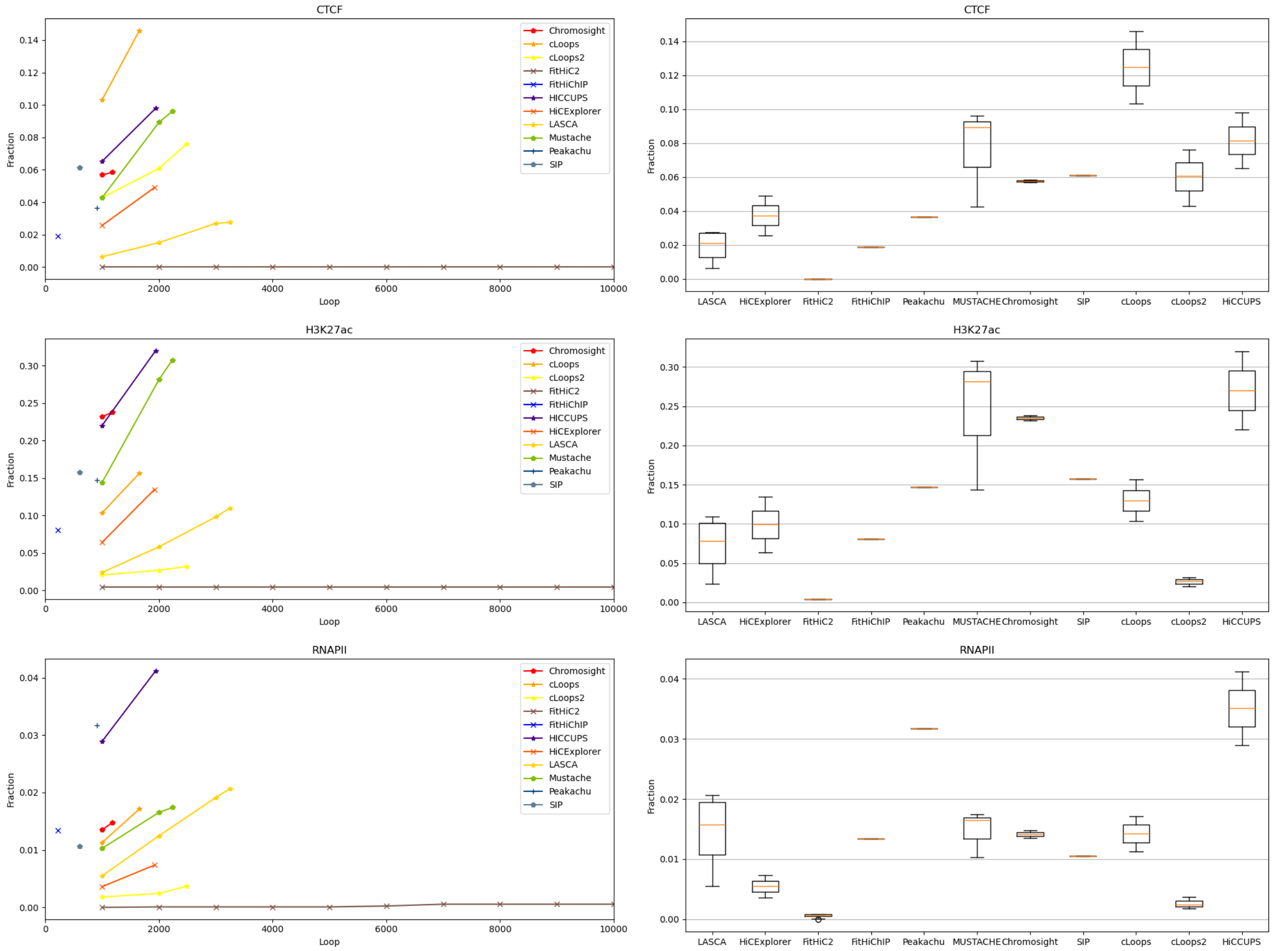


**Figure 35.** Loop recovery fraction of CTCF, H3K27ac, and RNAPII for chromosome 6 at 5KB using replicate GM12878. cLoops recovers above 0.14 fractional CTCF loops, HiCCUPS and Mustache recovers above 0.30 H3K27ac loops. HiCCUPS recovers above 0.04 fractional RNAPII loops. LASCA recovers average in terms of CTCF and RNAPII loops recovery. FitHiC2 shows an outlier point in RNAPII recovery.

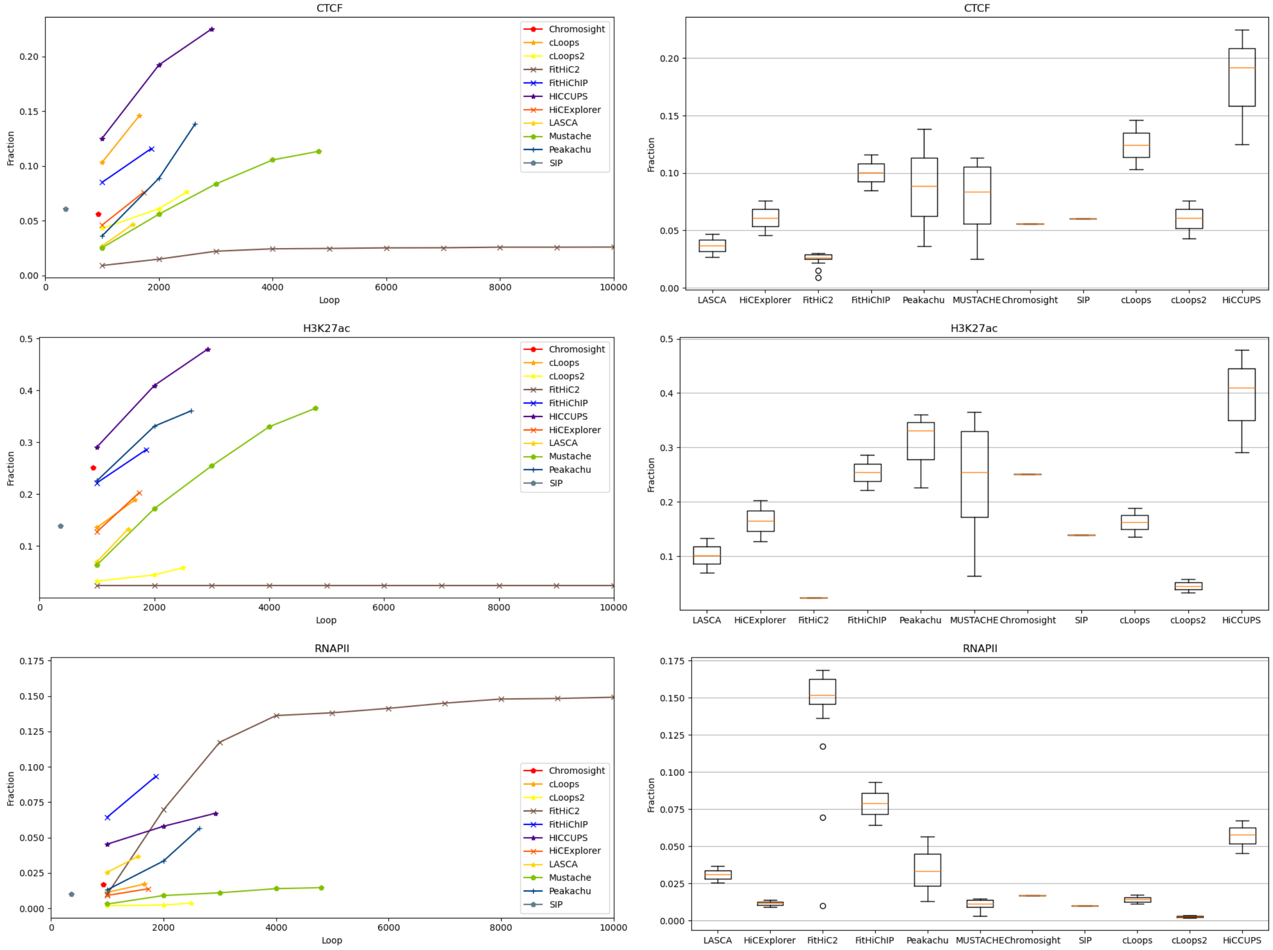


**Figure 36.** Loop recovery fraction of CTCF, H3K27ac, and RNAPII for chromosome 6 at 10KB using replicate GM12878. HiCCUPS recovers most of the CTCF and H3K27ac loops whereas FitHiC2 recovers RNAPII loops and has outlier points in CTCF and RNAPII recovery. cLoops2 recovers least fraction of H3K27ac and RNAPII loops.

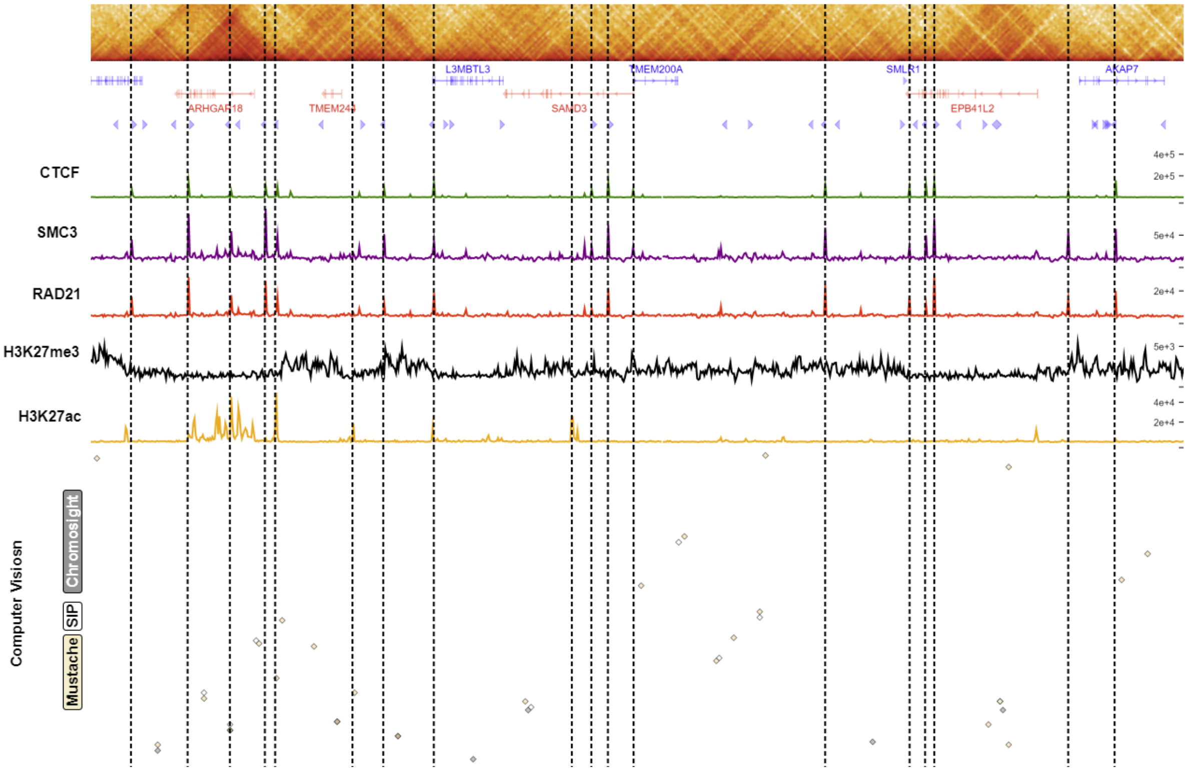


**Figure 37.** Comparison of computer vision based tools (Mustache, SIP and Chromosight) in a random region (129.7M-131.6M and 62.4M - 62.5M) for chromosome 6 using GM12878 cell line. We depicted gene annotation, CTCT motif orientation, ChIP signals for CTCF, SMC3, RAD21, H3K27me3 and H3K27ac below the contact map (Plotted using HiGlass). At the bottom, loops from three tools show their regions with biologically significant area.

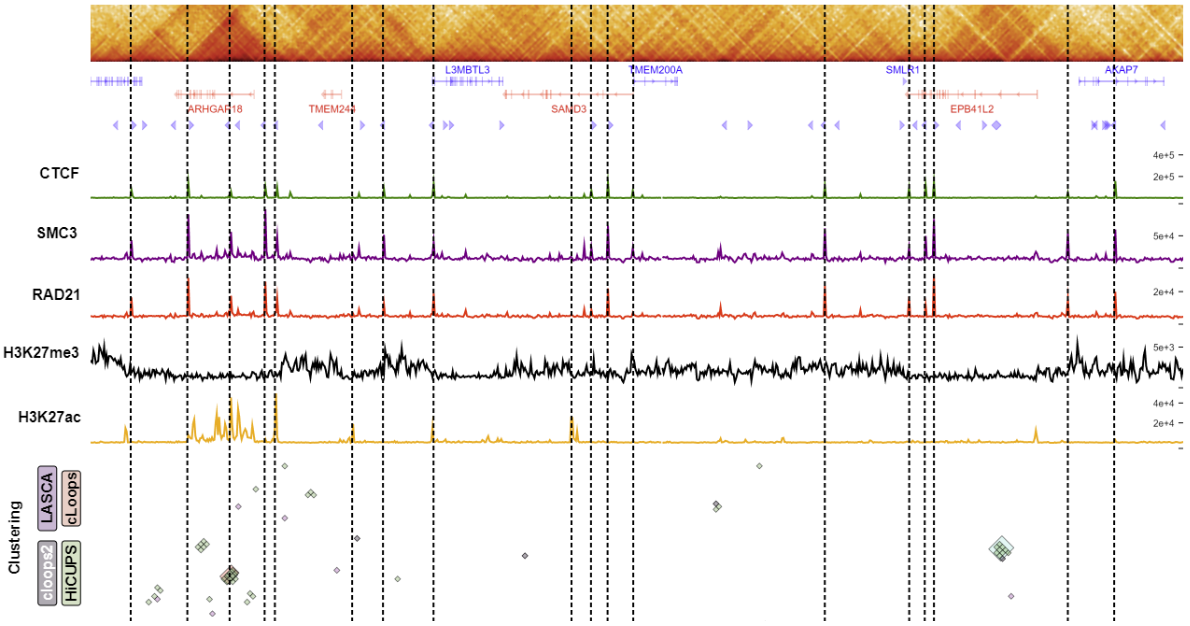


**Figure 38.** Comparison of clustering based tools (HiCCUPS, cLoops2, cLoops and LASCA) in a random region (129.7M-131.6M and 62.4M - 62.5M) for chromosome 6 using GM12878 cell line. We depicted gene annotation, CTCT motif orientation, ChIP signals for CTCF, SMC3, RAD21, H3K27me3 and H3K27ac below the contact map (Plotted using HiGlass). At the bottom, loops from HiCCUPS, cLoops2, cLoops and LASCA show their regions with biologically significant area.

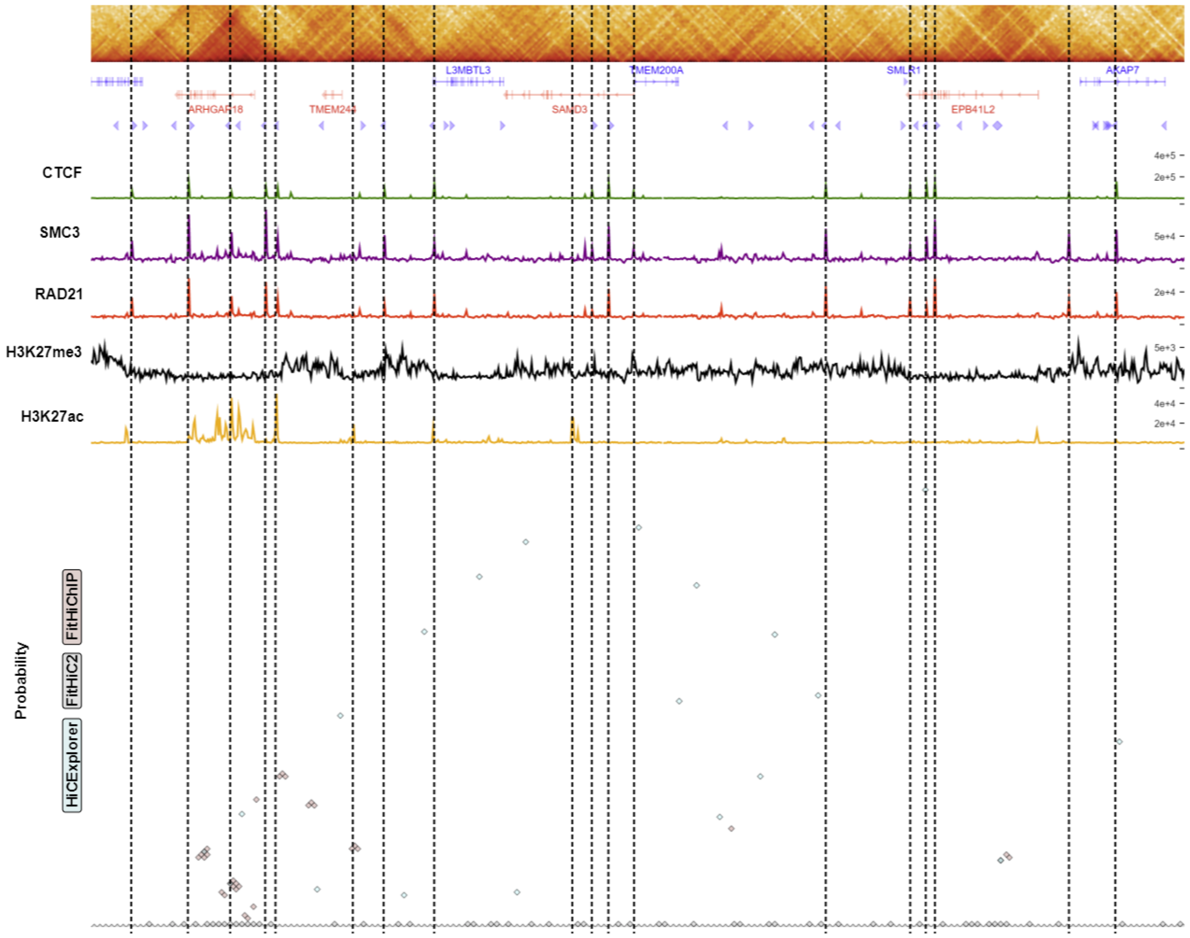


**Figure 39.** Comparison of probability based tools (HiCExplorer, FitHiC2, and FitHiChIP) in a random region (129.7M-131.6M and 62.4M - 62.5M) for chromosome 6 using GM12878 cell line. We depicted gene annotation, CTCT motif orientation, ChIP signals for CTCF, SMC3, RAD21, H3K27me3 and H3K27ac below the contact map (Plotted using HiGlass). At the bottom, loops from three tools show their regions with biologically significant area.

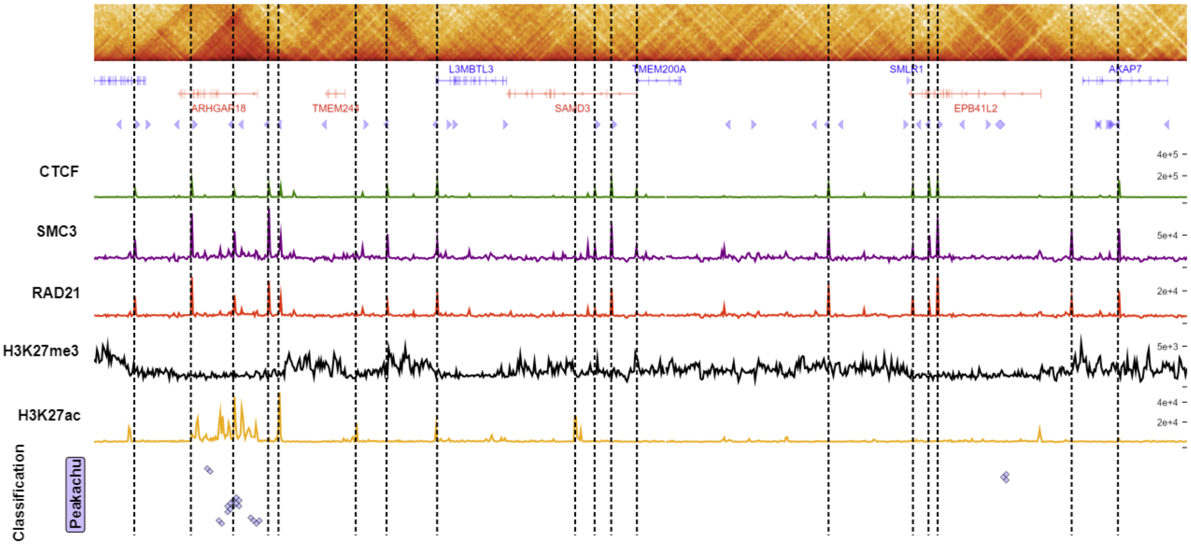


**Figure 40.** Comparison of classification based tools (Peakachu) in a random region (129.7M-131.6M and 62.4M - 62.5M) for chromosome 6 using GM12878 cell line. We depicted gene annotation, CTCT motif orientation, ChIP signals for CTCF, SMC3, RAD21, H3K27me3 and H3K27ac below the contact map (Plotted using HiGlass). At the bottom, loops from Peakachu shows their regions with biologically significant areas.
